# Population-level whole genome sequencing of *Ascochyta rabiei* identifies genomic loci associated with isolate aggressiveness

**DOI:** 10.1101/2024.04.02.587819

**Authors:** Niloofar Vaghefi, Ido Bar, Jonathan Wanderley Lawley, Prabhakaran Thanjavur Sambasivam, Melody Christie, Rebecca Ford

## Abstract

Ascochyta blight caused by the ascomycete *Ascochyta rabiei* poses a major biotic threat to chickpea (*Cicer arietinum*) industries worldwide and incurs substantial costs to the Australian multimillion-dollar chickpea industry both in disease control and yield loss. The fungus was introduced to Australia in the 1970s from an unknown source population, and within a few decades, successfully established in all Australian agroecological chickpea growing regions. Although genetically highly clonal, a broad range of phenotypic variation in terms of aggressiveness exists among the Australian *A. rabiei* isolates. More recently, highly aggressive isolates capable of causing severe disease symptoms on moderate to highly resistant chickpea cultivars have increased in frequency. To identify genetic loci potentially associated with *A. rabiei* aggressiveness on Australian chickpea cultivars, we performed deep genome sequencing of 230 isolates collected from a range of agroecological chickpea growing regions between 2013 and 2020. Population genetic analyses using genome-wide single nucleotide polymorphism data identified three main clusters of genetically closely related isolates in Australia. Phylogenetic analyses showed that highly aggressive phenotypes developed multiple times independently throughout the phylogeny. Results point to minor contribution of multiple genetic regions and most likely epigenomic variations to aggressiveness of *A. rabiei* isolates on Australian chickpea cultivars.

**IMPACT STATEMENT:** This research introduces new knowledge on the Australian *A. rabiei* population structure, molecular pathogenicity drivers and evolution as a clonal pathogen through comprehensive whole-genome sequencing approach. The knowledge generated on the structure and origin of the Australian *A. rabiei* and existence of only one mating type continues to inform researchers, growers, breeders and the broader industry on the importance for continued tight biosecurity measures and inform development of accurate and informed disease management and resistance breeding strategies. This research provides a rare real-life example to the effect of genetic drift on a clonal pathogen population and the importance of biosecurity to protect introduction from non-endemic isolates through seed importation in the current era of international markets.

**DATA SUMMARY:** An online dataset containing the data and code required to reproduce the results found in this publication have been deposited at Zenodo (DOI 10.5281/zenodo.12575659). Isolate aggressiveness and collection metadata are available in the Ascochyta dashboard at http://bit.ly/asco-dashboard. Raw sequencing data used in this study was deposited to the NCBI Short Read Archive (SRA) and is available through BioProject PRJNA1175002. The authors confirm all supporting data, code and protocols have been provided within the article or through supplementary data files.

## INTRODUCTION

Diseases caused by invasive agricultural pathogens, and specifically necrotrophic fungi, present a massive threat to food production industries. Of these, Ascochyta Blight caused by *Ascochyta rabiei* poses a major biotic threat to chickpea (*Cicer arietinum*) worldwide and costs the Australian multimillion-dollar chickpea industry an estimated $34.9 million in disease control expenses and $4.8 million in yield losses (1). The fungus was introduced to Australia in the 1970s from an unknown source population and, within a few decades, successfully established in all Australian agroecological chickpea growing regions where management is heavily reliant on a combination of fungicide application and host resistance (2–4).

Meanwhile, increases in aggressiveness within *A. rabiei* populations has resulted in breakdown of resistance in many chickpea varieties, limiting the efficacy of their host resistance (3–5). Within Australia, extensive annual *A. rabiei* sampling from all major chickpea growing regions and subsequent phenotypic assessment has revealed temporal increases in population aggressiveness (6), measured as increases in quantitative pathogenicity, *i.e.*, the severity of symptoms caused by the pathogen, or pathogenic fitness (7,8). This is driving the ongoing search for resistance sources within wider germplasm collections (9,10) and the exploration into the genetic factors and genomic regions underpinning the resistance mechanisms (11).

Although highly clonal, a broad range in aggressiveness exists among the Australian isolates (3,4,12,13) and highly aggressive isolates, able to cause severe disease on recently released ‘resistant’ chickpea varieties, are increasing in frequency (6,14). However, the biological and genetic factors underlying this apparent rapid evolution of isolate aggressiveness are unknown.

The clonal nature of the Australian *A. rabiei* population contrasts with most other regions of the world, where *A. rabiei* populations are genetically diverse and regularly undergo sexual recombination in a heterothallic manner. This is due to the presence of isolates with alternate mating-type idiomorphs (MAT1-1 and MAT1-2) within the population (15–18). However, after introduction to Australia, *A. rabiei* underwent a severe bottleneck and reduction in genetic diversity (13,19). Moreover, it was forced into asexuality, likely due to the introduction of a single mating-type (MAT1-2; (20)). Intensive infield sampling of diseased crops across chickpea growing regions and screening of *A. rabiei* populations over the past decade has identified only the *MAT1-2* locus in the Australian population (6,13,14). Therefore, all evidence to date point to a genetically homogenous and stable clonal population with low genetic diversity and no signs of recombination in Australia. On the other hand, the increased aggressiveness indicates that the Australian *A. rabiei* population is evolving rapidly, with more frequent emergence of highly aggressive isolates, despite its asexuality and very limited genetic diversity (4).

Asexual organisms with low genetic diversity are expected to have a slow rate of evolution due to reduced genetic variation, accumulation of deleterious mutations, and clonal interference (21). However, many asexual filamentous fungi are able to generate *de novo* genetic variation via various other mechanisms, including parasexuality (a nonsexual mechanism of transferring genetic material without meiosis) (22), chromosomal rearrangements (23), and conditionally dispensable chromosomes (24). However, these mechanisms have not been reported in *A. rabiei*. Rather, the homogenous population structure, high levels of clonality and lack of temporal changes in the Australian *A. rabiei* population are congruent with lack of sexual reproduction and slow generation of genetic variation. The recent emergence of isolates with high aggressiveness on previously resistant chickpea cultivars is, therefore, unexpected, and potentially due to strong selective forces on the standing genetic variation, rather than generation of *de novo* variation in aggressiveness. In any case, significant differences in aggressiveness of isolates that show little genetic variation is interesting and warrants further investigation.

The most recent Australian *A. rabiei* population genomics study used DArTseq^TM^ genotyping to generate a set of single-nucleotide polymorphism (SNP) markers from a reduced representation of the *A. rabiei* genome for the analysis of genetic relatedness among isolates, to assess population structuring and to identify genetic regions associated with aggressiveness (6). The discovery of only 212 polymorphic SNP loci among 180 isolates collected from diverse growing regions in Australia further highlighted the clonality of the Australian *A. rabiei* population and demonstrated the need for a higher marker density to obtain accurate marker-phenotype associations.

In the current study, we hypothesized that the increased frequency of highly aggressive isolates in the Australian *A. rabiei* population is the result of minor genetic modifications which alter the structure or expression of existing effector loci or create novel ones to affect aggressiveness traits. To test this, we conducted deep genome sequencing of 230 *A. rabiei* isolates collected from a range of agroecological zones between 2013 and 2020, to (i) obtain a high-resolution genetic marker set to better characterise the Australian *A. rabiei* population structure, and (ii) identify genetic regions potentially associated with *A. rabiei* aggressiveness in Australia.

## METHODS

### *A. rabiei* isolates, species identification and mating-type screening

A total of 230 Australian *A. rabiei* isolates collected from commercial chickpea fields or naturally infected experimental trial sites from 2013 to 2020 were included in this study (Table S1). Most of the isolates (*n* = 193) were collected in 2020 from chickpea growing regions in 10 agroecological zones with distinct climate and cropping systems (Fig. 1A, Table 1). These were collected using two sampling approaches. First, in fields with high disease incidence, structured, intensive sample collection was conducted in a hierarchical manner, collecting symptomatic chickpea tissue from the four corners and one central location within each field. Second, opportunistic sampling was conducted in the fields where disease incidence was low by haphazardly collecting symptomatic chickpea leaves where disease was found. Fungal isolations from the 2020 samples were conducted according to Christie and Moore (2020) (25) and single-conidium cultures were established on potato dextrose agar (Amyl Media). Additional isolates (*n* = 37) were collected from 2013 to 2017 as reported in earlier studies (6,13,14). Metadata on all the isolates included in this study are provided in Table S1.

**Fig. 1.**
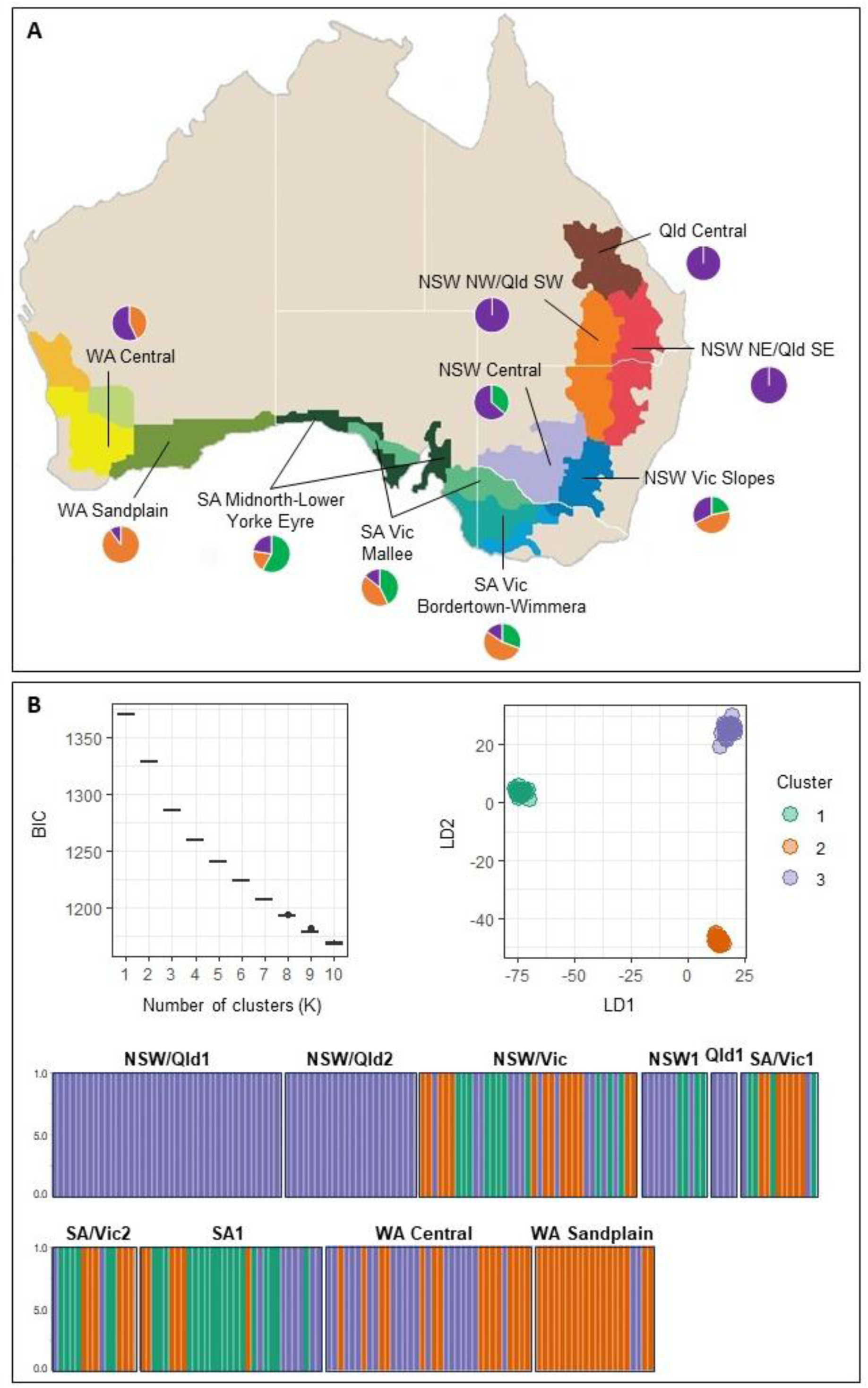
**A.** Sampling locations of 230 *Ascochyta rabiei* isolates sequenced in this study. Geographical regions with different colours represent the Australian agroecological zones with distinct climate and cropping systems, as defined by the Australian Grains Research and Development Corporation (GRDC). Names for the 10 agroecological zones from which *A. rabiei* isolates were collected are provided. Circles depict the frequency of isolates in each agroecological zone from each of the three clusters identified via Discriminant Analysis of Principal Components (DAPC), with green, orange and purple representing cluster 1, 2, and 3, respectively. **B.** DAPC analysis for *A. rabiei* isolates based on 2,554 neutral genome-wide single nucleotide polymorphisms. The graph on the left shows Bayesian Information Criterion (BIC) plotted against the number of inferred clusters. The ordination plot on the right shows the three assigned clusters of genetically related *A. rabiei* isolates. The graph at the bottom shows membership probability of *A. rabiei* isolates to the three genetic clusters detected through DAPC analysis. Each bar on the horizontal axis indicates an individual, the vertical axis shows probability of membership and colors represent membership of individuals in the predicted clusters.

**Table 1.**
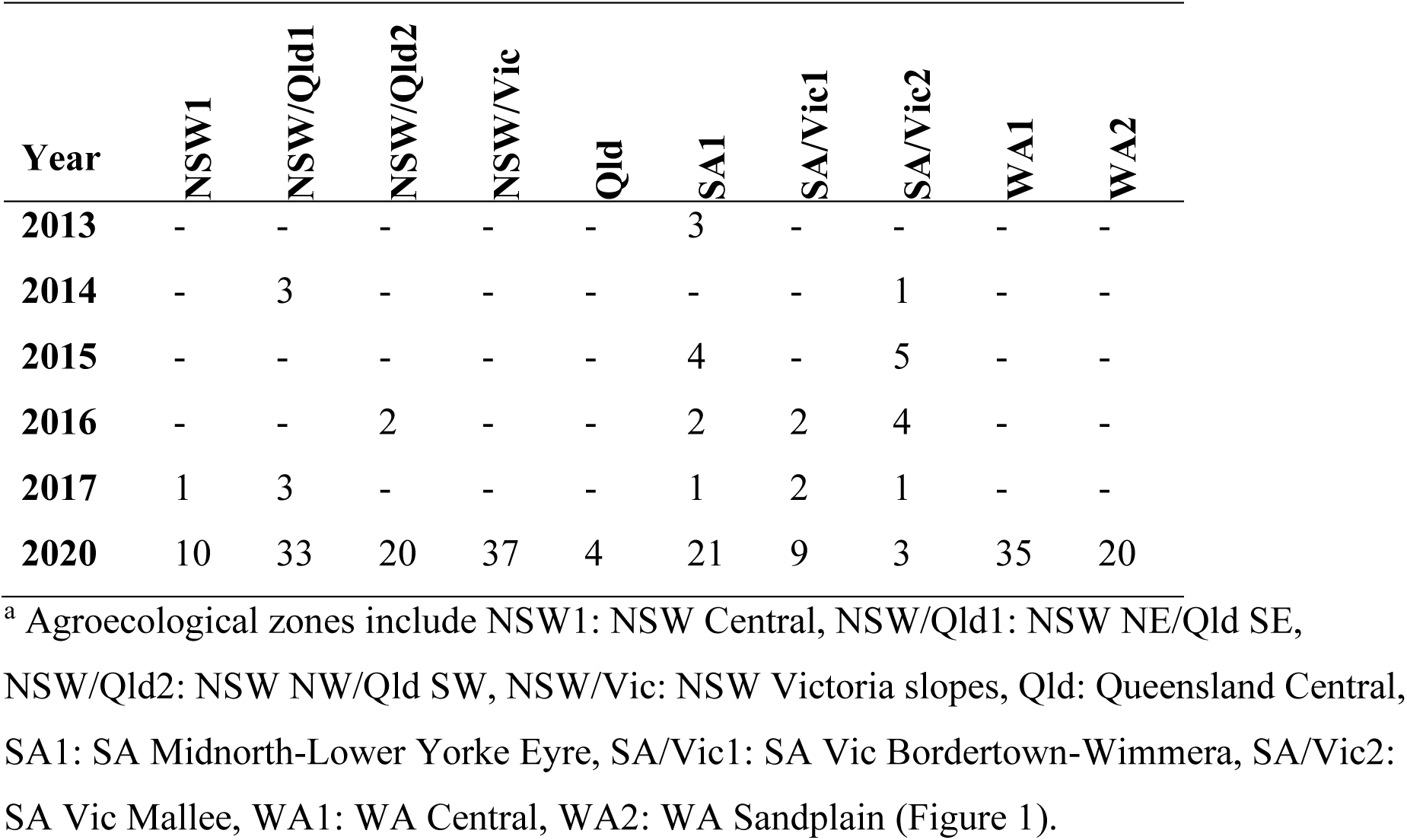
Number of Australian *Ascochyta rabiei* isolates used for whole genome genotyping by year of collection and agroecological zone^a^.

| Year | NSW1 | NSW/Qld1 | NSW/Qld2 | NSW/Vic | Qld | SA1 | SA/Vic1 | SA/Vic2 | WA1 | WA2 |
| --- | --- | --- | --- | --- | --- | --- | --- | --- | --- | --- |
| 2013 | - | - | - | - | - | 3 | - | - | - | - |
| 2014 | - | 3 | - | - | - | - | - | 1 | - | - |
| 2015 | - | - | - | - | - | 4 | - | 5 | - | - |
| 2016 | - | - | 2 | - | - | 2 | 2 | 4 | - | - |
| 2017 | 1 | 3 | - | - | - | 1 | 2 | 1 | - | - |
| 2020 | 10 | 33 | 20 | 37 | 4 | 21 | 9 | 3 | 35 | 20 |

<sup>a</sup> Agroecological zones include NSW1: NSW Central, NSW/Qld1: NSW NE/Qld SE, NSW/Qld2: NSW NW/Qld SW, NSW/Vic: NSW Victoria slopes, Qld: Queensland Central, SA1: SA Midnorth-Lower Yorke Eyre, SA/Vic1: SA Vic Bordertown-Wimmera, SA/Vic2: SA Vic Mallee, WA1: WA Central, WA2: WA Sandplain (Figure 1).

For DNA extraction, single-conidium isolates were grown in 20 ml Czapek-Dox broth (Oxoid, Australia) in 50 ml Falcon® tubes (TechnoPlas, Australia) on a shaker (Ratek orbital shaker incubator) at 180 revolutions per minute at room temperature. Two-week-old mycelia were harvested, dried on autoclaved paper towel, flash-frozen in liquid nitrogen and lyophilized for 48 hours. The lyophilized mycelium of each isolate was ground to a fine powder in a 2 mL microcentrifuge tube using stainless steel beads (2.8 mm diameter; Sigma-Aldrich) in a FastPrep-24 kit (MP Biomedicals) (6.5 m/s for 15 s). Genomic DNA was extracted from each isolate using a DNeasy Plant Mini kit (Qiagen) according to the manufacturer’s instructions, with the following modifications. Buffer AP1 was pre-heated to 65°C before 450 µL buffer AP1 was added to the tubes on ground frozen tissue, which were subsequently incubated at 65°C for 15 minutes. Also, 160 µL buffer P3 was added to the tubes and incubated on ice for 10 minutes. Finally, DNA was eluted twice in 35 µL filter-sterilized 10 mM Tris-HCl (pH 8.5). The DNA was quantified using a Qubit dsDNA Broad Range kit (Qubit 3.0, Invitrogen) and the integrity was assessed by agarose gel electrophoresis (1% wt/vol agarose in Tris-acetate-EDTA) amended with 1:20,000 (v/v) nucleic acid stain GelRed (Biotium). The isolates were confirmed as *A. rabiei* and their mating-type was assigned using the *A. rabiei*-specific multiplex PCR assay described by Barve et al. (15).

### Plant materials, inoculum and bioassay conditions

A differential host set of chickpea genotypes with industry established disease reactions in 2021, that is, ICC3996 (moderately resistant), Genesis 090 (moderately susceptible), PBA Seamer (moderately resistant), and PBA HatTrick (moderately susceptible) was used to assess isolate aggressiveness (note that these ratings have since changed, see the current disease ratings at the [GRDC NVT Disease Ratings](https://nvt.grdc.com.au/nvt-disease-ratings?crop=17&state=NSW)). The highly susceptible cultivar Kyabra was used as a susceptible check in the bioassays. Seeds were provided by the National Chickpea Breeding Program, Tamworth, New South Wales (NSW), Australia. For disease severity bioassays, seedlings were grown in 15 cm diameter pots containing commercial grade potting mix (Richgro premium mix), fertilized with Nitrosol®, Amsgrow® (4.5 mL/L) every two weeks and watered as required. Plants were grown and maintained at 23 ± 2°C under an 8-hour night/16-hour day photoperiod with near-UV light irradiation of 350–400 nm (Supplied by Valoya LED grow lights, Finland) at Griffith University, Nathan campus, Queensland, Australia.

Single-spored Australian *A. rabiei* isolates were cultured on V8 juice agar plates and maintained at 22 ± 2°C with 12-hour night/12-hour day near-UV light irradiation (350–400 nm) for 14 days. Subsequently, spore suspensions were prepared by harvesting pycnidiospores with sterile water, filtering through sterile cheese cloth (250 mm) and the final concentration of inoculum was adjusted to 1×10^5^ spores/mL using a haemocytometer. Two to three drops of Tween 20 (0.02% v/v) were added per 100 mL of spore suspension as a surfactant. Bioassays were conducted in a completely randomized design with two replicates for each genotype × isolate combination and five seedlings per pot in each replicate. Inoculation was carried out using the modified mini-dome technique (26,27). Seedlings were misted to run-off with a single spore inoculum and placed randomly in dark and humid sealed 20 L plastic crates (one crate/isolate). At 24 hours post inoculation (hpi), crate lids were removed, and the seedlings were placed back into the 23 ± 2°C and 8-hour night/16-hour day growing conditions until tissue sample collection and disease assessment (28). Humidity was maintained at 70-75% by misting every 2 days or as needed.

### Assessment of host reaction and assignment of isolate aggressiveness

The disease reaction of each isolate on each of the host genotypes was assessed using the qualitative 1–9 scale of Singh (1981) (29) at 21 days post inoculation (dpi) where; scores of 1 or 3 represented ‘low disease severity’; 5 represented ‘moderate disease severity’ without significant stem infection, and 7 or 9 represented ‘high disease severity’ with stem lesions that led to impact on transpiration, photosynthesis and possible stem breakage. Isolates able to produce a leaf score of at least 7 on >80% and a stem score of at least 7 on >10% of the seedlings of any of the host genotypes assessed were classified as ‘highly aggressive’. Classification into pathogenicity groups (PG) was determined by the following criteria: low disease response on all hosts (except Kyabra, which is the susceptible check) – PG0; high on PBA HatTrick and low on ICC3996 and Genesis090 – PG1; high on PBA HatTrick and a combination of low and moderate on ICC3996 and Genesis090 – PG2; high on PBA HatTrick and Moderate on both ICC3996 and Genesis090 – PG3; high on PBA HatTrick and Genesis090 and moderate on ICC3996 – PG4; and high on ICC3996, Genesis090 and PBA HatTrick – PG5, as described by Bar et al. (6). Isolate aggressiveness and collection metadata were deposited in the Ascochyta dashboard and are available at https://bit.ly/asco-dashboard.

### Whole genome sequencing and variant calling

Whole genome sequencing was conducted at the Australian Genome Research Facility (AGRF, Melbourne) using Illumina Nextera library preparation and NextSeq sequencing platform. To assess the reproducibility of variant calling and allow for stringent filtering of the raw data, three DNA samples (AR0039, AR0052, and AR0242) were replicated within and across plates as technical controls. The raw sequencing data was deposited to the NCBI Short Read Archive (SRA) and is available through BioProject PRJNA1175002.

Whole genome sequencing data processing, mapping and reference based variant discovery was performed on the Griffith University High-Performance Computing Cluster ‘Gowonda’. Raw sequencing reads were pre-processed to remove adaptor sequences and to trim low-quality bases from the raw reads using fastp v0.20.1 (30) with default parameters. Trimmed reads were aligned to *A. rabiei* reference genome, strain ArME14 (NCBI accession [GCA_004011695.2](https://www.ncbi.nlm.nih.gov/datasets/genome/GCF_004011695.2/); (31)) using bwa-mem v0.7.17-r1188 (32), followed by several commands of Samtools v1.12 (33), Sambamba v1.0.0 (34) and Picard v2.27.5 (35) to specify a Read Group for each sample, mark duplicates, and convert the alignments into sorted coordinates indexed BAM files, respectively. The alignment files were then processed using Freebayes v1.3.5 (36) to call single nucleotide variants (SNVs) and insertion/deletions (INDELs) from all samples.

### Quality assessment and filtering

Quality filtering of the variant call format (VCF) file produced by Freebayes was conducted using vcfTools v0.1.16 (37) and SnpSift v4.3 (38) on the Linux cluster at the Cornell University BioHPC computing cloud (Ithaca, New York) in multiple steps. First, the dataset was filtered using vcfTools to retain only high-quality genotype calls (GT defined as the assignment of alleles at each site for each individual) with a minimum depth of five and minimum quality of 20 (*--minDP 5 --minGQ 20 --recode --recode-INFO-all*). The INFO column was recoded every time the data was filtered in vcfTools (*--recode --recode-INFO-all*) to retain all the INFO in the original files and re-estimate INFO values after applying the specified filters. Sites with more than twice the total expected read depth were removed using SnpSift to exclude potential errors due to mapping/assembly errors in repetitive regions (*filter “(DP<42000)”*). Only biallelic SNVs were retained and sites with more than 50% missing data and average quality <30 were removed (*--min-alleles 2 --max-alleles 2 --max-missing 0.5 --minQ 30 --recode --recode-INFO-all*). Individual levels of missing data were estimated using vcfTools (*--missing-indv*) which was estimated to be <10% for all individuals, thus, no isolates were filtered out. Sites were filtered for polymorphism, average depth and missingness, and only those with a minor allele frequency of 0.01 or greater, average depth of 20 and missing data <10% were retained (*--min-meanDP 20 --max-missing 0.9 --maf 0.01 -- recode --recode-INFO-all*). The entire dataset was subsequently filtered using SnpSift, removing any sites with different genotype calls (GT) calls between the replicated DNA samples to ensure that only high-quality loci with reproducible results were retained. The three replicated technical controls were subsequently removed from the dataset after establishing that bitwise genetic distance estimated for replicated DNA samples was zero. Only polymorphic sites were retained using a polymorphism threshold of 1% (minor allele frequency = 0.01) (*--remove list.txt --maf 0.01 --recode --recode-INFO-all*). The polymorphic SNVs are hereafter referred to as single nucleotide polymorphisms (SNPs).

In order to investigate the location of SNPs in high-GC or low-GC regions of the genome, the program OcculterCut v1.1 (39) was used to scan the genomes of *A. rabiei* strain ArME14 and *Leptosphaeria maculans* strain v23.1.3 (NCBI accession [ASM23037v1](https://www.ncbi.nlm.nih.gov/datasets/genome/GCF_000230375.1/); (40)) to assess and compare GC content and distribution. Gene annotations, GC content and frequency of variants were subsequently plotted against the ArME14 genome using CIRCOS (41). Variants were annotated against the gene models of the ArME14 *A. rabiei* reference genome using SnpEff v4.3 (38) to identify SNPs with high, moderate, low, or modifier (intergenic) impact. For population genetic analyses that require selectively neutral markers (population differentiation, indices of genetic and genotypic diversity, and *a priori* population structure), a subset of variants after removal of high and medium impact sites was used.

### Identification of clonal lineages

An essential step in population genomic analyses of clonal organisms is identification of clones, often defined as unique multi-locus genotype (MLGs) without inflating the number of clones due to variant-calling errors or missing data (42–45). Our preliminary analyses of the *A. rabiei* population in Australia demonstrated that many unique MLGs differed at only a small proportion of loci (Fig. S1). We, therefore, used the functions *cutoff_predictor* and *mlg.filter* in *poppr*, to collapse all MLGs with genetic distance smaller than the estimated threshold of 0.0045 into the same MLL. Therefore, all MLGs with nine or fewer differences were grouped into the same MLL (45,46).

### Population differentiation and structure

The term population is used here to refer to a group of isolates defined under certain geographical or temporal criteria, *e.g.*, collected in the same year or within the same agroecological zone. Since our dataset included individuals from 2013 to 2020, we first tested for temporal population differentiation to investigate whether individuals collected in different years from the same agroecological zones can be pooled together. Analysis of molecular variance (AMOVA) was conducted using *ade4* v1.7-18 (47) in *poppr* v2.9.3 (46) to test for population structuring within and among different years without making assumptions about Hardy-Weinberg equilibrium. For this, we used the subset of low impact and intergenic SNPs (total of 2,554 sites) to investigate the partitioning of genetic variation within and among locations, zones, and states in each year and among years. The significance of Phi (ϕ) statistic was tested using 999 permutations. Due to the small number of individuals in the 2013 and 2014 populations, the AMOVA analysis was repeated after removing these populations from the dataset.

For Pairwise Hedrick’s G’’ST (48,49), which is a standardized measure of population differentiation calculated by dividing G_ST_ for a given marker by the maximum theoretical GST, was estimated using the package *mmod* v1.3.3 (50). Jost’s pairwise index of differentiation (D), which measures allelic differentiation between populations (51), was also estimated using *mmod*. Both G’’ST and Jost’s D may range from 0 (no differentiation) to 1 (complete differentiation). Dendrograms of genetic distance among individuals was produced through neighbour-joining (NJ) method in *poppr* based on bitwise genetic distance (52) and also using Maximum Likelihood algorithm in RAxML v8 (53) using acquisition bias correction for SNP datasets (54).

To investigate temporal and geographical population differentiation, Discriminant Analysis of Principal Components (DAPC) analyses were also conducted in R package *adegenet* v2.1.5 (55) with *a priori* populations defined as year of collection and also agroecological zones. Unlike AMOVA, DAPC allowed us to include 2013 and 2014 populations as this approach is not affected by smaller sample sizes. The optimal number of principal components (PCs) to retain for each analysis was determined using the *xvalDapc* function and the DAPC analyses were conducted using *adegenet*.

The existence of an underlying structure without *a priori* population assignment was assessed using DAPC, which does not assume linkage equilibrium of loci. The reduced SNP dataset including only low impact and intergenic SNPs was used for this analysis (a total of 2,554 sites). The optimal number of clusters was determined using the function *find*.*clusters,* and *dapc* was used to assign individuals into clusters, retaining the number of principal components encompassing 94% of the cumulative variance.

### Indices of gene and genotypic diversity

Lack of significant genetic differentiation among populations from different years allowed us to pool the individuals collected from the same agroecological zones in different years together to calculate indices of genetic diversity using *poppr*. Allele frequencies for all loci were calculated using vcfTools and visualized in R. The genetic distances between *A. rabiei* isolates were estimated with the *diss*.*dist* function in *poppr*.

### Identification of loci associated with isolate aggressiveness

To identify loci underlying aggressiveness, the entire data set including SNPs and INDELs with high, moderate, low, and modifier impacts were used (a total of 3,283 sites). Linkage disequilibrium (LD) decay plots generated in GAPIT3 (56) showed long-range linkage of loci in *A. rabiei* (Fig. S2). Therefore, the homoplasy-based genome-wide association (GWAS) analysis implemented in POUTINE (57) was implemented using the entire dataset (total of 3,283 sites with high, moderate, low, and modifier effect). Ancestral trees required as input for POUTINE were generated in RAxML v8 (53) using acquisition bias correction for SNP datasets (54). POUTINE can only handle binary phenotypes; therefore, separate analyses were conducted to find genotype associations with high/low, high/medium, and low/medium aggressiveness.

We also performed a second GWAS analysis in pyseer v1.3.10 (58) using the qualitative 1–9 disease severity scale of Singh (29) as phenotypes, with minimum and maximum allele frequencies of 0.04 and 0.98, respectively, and a whole genome (elastic net) association model. Pyseer includes strategies to account for clonality and population structure, and *p* value corrections for multiple hypothesis testing, which can lead to false positives in traditional GWAS approaches.

As a third approach to detect loci underlying aggressiveness, we used DAPC, which is not affected by linkage of loci. In order to remove the complicating effect of population structure, association of SNPs with aggressiveness of isolates was assessed within each of the three clusters identified in the initial DAPC analysis using the entire dataset (total of 3,283 sites with high, moderate, low, and modifier effect). Sub-populations within each cluster were defined as groups of isolates belonging to the same aggressiveness group (high, medium, and low) on each of the chickpea varieties tested, and their differentiation was assessed using DAPC. The optimal number of principal components (PCs) to retain for each analysis was determined using the *xvalDapc* function and the DAPC analyses were conducted using *adegenet*. The function *loadingplot* was subsequently used to find the SNPs contributing to the separation of isolates belonging to each aggressiveness group.

Finally, pairs of closely related isolates (different at < 15 sites) with contrasting aggressiveness (pathogenicity group 0/1 versus 4/5) were compared to identify genetic variations potentially associated with aggressiveness. Upon identification of variations potentially associated with aggressiveness of *A. rabiei*, the impact and location of the site was further investigated, upstream and downstream coding sequences were identified, and predicted proteins were used in NCBI BLASTp searches to characterize their potential function. The probability of the encoded proteins to be fungal effectors was investigated using Predector v1.2.7 (59) and EffectorP v3.0 (60). For accurate effector prediction, SignalP v6 (61) and Phobius (62) were implemented for computational evidence of secretion for predicted effectors, as EffectorP recognizes a cytoplasmic signal also in intracellular, non-secreted proteins (60). For intergenic SNPs, we utilised the Neural Network Promoter Prediction server (https://www.fruitfly.org/seq_tools/nnppHelp.html) to explore whether these variants were located within promoter regions, potentially influencing gene expression.

## RESULTS

### Species identification and mating-type screening

Mating-type PCR screening of all *A. rabiei* isolates (*n* = 230) confirmed their identity as *A. rabiei* and resulted in an amplicon of approximately 400 bp for all isolates, as expected for MAT1-2.

### Aggressiveness of *A. rabiei* isolates

Of the 230 isolates assessed, 36 and 24 were aggressive (16%) and highly aggressive (10%), identified as belonging to pathogenicity groups four and five, respectively. A significant increase in frequency of detection of isolates able to cause severe damage on “moderately resistant” cultivars was observed since 2016 (Fig. 2). In particular, the proportion of isolates that were highly aggressive on PBA HatTrick and Genesis 090, both widely adopted cultivars, had steadily increased. Also, isolates able to cause severe disease on a major resistance source used within the Australian chickpea breeding program, ICC3996, had emerged. Of most immediate concern for the growers was the proportion of isolates able to cause severe disease on a relatively recently released variety PBA Seamer, increasing from 7.4% in 2018 to 44.4% in 2020 (Fig. 2).

**Fig. 2.**
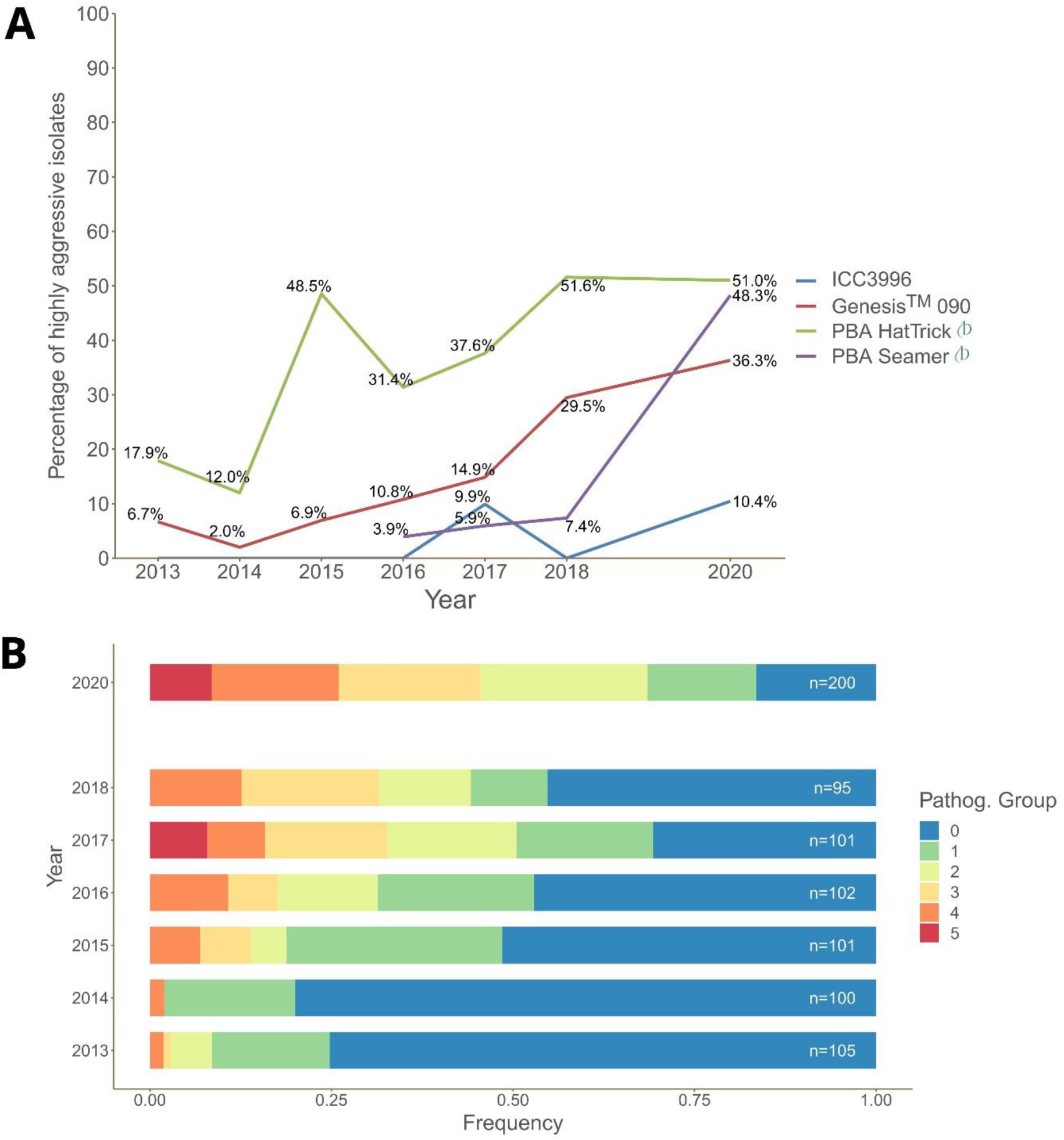
Summary of *Ascochyta rabiei* aggressiveness between 2013-2020. **A**. Frequency of highly aggressive isolates (able to produce a leaf score of at least 7 on >80% and a stem score of at least 7 on >10% of the seedlings, see Assessment of host reaction in Methods section for details) on each chickpea host genotype at each year. **B**. Distribution of isolates by Pathogenicity Group classification (cumulative disease response to the set of host genotypes, see Assessment of host reaction in Methods section for details) in each year. These figures were retrieved from the *[Ascochyta dashboard](https://shinotate.pp18000.cloud.edu.au/shiny/Asco_dashboard/)* on June 30^th^ 2022. Updated information from recent years can be found at https://bit.ly/asco-dashboard.

### Quality assessment and filtering

Illumina sequencing of the 230 *A. rabiei* isolates resulted in a minimum of 35× coverage per isolate with an average GC% nucleotide content of 48-50%. Filtering of the VCF file output from Freebayes for high quality and depth variants resulted in a total of 13,259 variants for 230 *A. rabiei* isolates. Further filtering for quality, depth, minor allele frequency of 0.01 and removal of loci that gave contradictory results for internal replicates resulted in a total of 3,283 high quality variants sites, including 2,723 SNPs (the term SNP here is used to refer to those SNVs that passed the polymorphism threshold of 1%) and 560 INDELs (insertions and deletions).

SNP annotation in SnpEff assigned 13, 156, and 134 SNPs in the final dataset as high, moderate, and low impact (present in the coding region of a gene). The remaining 2,420 SNPs were identified as modifier (intergenic) loci. For INDELs, seven, 49, 11 and 493 sites were identified as high, moderate, low, and modifier impact. High and moderate impact variants are predicted to have a disruptive effect on protein function (truncation, loss of function, or nonsense mediated decay) or change protein effectiveness while low impact variants are assumed to be harmless and unlikely to change protein function. Modifier (or intergenic) variants, on the other hand, reside in non-coding regions of the genome, where predictions are difficult or there is no evidence of impact (38). For population genetic analyses that require selectively neutral markers, SNPs with high and moderate impact were removed from the data set and only biallelic SNPs with low and modifier impacts were used (total of 2,554 sites). Analysis using OcculterCut revealed that 22.5% of *A. rabiei* genome consisted of AT-rich regions (Fig. 3), which also harboured the majority of the SNPs. The position of SNPs on the *A. rabiei* genome and in relation to GC percent and gene content is shown in Fig. S3.

**Fig. 3.**
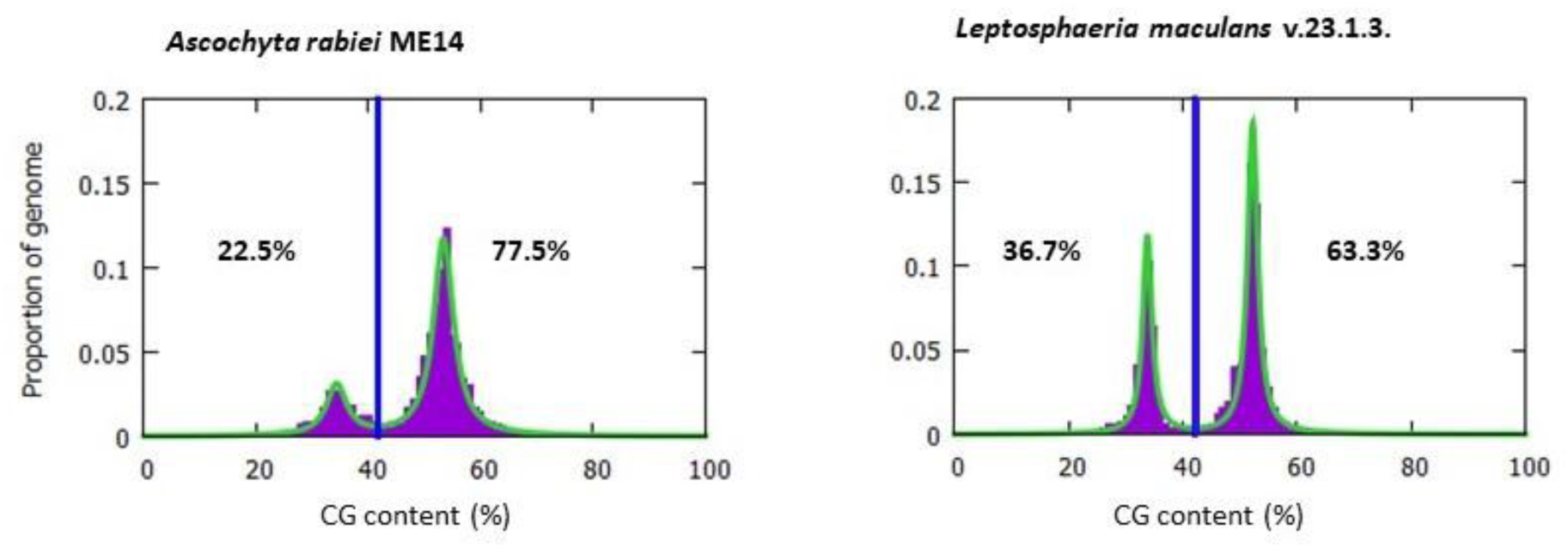
The GC content distribution of *Ascochyta rabiei* strain ArME14 in comparison with another plant pathogenic fungus, *Leptosphaeria maculans* strain v23.1.3. Vertical blue lines show the GC cut-off points selected by OcculterCut (37) to classify genome segments into AT-rich and GC-balanced regions. The percentage values shown on the left and right sides of the vertical blue lines indicate the percentage of the genome classified as AT-rich and GC-balanced, respectively.

### Population differentiation

No significant differentiation was detected among *A. rabiei* collected in different years after removing the smaller 2013 and 2014 populations (Tables 2 and 3). Low but statistically significant genetic differentiation was detected between populations from different States (14%) and agroecological zones (21%) and most of the genetic variation (66%) was found among individuals within states and agroecological zones (*P* = 0.001). This indicated low but significant structuring of the population based on isolate geographical location, while no significant temporal differentiation was detected among populations collected from 2015 to 2020 using AMOVA. Adding 2013 and 2014 populations to the analyses did not change the results of AMOVA analyses (Tables S2 and S3).

**Table 2.** Summary of analysis of molecular variance (AMOVA) results for 223 *Ascochyta rabiei* isolates collected in 2015, 2016, 2017, and 2020 from different States and locations in Australia.

| Source | d.f.<sup>a</sup> | SS<sup>b</sup> | MS<sup>c</sup> | variance | proportion of variation | P value |
| --- | --- | --- | --- | --- | --- | --- |
| Among years | 3 | 1,471.32 | 490.44 | 0.45 | 0% | 0.423 |
| Among States within years | 10 | 7,834.44 | 783.44 | 31.70 | 13.5% | 0.001 |
| Among locations within States | 34 | 12,111.72 | 367.02 | 46.57 | 20.5% | 0.001 |
| Within samples | 175 | 25,207.65 | 144.87 | 144.87 | 66.0% | 0.001 |
| Total | 222 | 46,625.130 | 211.93 | 222.69 | 100.0% | - |

<sup>a</sup>Degrees of freedom
<sup>b</sup>Sum of squared observations
<sup>c</sup>Mean of squared observations

**Table 3.** Summary of analysis of molecular variance (AMOVA) results for 223 *Ascochyta rabiei* isolates collected in 2015, 2016, 2017, and 2020 from different agroecological zones and locations in Australia.

| Source | d.f.<sup>a</sup> | SS<sup>b</sup> | MS<sup>c</sup> | variance | proportion<br>of variation | P<br>value |
| --- | --- | --- | --- | --- | --- | --- |
| Among years | 3 | 1,471.32 | 490.44 | 4.65 | 2.0% | 0.185 |
| Among zones within years | 19 | 13,297.69 | 738.76 | 41.46 | 18.0% | 0.001 |
| Among locations within<br>Zones | 25 | 6.648.48 | 265.93 | 30.15 | 13.5% | 0.001 |
| Within samples | 175 | 25,207.65 | 144.87 | 144.87 | 66.5% | 0.001 |
| Total | 222 | 46,625.13 | 211.93 | 221.14 | 100.0% | - |

<sup>a</sup>Degrees of freedom
<sup>b</sup>Sum of squared observations
<sup>c</sup>Mean of squared observations

Jost’s and Hedrick’s G’’ST indices of pairwise index of differentiation (D), were also low between populations from different years (Table 4). On the other hand, DAPC analysis using year of collection as *a priori* population assignment, which is not affected by population size and allowed us to also include populations from 2013 and 2014 in the analyses, separated the majority of the 2013 isolates from the rest of the population (Fig. 4A). Structuring of the *A. rabiei* population based on geographical location (agroecological zones) was confirmed by the results from the DAPC analysis (Fig. 4B).

**Fig. 4.**
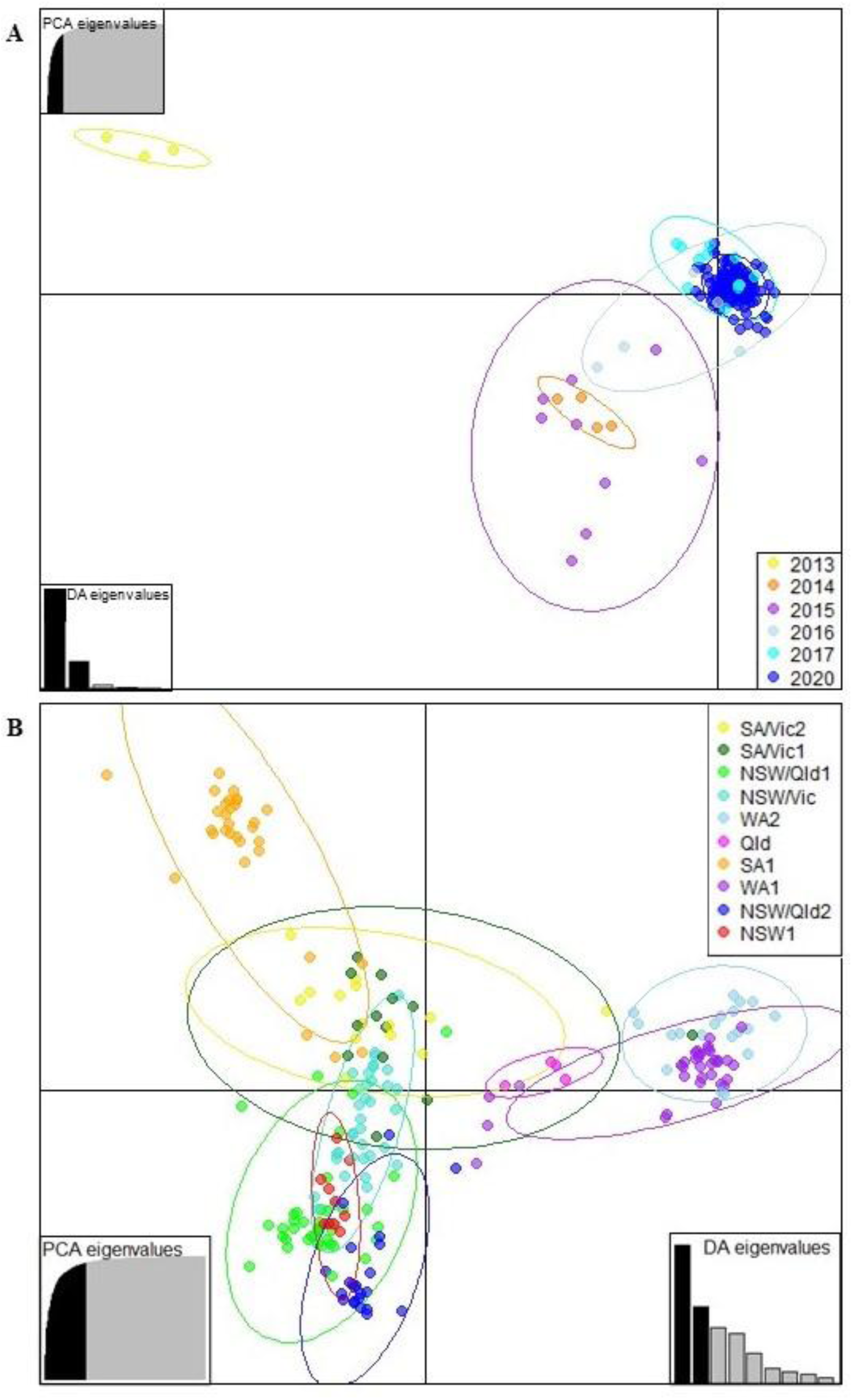
**A.** Discriminant Analysis of Principal Components (DAPC) for *Ascochyta rabiei* isolates collected in Australia from 2013 to 2020, using year of collection as *a priori* populations. The optimal number of principal components (PCs) to retain for each analysis was determined by *xvalDapc* function (55) to be 32, encompassing 88% of the cumulative variance. The ellipses represent the maximum area spanned by 95% of the data in a population by year of collection. **B.** Discriminant Analysis of Principal Components (DAPC) for *Ascochyta rabiei* isolates collected in Australia from 2013 to 2020, using agroecological zones (Table 1) as *a priori* populations. The optimal number of principal components (PCs) to retain for each analysis was determined by *xvalDapc* function (55) to be 58, encompassing 98% of the cumulative variance. The ellipses represent the maximum area spanned by 95% of the data in an agroecological zones.

**Table 4.** Pairwise population differentiation (Jost’s D) between *Ascochyta rabiei* populations collected in different years in Australia shown at the bottom of the table, and Pairwise population differentiation (Hedrick’s G’’ST) between *Ascochyta rabiei* populations collected in different years in Australia at the top of the table in parentheses.

|  | 2015 | 2016 | 2017 | 2020 |
| --- | --- | --- | --- | --- |
| 2015 | 0 | (0.142) | (0.206) | (0.232) |
| 2016 | 0.010 | 0 | (0.039) | (0.047) |
| 2017 | 0.015 | 0.002 | 0 | (0.078) |
| 2020 | 0.020 | 0.003 | 0.006 | 0 |

**Table 5.** Genetic variations predicted to be associated with aggressiveness of *Ascochyta rabiei* isolates on different chickpea cultivars in Australia, detected using both GWAS (pyseer) and DAPC analyses.

| Variation position | P value |  |  |  | Variation type, impact and location on the genome | Putative protein function | Predector Score (EffectorP 3.0 score)<sup>a</sup> | Repeat family |
| --- | --- | --- | --- | --- | --- | --- | --- | --- |
|  | Genesis090 | ICC3996 | PBA HatTrick | PBA Seamer |  |  |  |  |
| ctg_01_1654657 | - | - | - | 0.0273 | intergenic_region, modifier, EKO05_000484-EKO05_000485 | EKO05_000484: hypothetical protein with high similarity to fungal GTPases, EKO05_000485: hypothetical DNA binding protein with a transmembrane domain | non-effectors | LTR/Copia |
| ctg_01_1665154 | - | - | - | 0.0273 | same as ctg01_1654657 |  |  |  |
| ctg_02_2690139 | - | - | - | 0.0273 | missense_variant, moderate impact, EKO05_001768 | hypothetical protein with similarity to fungal transcription factors with a zinc-finger domain | non-effector | - |
| ctg_02_364191 | - | - | - | 0.0273 | intergenic_region, modifier, EKO05_001071-EKO05_001072 | EKO05_001071: hypothetical HET domain containing protein with a transmembrane domain, EKO05_001072: hypothetical phosphotransferase | non-effectors | LTR/Gypsy |
| ctg_03_1415178 | - | - | - | 0.0273 | intergenic_region, modifier, EKO05_002061-EKO05_002062 | EKO05_002061: hypothetical protein of unknown function, EKO05_002062: **hypothetical apoplastic effector** | EKO05_002061: non-effector, EKO05_002062: 0.676 (0.709, Apoplastic effector) | LTR/Gypsy |
| ctg_08_448981 | - | - | - | 0.0364 | intergenic_region, modifier, EKO05_005280-EKO05_005281 | EKO05_005280-EKO05: **hypothetical apoplastic effector**, EKO05_005281: hypothetical protein of unknown function | EKO05_005280: - 2.911 (0.509, Apoplastic effector), EKO05_005281: non-effector | LTR/Gypsy |
| ctg_08_98214 | - | - | - | 0.0273 | intergenic_region, modifier, EKO05_005201-EKO05_005202 | EKO05_005201: **hypothetical apoplasic effector**, EKO05_005202: hypothetical protein of unknown function | EKO05_005201: - 1.476 (0.654, Apoplasic effector), EKO05_005202: - 3.378 (0.675, Cytoplasmic effector but no signal peptide detected) | LTR/Gypsy |
| ctg_10_10643 | - | - | 0.00895 | 0.0374 | synonymous_variant, low impact, EKO05_006140<br>On a potential promoter region with prediction score of 0.9 | hypothetical protein of unknown function | -2.614 (0.522, Cytoplasmic effector but no signal peptide detected) | DNA/Kolobok-H |
| ctg_10_157955 | - | - | - | 0.0364 | intergenic_region, modifier, EKO05_006148-EKO05_006149 | EKO05_006148: hypothetical protein of unknown function, EKO05_006149: hypothetical oxidoreductase | non-effectors | DNA/TcMar-Fot1 |
| ctg_10_157964 | - | - | - | 0.0438 | Same as ctg_10_157955 |  |  |  |
| ctg_10_158053 | - | - | - | 0.0364 | Same as ctg_10_157955 |  |  |  |
| ctg_12_1234964 | - | - | - | 0.0273 | intergenic_region, modifier, EKO05_007471-EKO05_007472 | EKO05_007471: hypothetical protein of unknown function, EKO05_007472: **hypothetical cytoplasmic effector** | EKO05_007471: non-effector, EKO05_007472: - 1.071 (0.535, Cytoplasmic effector, signal peptide detected) | LTR/Gypsy |
| ctg_12_1234984 | - | - | - | 0.0273 | same as ctg_12_1234964 |  |  |  |
| ctg_15_1503696 | 0.0199 | - | - | - | downstream_gene_variant, modifier, EKO05_008905 | EKO05_008905: hypothetical DNA helicase with a transmembrane domain | non-effectors | unknown family |
| ctg_19_1154999 | 0.00638 | - | 0.00293 | 0.000122 | intergenic_region, modifier,<br>EKO05_010492-EKO05_010493 | EKO05_010492: **hypothetical**<br>**apoplastic effector**, EKO05_010493:<br>amino acid transporter with multiple<br>transmembrane domains | EKO05_010492:<br>2.244 (0.895,<br>Apoplastic<br>effector),<br>EKO05_010493:<br>non-effector | LTR/Gypsy |
| ctg_19_1155004 | 0.00638 | - | 0.00293 | 0.000122 | same as ctg_19_1154999 |  |  |  |

<sup>a</sup> Predector score is provided with EffectorP 3.0 score in parentheses. Non-effector indicates a negative Predector score and > 0.5 non-effector score from EffectorP 3.0.

### Genetic diversity

The *A. rabiei* population consisted of 167 multilocus lineages (MLLs) and a genotypic diversity of 0.99 (Simpson’s complement λ = 0.99) (63), high evenness (E_5_ = 0.85) and low clonality (clonal fraction = 26%). Despite the high genotypic diversity, gene diversity (allelic diversity) of the population was very low (H_exp_ ≈ 0) due to the low level of polymorphism in the majority of loci (median allele frequency = 0.017). Therefore, despite identification of 167 unique MLLs, these clonal lineages differed at only a small proportion of loci. The maximum number of allelic differences between genotypes was 820 SNPs and the percentage of SNP differences between genotypes (ratio of the number of observed differences by the number of possible differences) was 14% (Supplementary Fig. S1).

### Population structure

Population structure analysis with no *a priori* population assignment through DAPC analysis detected three distinct clusters in the population (Fig. 1B). Although the Bayesian Information Criterion (BIC) consistently decreased with the number of clusters, K = 3 was selected to best explain the population structure since the BIC value decreased only incrementally when K > 3. The detected three clusters consisted of 43 (Cluster 1), 66 (Cluster 2), and 121 (Cluster 3) isolates, which did not correspond to sampling locations, agroecological zones, or states. The first two discriminant functions explained 96% of the total conserved variance. Neighbor-Joining (NJ) dendrogram of the isolates based on bitwise genetic distance using 2,554 putatively neutral SNPs was generally in agreement with the DAPC results (Fig. 5). Isolates belonging to DAPC cluster 2 were further divided into 2 sub-clusters 2a and 2b in the NJ dendrogram (Fig. 5). Cluster 2a included the highest frequency (53%) of aggressive isolates (pathogenicity groups 4 and 5, followed by cluster 3 (26%) and cluster 2b (19%). Cluster 1 included the lowest frequency of aggressive isolates with only one isolate belonging to pathogenicity group 5 and three isolates belonging to pathogenicity group 4. Interestingly, all isolates in cluster 2a were from WA, except for a single, highly aggressive isolate (AR0304) from Victoria (SA/Vic1). Repeating the NJ analysis using the entire dataset including moderate and high impact loci (a total of 3,283 loci) did not change the results (Fig. S4). Dendrogram generated via Maximum Likelihood algorithm in RAxML resulted in a similar topology (*data not shown*).

**Fig. 5.**
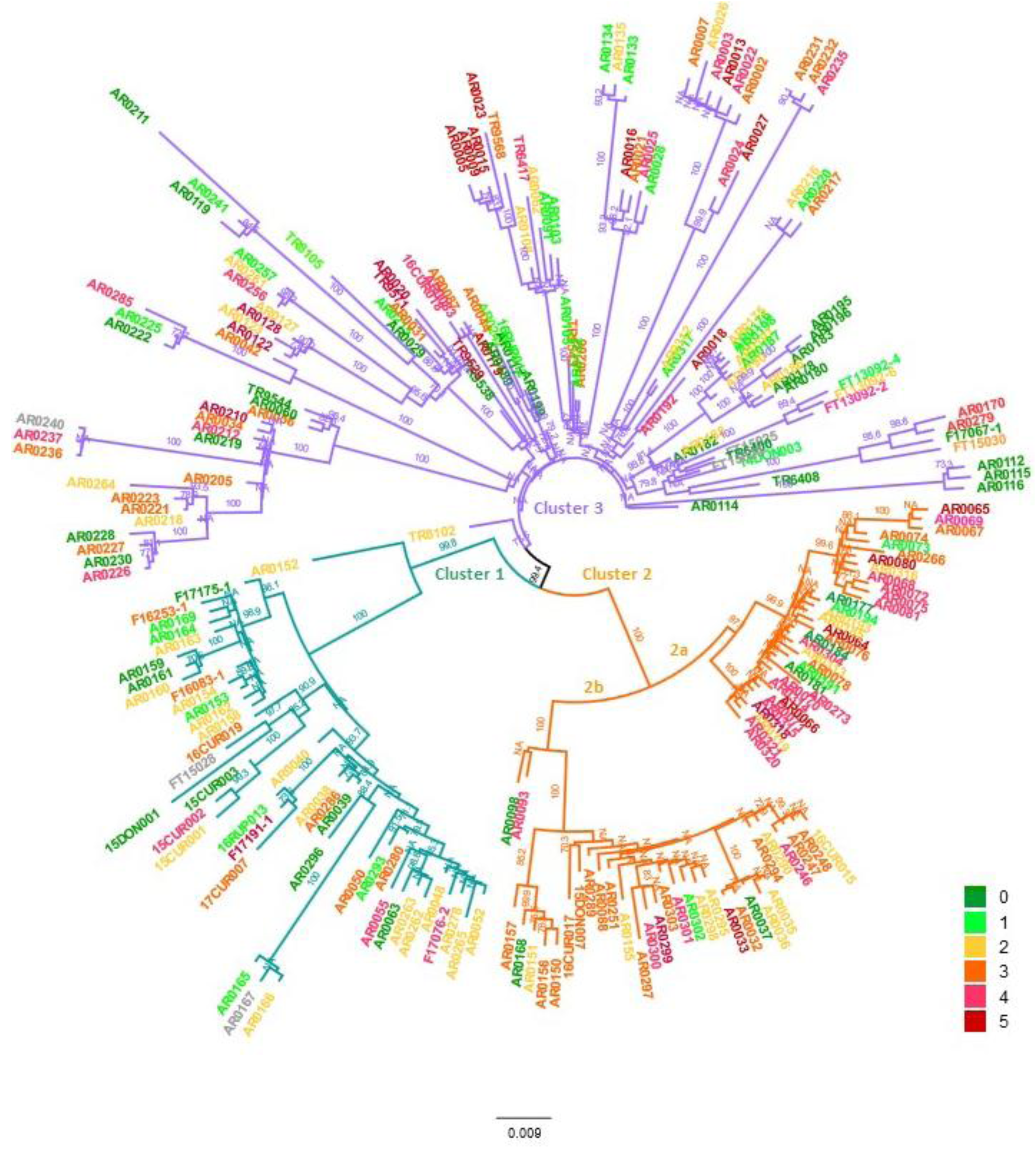
Neighbour-joining dendrogram of 130 *Ascochyta rabiei* isolates sequenced in this study based on bitwise genetic distance using 2,554 putatively neutral single nucleotide polymorphisms. Tip colours correspond to the aggressiveness (pathogenicity group) of the isolates as indicated in the legend. Branches coloured green, orange and purple denote isolates identified in clusters 1, 2, and 3 in the DAPC analysis shown in Figure 1.

### Identification of SNPs potentially associated with *A. rabiei* aggressiveness

GWAS analysis using POUTINE did not find any variants significantly associated with aggressiveness of *A. rabiei* on different host varieties. While multiple mutations were identified to have a significant pointwise *p* value (< 0.05), none survived correction for multiple hypothesis testing (family-wise *p* value > 0.05). GWAS analyses using pyseer, on the other hand, identified 66 variants to be significantly associated with aggressiveness of *A. rabiei* on different host varieties (Fig. 6; Table S5). A SNP at locus ctg_08_1872963 was identified to be significantly associated with *A. rabiei* aggressiveness on all four chickpea cultivars and SNPs at loci ctg_19_1154999 and ctg_19_1155004 were associated with aggressiveness on three cultivars (Genesis090, PBA HatTrick and PBA Seamer). Of the remaining variations, 17 were associated with two chickpea cultivars while 34, six, five, and one variation were associated with PBA Seamer, PBA HatTrick, ICC3996, and Genesis090 only. The majority of detected variations were located at intergenic regions, with only six variations located on coding regions with low to moderate impact (Table S5).

**Fig. 6.**
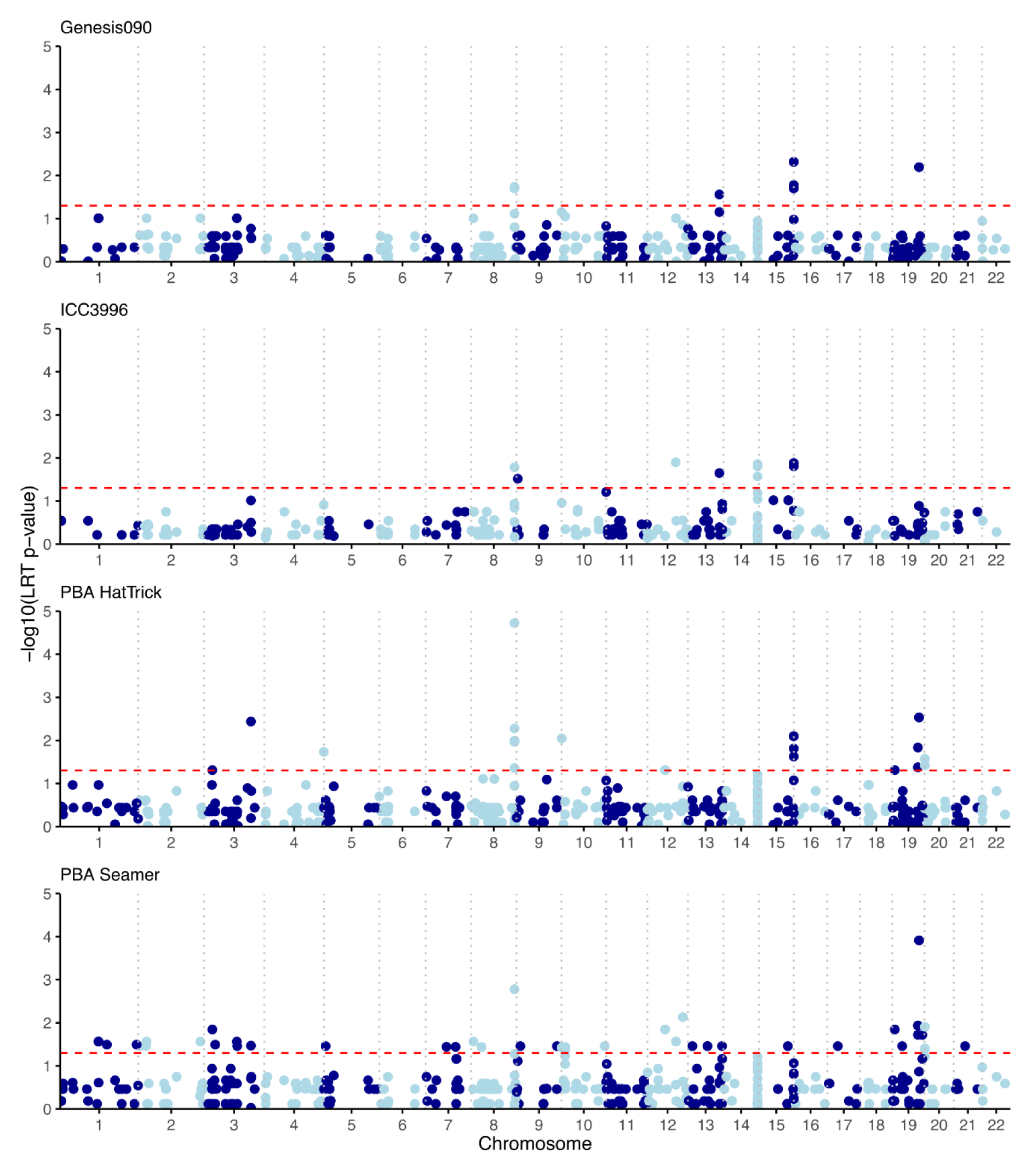
Manhattan plot showing variations with statistically significant association (the significance threshold at *p*-value < 0.05, indicated by the red horizontal dotted line) with aggressiveness of *Ascochyta rabiei* in Australia isolates based on pyseer (58) results.

DAPC analysis for each of the three *A. rabiei* clusters using isolates aggressiveness towards each chickpea variety (low, moderate, and high) as pre-defined populations showed various degrees of separation. A total of 151 variations were identified to be associated with aggressiveness on different chickpea cultivars (loading > 0.005), 22 of which were located on coding regions with low to moderate impact and the rest were intergenic variations (Fig. 7; Table S6). Of these, five SNPs (ctg05_2328653, ctg14_1473121, ctg14_1473129, ctg15_1503638, ctg15_1503666) were found to be linked to aggressiveness of isolates on three chickpea cultivars. Twenty-five SNPs were linked to aggressiveness on two chickpea cultivars (Table S6).

**Fig. 7.**
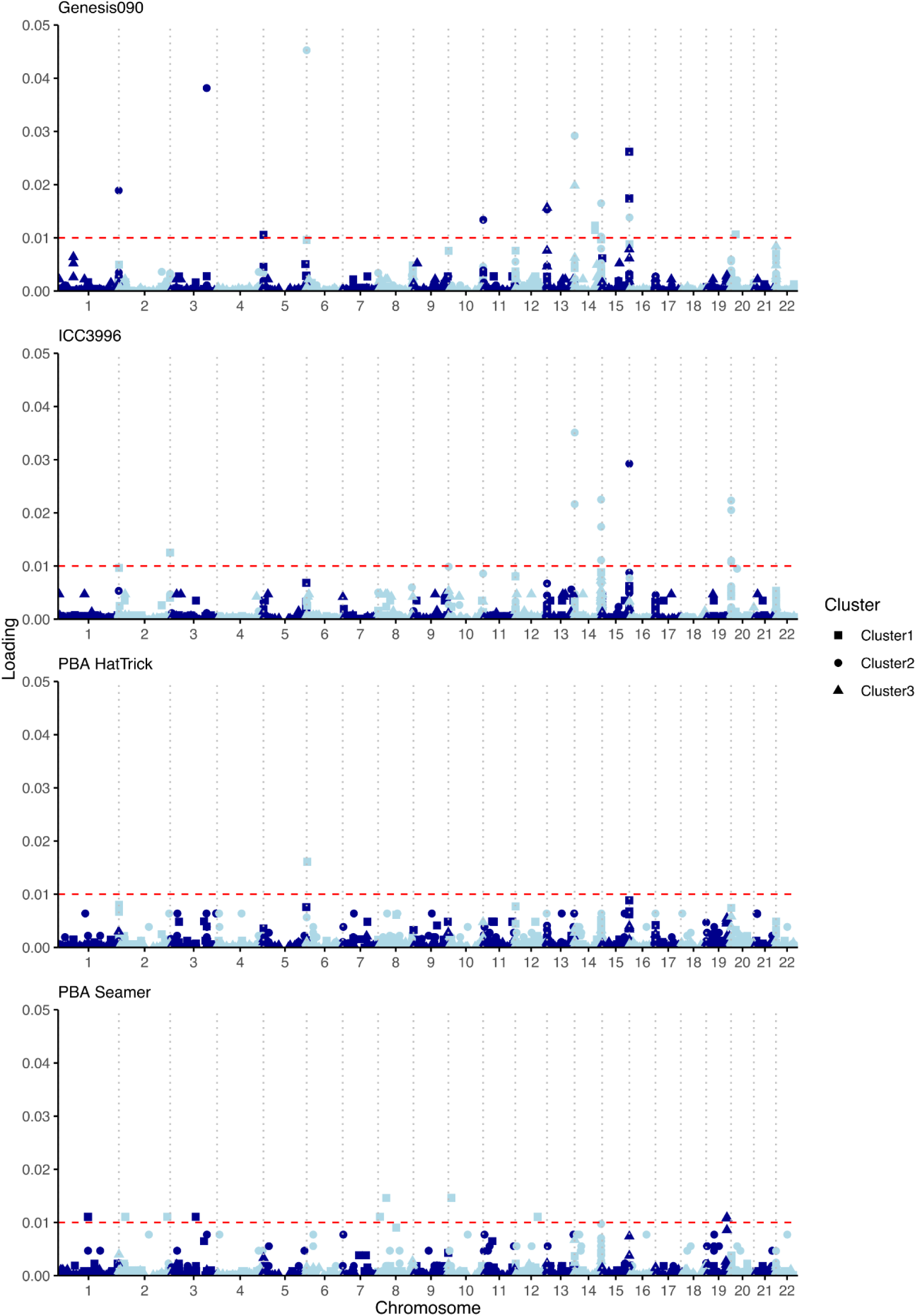
Single nucleotide polymorphism (SNP) markers potentially associated with aggressiveness of *Ascochyta rabiei* isolates on four different chickpea cultivars, namely ICC3996, PBA Seamer, Genesis090, and PBA HatTrick identified using DAPC analyses of three subpopulations (clusters) detected in Australia. The red horizontal line delineates the loading threshold of 0.01.

Sixteen variations were identified to be associated with *A. rabiei* aggressiveness in both DAPC and pyseer analyses (Table 4). Of the remaining variations, although the same variation was not a hit in both analyses, the coding regions associated with the variation were implicated in both DAPC and pyseer analyses. For example, ctg_08_1872963 variation, which was identified in pyseer to be associated with *A. rabiei* aggressiveness on all chickpea cultivars was not a hit in the DAPC analyses, but other closely linked variations with the same predicted impact were identified in the DAPC analysis (intergenic, modifier variations downstream EKO05_005581 coding region).

The phylogenetic analysis showed that aggressive isolates developed independently multiple times across the tree (Fig. 5). Comparison of pairs of closely related individual isolates (different at < 15 loci) with contrasting aggressiveness (pathogenicity group 0/1 versus 4/5) identified genetic variations which were also detected through pyseer and DAPC analyses to be potentially associated with *A. rabiei* aggressiveness (Table S7).

## DISCUSSION

Population genomic analyses of Australian *A. rabiei* isolates using genome-wide single nucleotide polymorphisms confirmed its clonality in Australia, in line with results from previous studies that reported lack of sexual reproduction (6,13,14) as opposed to elsewhere in the world, where *A. rabiei* is known be sexually recombining and genotypically diverse (15–17). The three main clonal lineages detected in the Australian *A. rabiei* population are not evenly distributed in all agroecological zones. Isolates from cluster 3 were found in all regions, while cluster 1 isolates are absent from WA and northern NSW and Qld zones, and cluster 2 isolates absent from northern NSW and Qld zones. Presence of all three lineages in the southern agroecological zones indicated higher genotypic diversity of *A. rabiei* in these regions and is in line with previous findings that South Australia is most likely where *A. rabiei* was first introduced to Australia (13). This also emphasizes the continuous need for biosecurity measures to inhibit long distance dispersal of the pathogen within Australia, which may result in fundamental changes to *A. rabiei* populations in WA and Qld.

The genotyped population was further phenotyped for aggressiveness on different chickpea cultivars to explore genetic variations potentially associated with aggressiveness of *A. rabiei* in Australia. This is the first study to reveal the complex nature of aggressiveness in the Australian *A. rabiei* population, where aggressive isolates have evolved independently several times within different clonal lineages and as a result of multiple small-scale genetic variations.

Classic genome-wide association studies, which are commonly used for identification of loci associated with target phenotypic traits in fungal plant pathogens with frequent cycles of sexual reproduction (64–66), are only relevant in the presence of recombination (67). Such methods are not appropriate for identifying associations of phenotype and genotypes in strongly clonal populations with long-range linkage disequilibrium and population stratification. While non-sexual recombination (for example, parasexuality) may still occur in asexual organisms, significant linkage disequilibrium of loci and high clonality of *A. rabiei* in this study and previous research points to lack of recombination in the Australian *A. rabiei* population. In such situations, homoplasy-counting genome-wide association (GWAS) methods are more appropriate because these methods make use of homoplasic alleles (identical-by-state but not identical-by-descent), which arise repeatedly and independently in different genetic backgrounds, thus, are not in LD (56).

To detect loci that may be associated with aggressiveness of *A. rabiei* in Australia, we made use of DAPC analysis, which does not require assumptions of linkage equilibrium of loci (55), as well as pyseer, which accounts for confounding effects of population structure and clonality (58) and POUTINE (56), a homoplasy-counting GWAS method. DAPC resulted in identification of 151 variations contributing to the separation of *A. rabiei* isolates based on their aggressiveness on different cultivars and within different clonal lineages. Only 22 of these were located on coding sequences: 14 causing missense mutations and eight resulting in synonymous substitutions (Table S6). Pyseer analysis detected 66 variations underlying aggressiveness of *A. rabiei* isolates, only six of which were located on coding regions (two synonymous mutations and four missense mutations, Table S5). Of these, 16 variations were detected in both analyses, including ctg_02_2690139, which resulted in a missense variation on a gene coding for a hypothetical transcription factor with a zinc-finger domain, a known regulator of pathogenicity in fungal plant pathogens (68–70).

Several variations detected using DAPC analyses and pairwise comparison of genetically close isolates with contrasting aggressiveness (ctg05_2388561, ctg14_6054, ctg14_1527717, ctg15_1503638 and ctg20_6490) resulted in missense mutations in genes coding for hypothetical DNA helicases. Several other variations were found in association with synonymous changes in DNA helicase genes or upstream and downstream of putative DNA helicase genes. All these variations were located in telomeric regions. DNA helicases are nucleic acid-dependent ATPases with the ability to separate DNA strands, which is an important step in genome replication, expression, and repair (71). Therefore, helicases have diverse functions in the cell and their substrates substantially vary as they facilitate almost all transactions in nucleic acid metabolism. In fungi, helicases have been associated with post-transcriptional gene silencing (72). Telomere-linked helicases (TLH) have been detected in subtelomeric DNA of filamentous fungi such as *Metarhizium anisopliae* (73), *Ustilago maydis* (74), and *Magnaporthe oryzae* (75,76), and their function is currently unclear, although it may be related to the maintenance of genome stability (76). An RNA helicase gene in *M. oryzae* encodes for the MoDHX35 protein which was reported to have a role in appressorium formation and virulence of this pathogen on rice (77,78). In the mycoparasitice fungus, *Coniothyrium minitans*, disruption of a DNA helicase gene resulted in its altered morphology, reduced growth rate, and loss of mycoparasitism towards *Sclerotinia sclerotiorum* (79). Telomeric regions in several fungi have been reported to be highly dynamic and accommodate genes that confer an adaptive advantage in evading host recognition (80,81). Therefore, it is plausible that the identified variations in association with DNA helicases in sub-telomeric regions may have a role in transcription and epigenetic changes associated with aggressiveness in *A. rabiei*.

Many of the SNPs identified in the current study occur on genes encoding hypothetical proteins of unknown function with no predicted signal peptide or transmembrane domain. For example, ctg07_899731 and ctg21_487980 identified by pyseer and ctg13_5391 and ctg13_5645 identified through DAPC analyses and pairwise comparison, cause missense variations on genes coding for hypothetical proteins of unknown function. These highlight the limitation of the current reference genome of *A. rabiei*, which still lacks accurate functional annotation for many hypothetical predicted genes. Further studies are required to understand the potential role of these loci in aggressiveness of *A. rabiei* on chickpea.

Variations ctg07_1292465 and ctg20_254353 were found to cause missense mutations in genes coding for a hypothetical alkaline phosphatase and a hypothetical ubiquitin transferase, both of which have been associated with fungal pathogenesis on various pathosystems (82,83). Several variations, however, were identified in intergenic regions in the vicinity of coding sequences that code for hypothetical proteins with potential function in pathogenicity and aggressiveness. These include genes encoding a secreted carbohydrate-binding trehalase (EKO05_010756), a secreted protein with similarity to glycosylphosphatidylinositol (GPI)- anchored, serine-threonine rich proteins (EKO05_007578), and multiple hypothetical proteins with transmembrane domains and high similarity to sugar, oligopeptide, and copper transporters. Trehalases have an important role in the hydrolysis of external trehalose in fungi and bacteria, where they play an important role in pathogenicity, virulence, and stress resistance (74,84–86). Some trehalases may have other cellular functions, *e.g.*, a trehalase-encoding gene was primarily expressed in spores of *Plasmodiophora brassicae*, believed to release glucose as a source of energy during the germination process (87). GPI-anchored, serine-threonine rich proteins in *Candida albicans* have been reported as key effectors of fungal adherence to host cells (88). The role of transmembrane proteins in cellular transport, metabolism, signal transduction, conidial germination and aggressiveness of fungal plant pathogens is well documented (75,89–92).

Several variations associated with *A. rabiei* aggressiveness on chickpea varieties were found in proximity of loci coding for predicted apoplastic or cytoplasmic effectors on the reference genome (strain ArME14). Effectors are secreted proteins that facilitate infection through suppression of plant defense responses and alteration of host cell structure and function (93). Apoplastic effectors are secreted into the host extracellular space and can function in the apoplast or may be bound to the fungal cell wall (94,95). On the other hand, cytoplasmic effectors are secreted into the cytoplasm and may subsequently target specific plant cell compartments (60). One of the genes predicted to encode for an apoplastic effector (EKO05_005328) shows high similarity to fungal genes encoding rapid alkalinization factors (RALFs). Members of the RALF family are a family of conserved plant regulatory peptides, homologues of which have also been detected in plant pathogenic fungi with potential role in pathogenicity. For example, *Fusarium oxysporum* mutant strains lacking a functional RALF showed reduced virulence in tomato plants (96) and a RALF peptide has been suggested to be a minor contributor to *F. graminearum* virulence on wheat (97). Presence/absence of these effectors in different *A. rabiei* isolates and the potential for differential expression of these effectors in isolates with different levels of aggressiveness should be further investigated. Investigation of potential association of SNPs with recently functionally characterised *A. rabiei* effectors detected one modifier SNP located upstream of the gene encoding ArCRZ1 protein (Supplementary Table S8), which is a putative transcriptional regulator of virulence factors in *A. rabiei* (98).

Many genetic variations detected here were also located within transposable elements (TEs) and repetitive regions of the genome. Most of these repeat regions consisted of TEs, including the only SNP that was commonly associated with aggressiveness across all host chickpea cultivars, which occurred within a retrotransposon of the LTR/Gypsy family, known to be involved in rapid increases in fungal pathogenicity (99). Transposable elements are self-replicating segments in the genome, which are the target of epigenomic regulation that is also known to affect nearby genes. In other words, TEs have the ability to also alter gene regulation (100) and have been implicated in differential expression of effector loci and pathogenicity-related genes in plant pathogenic fungi (101,102).

Our study revealed that 22.5% of *A. rabiei* genome consists of AT-rich regions (secondary peak in Fig. 3), which are known to harbour repetitive regions and TEs. This is similar to the reported value of 21% reported for an earlier version of *A. rabiei* genome (31). This phenomenon in fungi is referred to as the ‘two-speed genome’ (103), where effectors with important role in virulence and aggressiveness of plant pathogens lie within rapidly evolving, AT-rich regions with highly repetitive content and increased TE activity. This will allow the effector loci within the fast-evolving, repetitive genome compartments to evolve through duplication, modification, and inactivation, or be lost entirely in a short period of time due to the activity of TEs. Therefore, although genetic variations detected in the repetitive regions of the genome were annotated as intergenic, these could potentially be located on effector loci, which have not been annotated due to their highly divergent sequences and low homology to known effectors. High levels of presence/absence polymorphisms of effectors have been reported within fungal plant pathogens such as *M. oryzae* (104) and *Blumeria graminis* (87). Although the percentage of AT-rich repetitive regions in *A. rabiei* is considerably lower than many other aggressive plant pathogens such as *Leptosphaeria maculans* (shown in the secondary peaks left of the vertical line in Fig. 3), our results show that high rate of variation in these regions may be a major driving force underlying the rapid evolution of aggressiveness in the *A. rabiei* population in Australia.

A commonly used approach for investigating genetic loci underlying adaptive mechanisms, which has been successfully applied to plant pathogens, is investigating signatures of positive selection. Two commonly used approaches to detect signatures of selection are the dN/dS ratio test and selective sweep scans (105). The dN/dS ratio test is more appropriate for macroevolutionary scenarios spanning speciation events (106) while sweep scans can be used to investigate recent evolution of populations undergoing strong positive selection similar to the situation for *A. rabiei* in Australia. However, detection of selection sweeps is reliant on presence of sexual reproduction in pathogen populations, where loci undergoing selective sweeps can be differentiated from neighbouring loci because of a significant reduction in sequence diversity, on which meiotic recombination has not yet had a chance to act to break the linkage of loci. Such analyses are, therefore, not appropriate for a highly clonal, asexually reproducing pathogen (107) like *A. rabiei* in Australia, with very low levels of genome wide polymorphism and a genome that is effectively entirely linked (Fig. S2).

This study presents the first step towards understanding the molecular mechanisms of aggressiveness in *A. rabiei* populations in Australia. Detection of multiple SNPs with incremental contribution towards aggressiveness of *A. rabiei* isolates is in line with the phenotypic observation demonstrating a quantitative variation in pathogenicity of *A. rabiei* isolates on different chickpea varieties. Differences in aggressiveness of pathogens has previously been suggested to be conditioned by minor gene-for-minor gene interactions and variation in aggressiveness traits defined as quantitative life-history traits, such as infection efficiency, latent period, sporulation rate, and lesion size (108).

The limited number of small variant (SNPs) differences between pairs of isolates with contrasting aggressiveness suggest that other forms of genetic modifications, including large structural variants such as accessory chromosomes, gene presence/absence, copy number variations and large indels, as well as epigenetic modifications may have an important role determining *A. rabiei* aggressiveness on chickpea. The short-read sequencing approach performed here proved cost-efficient to generate high coverage, high-quality, genome-wide SNP variants on a population-scale on a small genome such as *A. rabiei* and was effective to identify underlying population structure using a larger number of markers ever used. However, short-read data could not be used to identify larger structural and epigenetic variations. Therefore, our future efforts will focus on long-read sequencing of genetically close isolates with contrasting aggressiveness as well as epigenomic investigations to further unravel the underlying mechanisms of aggressiveness in *A. rabiei*. Identifying the genetic regions contributing to these traits will help with rapid and reliable detection of highly adapted and aggressive isolates.

## Acknowledgements

We would like to acknowledge the GRDC for funding this project and thank the wider Ascochyta blight research community in Australia, including state pathologists, growers, breeders, agronomists and research partners for kindly assisting in isolate collection and sharing resources and knowledge. This research was supported by the Griffith University Gowonda HPC Cluster and made use of the NeCTAR Research Cloud provided by QCIF (http://www.qcif.edu.au) for data analysis and visualisation. The NeCTAR Research Cloud is a collaborative Australian research platform supported by the National Collaborative Research Infrastructure Strategy (NCRIS) fund from the Australian Research Data Commons (ARDC).

