## Supplementary_File for "Population-level whole genome sequencing of *Ascochyta rabiei* identifies genomic loci associated with isolate aggressiveness"

**Table. S1.** List of *Ascochyta rabiei* isolates collected from 2013 to 2020 in Australia, which were used for whole genome sequencing and variant calling.

| **Isolate ID** | **Year** | **State** | **Host** | **Zone** | **Zone abbreviation** | **ICC3996** | **Genesis90** | **PBA HatTrick** | **PBA Seamer** |
| --- | --- | --- | --- | --- | --- | --- | --- | --- | --- |
| FT13092-2 | 2013 | SA | Genesis90 | SA Midnorth-Lower Yorke Eyre | SA1 | Moderate | High | High | unknown |
| FT13092-4 | 2013 | SA | Genesis90 | SA Midnorth-Lower Yorke Eyre | SA1 | Moderate | Moderate | High | unknown |
| FT13092-6 | 2013 | SA | Genesis90 | SA Midnorth-Lower Yorke Eyre | SA1 | Low | Moderate | High | unknown |
| 14DON003 | 2014 | VIC | Slasher | SA Vic Mallee | SA/Vic2 | Low | Low | High | unknown |
| TR6400 | 2014 | NSW | Hattrick | NSW NE/Qld SE | NSW/Qld1 | Low | Low | High | unknown |
| TR6408 | 2014 | NSW | Hattrick | NSW NE/Qld SE | NSW/Qld1 | Low | Low | Low | unknown |
| TR6417 | 2014 | NSW | Hattrick | NSW NE/Qld SE | NSW/Qld1 | Moderate | High | High | unknown |
| 15CUR001 | 2015 | VIC | Genesis90 | SA Vic Mallee | SA/Vic2 | Low | Moderate | High | unknown |
| 15CUR002 | 2015 | VIC | Genesis90 | SA Vic Mallee | SA/Vic2 | Moderate | High | High | unknown |
| 15CUR003 | 2015 | VIC | Genesis90 | SA Vic Mallee | SA/Vic2 | Low | Low | High | unknown |
| 15DON001 | 2015 | VIC | Genesis90 | SA Vic Mallee | SA/Vic2 | Low | Low | Low | unknown |
| 15DON007 | 2015 | VIC | Slasher | SA Vic Mallee | SA/Vic2 | Moderate | Moderate | High | unknown |
| FT15025 | 2015 | SA | Genesis90 | SA Midnorth-Lower Yorke Eyre | SA1 | Low | Moderate | High | unknown |
| FT15028 | 2015 | SA | Genesis90 | SA Midnorth-Lower Yorke Eyre | SA1 | Moderate | Moderate | High | unknown |
| FT15029 | 2015 | SA | Genesis90 | SA Midnorth-Lower Yorke Eyre | SA1 | Moderate | Moderate | High | unknown |
| FT15030 | 2015 | SA | Genesis90 | SA Midnorth-Lower Yorke Eyre | SA1 | Moderate | Moderate | High | unknown |
| 16CUR015 | 2016 | VIC | Genesis90 | SA Vic Mallee | SA/Vic2 | Low | Moderate | High | Low |
| 16CUR017 | 2016 | VIC | Genesis90 | SA Vic Mallee | SA/Vic2 | Moderate | Moderate | High | Low |
| 16CUR018 | 2016 | VIC | Genesis90 | SA Vic Mallee | SA/Vic2 | Moderate | High | High | Moderate |
| 16CUR019 | 2016 | VIC | Genesis90 | SA Vic Mallee | SA/Vic2 | Moderate | Moderate | High | Low |
| 16RUP012 | 2016 | VIC | Genesis90 | SA Vic Bordertown-Wimmera | SA/Vic1 | Low | Low | High | Low |
| 16RUP013 | 2016 | VIC | Genesis90 | SA Vic Bordertown-Wimmera | SA/Vic1 | Low | Low | High | Low |
| F16083-1 | 2016 | SA | Genesis90 | SA Midnorth-Lower Yorke Eyre | SA1 | Moderate | Moderate | High | Moderate |
| F16253-1 | 2016 | SA | Genesis90 | SA Midnorth-Lower Yorke Eyre | SA1 | Moderate | Moderate | High | Low |
| TR8102 | 2016 | NSW | Hattrick | NSW NW/Qld SW | NSW/Qld2 | Low | Moderate | High | Low |
| TR8105 | 2016 | NSW | Hattrick | NSW NW/Qld SW | NSW/Qld2 | Low | Low | High | Low |
| 17CUR007 | 2017 | VIC | Genesis90 | SA Vic Mallee | SA/Vic2 | Moderate | Moderate | High | Moderate |
| F17067-1 | 2017 | SA | Genesis90 | SA Vic Bordertown-Wimmera | SA/Vic1 | Low | Low | Low | Low |
| F17076-2 | 2017 | NSW | Genesis90 | NSW Central | NSW1 | Moderate | High | Moderate | Low |
| F17175-1 | 2017 | VIC | Genesis90 | SA Vic Bordertown-Wimmera | SA/Vic1 | Low | Low | Low | Low |
| F17191-1 | 2017 | SA | Genesis90 | SA Midnorth-Lower Yorke Eyre | SA1 | High | High | High | High |
| TR9529 | 2017 | QLD | Seamer | NSW NE/Qld SE | NSW/Qld1 | High | High | High | High |
| TR9538 | 2017 | QLD | Seamer | unknown | unknown | Low | Low | Low | Low |
| TR9543 | 2017 | QLD | Seamer | unknown | unknown | Low | High | Moderate | Low |
| TR9544 | 2017 | QLD | Seamer | unknown | unknown | Low | Low | Low | Low |
| TR9568 | 2017 | NSW | Seamer | NSW NE/Qld SE | NSW/Qld1 | Moderate | Moderate | High | Moderate |
| TR9571 | 2017 | NSW | Seamer | NSW NE/Qld SE | NSW/Qld1 | High | High | High | High |
| AR0002 | 2020 | NSW | Drummond | NSW NE/Qld SE | NSW/Qld1 | Moderate | Moderate | Moderate | High |
| AR0003 | 2020 | NSW | Drummond | NSW NE/Qld SE | NSW/Qld1 | Moderate | High | High | Moderate |
| AR0005 | 2020 | NSW | Kimberly | NSW NE/Qld SE | NSW/Qld1 | High | High | High | High |
| AR0007 | 2020 | NSW | Kimberly | NSW NE/Qld SE | NSW/Qld1 | Moderate | Moderate | High | Moderate |
| AR0009 | 2020 | NSW | ICCU96329 | NSW NE/Qld SE | NSW/Qld1 | High | High | High | High |
| AR0013 | 2020 | NSW | ACPFGRIL84 | NSW NE/Qld SE | NSW/Qld1 | High | High | High | High |
| AR0015 | 2020 | NSW | ILC05702 | NSW NE/Qld SE | NSW/Qld1 | High | High | High | High |
| AR0016 | 2020 | NSW | ICCU96329 | NSW NE/Qld SE | NSW/Qld1 | High | High | High | High |
| AR0018 | 2020 | NSW | 2-15 | NSW NE/Qld SE | NSW/Qld1 | High | High | High | High |
| AR0020 | 2020 | NSW | 2-164 | NSW NE/Qld SE | NSW/Qld1 | High | High | High | High |
| AR0021 | 2020 | NSW | ILC05658 | NSW NE/Qld SE | NSW/Qld1 | Moderate | Moderate | High | High |
| AR0022 | 2020 | NSW | ICCU96329 | NSW NE/Qld SE | NSW/Qld1 | Moderate | High | Moderate | High |
| AR0023 | 2020 | NSW | 2-216 | NSW NE/Qld SE | NSW/Qld1 | High | High | High | High |
| AR0024 | 2020 | NSW | 2-15 | NSW NE/Qld SE | NSW/Qld1 | Moderate | High | High | High |
| AR0025 | 2020 | NSW | 1-131xSel2 | NSW NE/Qld SE | NSW/Qld1 | Moderate | High | High | High |
| AR0026 | 2020 | NSW | BG112 | NSW NE/Qld SE | NSW/Qld1 | Low | Moderate | Moderate | Low |
| AR0027 | 2020 | NSW | 1-131xSel2 | NSW NE/Qld SE | NSW/Qld1 | High | High | High | High |
| AR0028 | 2020 | NSW | ICCU96329 | NSW NE/Qld SE | NSW/Qld1 | Low | Moderate | Low | Moderate |
| AR0029 | 2020 | NSW | ILC05600 | NSW NE/Qld SE | NSW/Qld1 | Low | Low | Low | Moderate |
| AR0031 | 2020 | NSW | unknown | unknown | unknown | Low | High | Moderate | Moderate |
| AR0032 | 2020 | NSW | CBA 2055 | NSW Vic Slopes | NSW/Vic | Low | High | Moderate | Moderate |
| AR0033 | 2020 | NSW | Slasher | NSW Vic Slopes | NSW/Vic | High | High | High | High |
| AR0034 | 2020 | NSW | Kyabra | NSW Vic Slopes | NSW/Vic | Low | High | High | Moderate |
| AR0035 | 2020 | NSW | Genesis90 | NSW Vic Slopes | NSW/Vic | Moderate | Low | Moderate | Low |
| AR0036 | 2020 | NSW | Seamer | NSW Vic Slopes | NSW/Vic | Moderate | Low | Moderate | Low |
| AR0037 | 2020 | NSW | Boundary | NSW Vic Slopes | NSW/Vic | Low | Low | Low | Moderate |
| AR0038 | 2020 | NSW | Kyabra | NSW Vic Slopes | NSW/Vic | Low | Moderate | Moderate | Low |
| AR0039 | 2020 | NSW | Royal | NSW Vic Slopes | NSW/Vic | Low | Low | Low | Moderate |
| AR0040 | 2020 | NSW | Royal | NSW Vic Slopes | NSW/Vic | Low | Moderate | Moderate | Low |
| AR0042 | 2020 | NSW | Hattrick | NSW Vic Slopes | NSW/Vic | Moderate | High | Low | Moderate |
| AR0044 | 2020 | NSW | Hattrick | NSW Vic Slopes | NSW/Vic | Moderate | Moderate | High | Moderate |
| AR0048 | 2020 | NSW | Hattrick | NSW Vic Slopes | NSW/Vic | Low | High | Low | Moderate |
| AR0050 | 2020 | NSW | Hattrick | NSW Vic Slopes | NSW/Vic | Low | High | High | Moderate |
| AR0052 | 2020 | NSW | Hattrick | NSW Vic Slopes | NSW/Vic | Low | Moderate | High | Moderate |
| AR0055 | 2020 | NSW | Hattrick | NSW Vic Slopes | NSW/Vic | Moderate | High | High | High |
| AR0056 | 2020 | NSW | Hattrick | NSW Vic Slopes | NSW/Vic | Low | High | High | High |
| AR0060 | 2020 | NSW | Hattrick | NSW Vic Slopes | NSW/Vic | Low | Low | Low | Low |
| AR0062 | 2020 | NSW | Hattrick | NSW Vic Slopes | NSW/Vic | Moderate | Low | Moderate | High |
| AR0063 | 2020 | NSW | Hattrick | NSW Vic Slopes | NSW/Vic | Low | Low | Low | High |
| AR0064 | 2020 | WA | Amber | WA Sandplain | WA2 | High | High | High | Moderate |
| AR0065 | 2020 | WA | Amber | WA Sandplain | WA2 | High | High | High | Moderate |
| AR0066 | 2020 | WA | Amber | WA Sandplain | WA2 | High | High | High | High |
| AR0067 | 2020 | WA | Amber | WA Sandplain | WA2 | Moderate | Moderate | High | High |
| AR0068 | 2020 | WA | Amber | WA Sandplain | WA2 | Moderate | High | High | Moderate |
| AR0069 | 2020 | WA | Amber | WA Sandplain | WA2 | Moderate | High | High | High |
| AR0070 | 2020 | WA | Amber | WA Sandplain | WA2 | Moderate | High | High | High |
| AR0071 | 2020 | WA | Amber | WA Sandplain | WA2 | Moderate | High | High | High |
| AR0072 | 2020 | WA | Amber | WA Sandplain | WA2 | Moderate | High | Moderate | High |
| AR0073 | 2020 | WA | Amber | WA Sandplain | WA2 | Low | Moderate | Low | Low |
| AR0074 | 2020 | WA | Amber | WA Sandplain | WA2 | Low | High | High | High |
| AR0075 | 2020 | WA | Amber | WA Sandplain | WA2 | Moderate | High | High | High |
| AR0076 | 2020 | WA | Amber | WA Sandplain | WA2 | Low | High | High | High |
| AR0078 | 2020 | WA | Amber | WA Sandplain | WA2 | Low | High | High | High |
| AR0080 | 2020 | WA | Amber | WA Sandplain | WA2 | High | High | High | High |
| AR0081 | 2020 | WA | Amber | WA Sandplain | WA2 | Moderate | High | High | High |
| AR0083 | 2020 | WA | Amber | WA Sandplain | WA2 | Moderate | High | High | High |
| AR0087 | 2020 | WA | Amber | WA Sandplain | WA2 | Low | High | High | High |
| AR0088 | 2020 | NSW | Hattrick | NSW Vic Slopes | NSW/Vic | Moderate | Moderate | High | High |
| AR0091 | 2020 | NSW | Hattrick | NSW Vic Slopes | NSW/Vic | Low | Low | High | High |
| AR0093 | 2020 | NSW | Hattrick | NSW Vic Slopes | NSW/Vic | Low | Moderate | Moderate | High |
| AR0098 | 2020 | NSW | Hattrick | NSW Vic Slopes | NSW/Vic | Low | Low | Low | Moderate |
| AR0103 | 2020 | NSW | Hattrick | NSW Vic Slopes | NSW/Vic | Low | Low | Moderate | High |
| AR0105 | 2020 | NSW | CICA1521 | NSW NE/Qld SE | NSW/Qld1 | Low | Low | Moderate | High |
| AR0108 | 2020 | NSW | Hattrick | NSW NE/Qld SE | NSW/Qld1 | Moderate | Low | High | High |
| AR0112 | 2020 | QLD | Jimbour | Qld Central | Qld | Low | Low | Low | Low |
| AR0114 | 2020 | QLD | Jimbour | Qld Central | Qld | Low | Low | Low | Low |
| AR0115 | 2020 | QLD | Jimbour | Qld Central | Qld | Low | Low | Low | Low |
| AR0116 | 2020 | QLD | Jimbour | Qld Central | Qld | Low | Low | Low | Low |
| AR0117 | 2020 | NSW | Seamer | NSW NE/Qld SE | NSW/Qld1 | Low | Low | Low | Low |
| AR0119 | 2020 | NSW | Seamer | NSW NE/Qld SE | NSW/Qld1 | Low | Low | Low | Low |
| AR0122 | 2020 | NSW | Boundary | NSW NE/Qld SE | NSW/Qld1 | High | Low | High | High |
| AR0123 | 2020 | NSW | Boundary | NSW NE/Qld SE | NSW/Qld1 | Low | Moderate | High | High |
| AR0127 | 2020 | NSW | Boundary | NSW NE/Qld SE | NSW/Qld1 | Low | Moderate | Moderate | Low |
| AR0128 | 2020 | NSW | Boundary | NSW NE/Qld SE | NSW/Qld1 | High | High | High | High |
| AR0132 | 2020 | NSW | Hattrick | NSW NE/Qld SE | NSW/Qld1 | Moderate | Low | Low | High |
| AR0133 | 2020 | NSW | Hattrick | NSW NE/Qld SE | NSW/Qld1 | Low | Low | Moderate | Moderate |
| AR0134 | 2020 | NSW | Hattrick | NSW NE/Qld SE | NSW/Qld1 | Low | Moderate | Low | High |
| AR0135 | 2020 | NSW | Hattrick | NSW NE/Qld SE | NSW/Qld1 | Low | Moderate | Moderate | Moderate |
| AR0150 | 2020 | SA | unknown | SA Midnorth-Lower Yorke Eyre | SA1 | Moderate | Moderate | High | High |
| AR0151 | 2020 | SA | unknown | SA Midnorth-Lower Yorke Eyre | SA1 | Moderate | Low | Moderate | Moderate |
| AR0152 | 2020 | SA | unknown | SA Midnorth-Lower Yorke Eyre | SA1 | Moderate | Low | Moderate | Moderate |
| AR0153 | 2020 | SA | Genesis90 | SA Midnorth-Lower Yorke Eyre | SA1 | Low | Low | Moderate | Moderate |
| AR0154 | 2020 | SA | Genesis90 | SA Midnorth-Lower Yorke Eyre | SA1 | Moderate | Low | Moderate | Low |
| AR0155 | 2020 | SA | Genesis90 | SA Midnorth-Lower Yorke Eyre | SA1 | Moderate | Low | High | High |
| AR0156 | 2020 | SA | Monarch | SA Midnorth-Lower Yorke Eyre | SA1 | Low | High | Moderate | High |
| AR0157 | 2020 | SA | Monarch | SA Midnorth-Lower Yorke Eyre | SA1 | Moderate | Moderate | Moderate | Moderate |
| AR0158 | 2020 | SA | Monarch | SA Midnorth-Lower Yorke Eyre | SA1 | Moderate | Low | High | Moderate |
| AR0159 | 2020 | SA | Monarch | SA Midnorth-Lower Yorke Eyre | SA1 | Low | Low | Low | Low |
| AR0160 | 2020 | SA | Monarch | SA Midnorth-Lower Yorke Eyre | SA1 | Moderate | Low | Moderate | Moderate |
| AR0161 | 2020 | SA | Monarch | SA Midnorth-Lower Yorke Eyre | SA1 | Low | Low | Low | Low |
| AR0162 | 2020 | SA | Genesis90 | SA Midnorth-Lower Yorke Eyre | SA1 | Low | Moderate | High | High |
| AR0163 | 2020 | SA | Genesis90 | SA Midnorth-Lower Yorke Eyre | SA1 | Moderate | Low | Moderate | Moderate |
| AR0164 | 2020 | SA | Genesis90 | SA Midnorth-Lower Yorke Eyre | SA1 | Low | Moderate | Low | Low |
| AR0165 | 2020 | SA | Monarch | SA Midnorth-Lower Yorke Eyre | SA1 | Low | Low | Moderate | Moderate |
| AR0166 | 2020 | SA | Monarch | SA Midnorth-Lower Yorke Eyre | SA1 | Moderate | Low | High | Moderate |
| AR0167 | 2020 | SA | Monarch | SA Midnorth-Lower Yorke Eyre | SA1 | Moderate | Low | unknown | Low |
| AR0168 | 2020 | SA | Monarch | SA Midnorth-Lower Yorke Eyre | SA1 | Low | Low | Moderate | Moderate |
| AR0169 | 2020 | SA | Monarch | SA Midnorth-Lower Yorke Eyre | SA1 | Moderate | Low | Low | Low |
| AR0170 | 2020 | SA | Monarch | SA Midnorth-Lower Yorke Eyre | SA1 | Moderate | High | High | High |
| AR0175 | 2020 | WA | Neelam | WA Central | WA1 | Moderate | Low | Moderate | Moderate |
| AR0176 | 2020 | WA | Neelam | WA Central | WA1 | Moderate | Low | Low | Low |
| AR0177 | 2020 | WA | Neelam | WA Central | WA1 | Low | Low | Low | Low |
| AR0178 | 2020 | WA | Neelam | WA Central | WA1 | Low | Low | Low | Low |
| AR0179 | 2020 | WA | Neelam | WA Central | WA1 | High | Moderate | Low | Low |
| AR0180 | 2020 | WA | Neelam | WA Central | WA1 | Low | Low | Low | Low |
| AR0181 | 2020 | WA | Neelam | WA Central | WA1 | Low | Low | Low | Moderate |
| AR0182 | 2020 | WA | Neelam | WA Central | WA1 | Low | Low | Low | Low |
| AR0183 | 2020 | WA | Neelam | WA Central | WA1 | Low | Low | Low | Low |
| AR0184 | 2020 | WA | Neelam | WA Central | WA1 | Low | Low | Low | Low |
| AR0185 | 2020 | WA | Neelam | WA Central | WA1 | Low | Moderate | Moderate | Low |
| AR0186 | 2020 | WA | Neelam | WA Central | WA1 | Moderate | Low | Moderate | Low |
| AR0187 | 2020 | WA | Neelam | WA Central | WA1 | Low | Low | Moderate | Low |
| AR0188 | 2020 | WA | Neelam | WA Central | WA1 | Moderate | Low | Moderate | Low |
| AR0189 | 2020 | WA | Neelam | WA Central | WA1 | Low | Low | Low | Low |
| AR0190 | 2020 | WA | Neelam | WA Central | WA1 | Low | Moderate | Moderate | Moderate |
| AR0191 | 2020 | WA | Neelam | WA Central | WA1 | Low | Low | Moderate | Low |
| AR0192 | 2020 | WA | Neelam | WA Central | WA1 | Moderate | High | High | High |
| AR0193 | 2020 | WA | Neelam | WA Central | WA1 | Low | Moderate | Moderate | Low |
| AR0194 | 2020 | WA | Neelam | WA Central | WA1 | Low | Moderate | Low | Low |
| AR0195 | 2020 | WA | Neelam | WA Central | WA1 | Low | Low | Low | Low |
| AR0196 | 2020 | WA | Neelam | WA Central | WA1 | Low | Low | Low | Low |
| AR0197 | 2020 | WA | Neelam | WA Central | WA1 | Moderate | Low | Moderate | Low |
| AR0198 | 2020 | WA | Neelam | WA Central | WA1 | Moderate | Low | Low | Low |
| AR0199 | 2020 | WA | Neelam | WA Central | WA1 | Low | Low | Low | Low |
| AR0205 | 2020 | NSW | Hattrick | NSW NE/Qld SE | NSW/Qld1 | Moderate | Moderate | High | High |
| AR0206 | 2020 | NSW | Hattrick | NSW NE/Qld SE | NSW/Qld1 | Moderate | Moderate | High | High |
| AR0210 | 2020 | NSW | Hattrick | NSW NW/Qld SW | NSW/Qld2 | High | Moderate | High | High |
| AR0211 | 2020 | NSW | Hattrick | NSW NW/Qld SW | NSW/Qld2 | Low | Low | Low | High |
| AR0212 | 2020 | NSW | Hattrick | NSW NW/Qld SW | NSW/Qld2 | Moderate | High | High | High |
| AR0215 | 2020 | NSW | Hattrick | NSW NW/Qld SW | NSW/Qld2 | Low | Low | Moderate | Moderate |
| AR0216 | 2020 | NSW | Hattrick | NSW Central | NSW1 | Low | Moderate | Moderate | High |
| AR0217 | 2020 | NSW | Hattrick | NSW Central | NSW1 | Low | High | Moderate | High |
| AR0218 | 2020 | NSW | Hattrick | NSW Central | NSW1 | Low | Moderate | High | High |
| AR0219 | 2020 | NSW | Hattrick | NSW Central | NSW1 | Low | Low | Low | Low |
| AR0220 | 2020 | NSW | Hattrick | NSW Central | NSW1 | Low | Low | Moderate | Moderate |
| AR0221 | 2020 | NSW | Seamer | NSW NW/Qld SW | NSW/Qld2 | Moderate | Moderate | High | High |
| AR0222 | 2020 | NSW | Seamer | NSW NW/Qld SW | NSW/Qld2 | Low | Low | Low | Moderate |
| AR0223 | 2020 | NSW | Seamer | NSW NW/Qld SW | NSW/Qld2 | Low | High | High | High |
| AR0225 | 2020 | NSW | Seamer | NSW NW/Qld SW | NSW/Qld2 | Low | Low | Moderate | Low |
| AR0226 | 2020 | NSW | Seamer | NSW NW/Qld SW | NSW/Qld2 | Moderate | High | High | High |
| AR0227 | 2020 | NSW | Seamer | NSW NW/Qld SW | NSW/Qld2 | Low | High | Moderate | Low |
| AR0228 | 2020 | NSW | Seamer | NSW NW/Qld SW | NSW/Qld2 | Low | Low | Low | Low |
| AR0230 | 2020 | NSW | Seamer | NSW NW/Qld SW | NSW/Qld2 | Low | Low | Low | Moderate |
| AR0231 | 2020 | NSW | Hattrick | NSW NW/Qld SW | NSW/Qld2 | High | High | High | High |
| AR0232 | 2020 | NSW | Hattrick | NSW NW/Qld SW | NSW/Qld2 | Low | High | High | High |
| AR0235 | 2020 | NSW | Hattrick | NSW NW/Qld SW | NSW/Qld2 | Moderate | High | High | High |
| AR0236 | 2020 | NSW | Hattrick | NSW NW/Qld SW | NSW/Qld2 | Low | High | High | High |
| AR0237 | 2020 | NSW | Hattrick | NSW NW/Qld SW | NSW/Qld2 | Moderate | High | High | High |
| AR0240 | 2020 | NSW | Hattrick | NSW NW/Qld SW | NSW/Qld2 | unknown | unknown | unknown | unknown |
| AR0241 | 2020 | NSW | Hattrick | NSW NW/Qld SW | NSW/Qld2 | Low | Low | High | Moderate |
| AR0242 | 2020 | NSW | Hattrick | NSW NW/Qld SW | NSW/Qld2 | Low | Low | High | Moderate |
| AR0246 | 2020 | NSW | Seamer | NSW Vic Slopes | NSW/Vic | Moderate | High | High | High |
| AR0247 | 2020 | NSW | Seamer | NSW Vic Slopes | NSW/Vic | Low | High | High | High |
| AR0248 | 2020 | NSW | Seamer | NSW Vic Slopes | NSW/Vic | Low | High | High | Moderate |
| AR0251 | 2020 | NSW | Hattrick | NSW Vic Slopes | NSW/Vic | Moderate | Moderate | High | Moderate |
| AR0256 | 2020 | NSW | Hattrick | NSW Vic Slopes | NSW/Vic | Moderate | High | High | High |
| AR0257 | 2020 | NSW | Hattrick | NSW Vic Slopes | NSW/Vic | Low | Low | Moderate | Moderate |
| AR0261 | 2020 | NSW | unknown | NSW Central | NSW1 | Low | Moderate | High | High |
| AR0262 | 2020 | NSW | unknown | NSW Central | NSW1 | Low | Moderate | Moderate | High |
| AR0263 | 2020 | NSW | unknown | NSW Central | NSW1 | Low | Moderate | High | High |
| AR0264 | 2020 | NSW | unknown | NSW Central | NSW1 | Low | Moderate | High | High |
| AR0265 | 2020 | NSW | unknown | NSW Central | NSW1 | Low | Moderate | High | Moderate |
| AR0266 | 2020 | WA | Amber | WA Sandplain | WA2 | Moderate | Moderate | High | High |
| AR0273 | 2020 | WA | Amber | WA Sandplain | WA2 | Moderate | High | High | High |
| AR0278 | 2020 | NSW | Hattrick | NSW Vic Slopes | NSW/Vic | Low | Moderate | High | High |
| AR0279 | 2020 | NSW | Hattrick | NSW Vic Slopes | NSW/Vic | Moderate | High | Moderate | Moderate |
| AR0280 | 2020 | NSW | Hattrick | NSW Vic Slopes | NSW/Vic | Moderate | Moderate | High | High |
| AR0285 | 2020 | NSW | Hattrick | NSW Vic Slopes | NSW/Vic | Moderate | High | High | High |
| AR0286 | 2020 | NSW | Hattrick | NSW Vic Slopes | NSW/Vic | Moderate | Moderate | Moderate | Moderate |
| AR0289 | 2020 | NSW | Hattrick | NSW Vic Slopes | NSW/Vic | Low | High | High | High |
| AR0290 | 2020 | NSW | Hattrick | NSW Vic Slopes | NSW/Vic | Low | Moderate | High | High |
| AR0293 | 2020 | VIC | Striker | SA Vic Bordertown-Wimmera | SA/Vic1 | Low | Low | High | Moderate |
| AR0294 | 2020 | VIC | Slasher | SA Vic Bordertown-Wimmera | SA/Vic1 | Low | High | Moderate | High |
| AR0295 | 2020 | VIC | Royal | SA Vic Bordertown-Wimmera | SA/Vic1 | Low | Moderate | Moderate | Moderate |
| AR0296 | 2020 | VIC | Kalkee | SA Vic Bordertown-Wimmera | SA/Vic1 | Low | Low | High | High |
| AR0297 | 2020 | VIC | Monarch | SA Vic Bordertown-Wimmera | SA/Vic1 | Low | High | High | High |
| AR0298 | 2020 | VIC | Genesis90 | SA Vic Bordertown-Wimmera | SA/Vic1 | Moderate | Low | High | High |
| AR0299 | 2020 | SA | Striker | SA Vic Mallee | SA/Vic2 | High | Moderate | High | Low |
| AR0300 | 2020 | VIC | Striker | SA Vic Bordertown-Wimmera | SA/Vic1 | Moderate | High | High | High |
| AR0301 | 2020 | VIC | Genesis90 | SA Vic Bordertown-Wimmera | SA/Vic1 | Moderate | High | High | High |
| AR0302 | 2020 | VIC | Genesis90 | SA Vic Mallee | SA/Vic2 | Low | Low | High | High |
| AR0303 | 2020 | VIC | Genesis90 | SA Vic Mallee | SA/Vic2 | Low | High | High | Low |
| AR0304 | 2020 | VIC | Genesis90 | SA Vic Bordertown-Wimmera | SA/Vic1 | Moderate | High | High | Moderate |
| AR0312 | 2020 | WA | Striker | WA Central | WA1 | Low | Moderate | Moderate | Moderate |
| AR0313 | 2020 | WA | Striker | WA Central | WA1 | Low | Moderate | Moderate | Moderate |
| AR0314 | 2020 | WA | Striker | WA Central | WA1 | Moderate | High | Moderate | Low |
| AR0315 | 2020 | WA | Striker | WA Central | WA1 | Moderate | High | High | High |
| AR0316 | 2020 | WA | Striker | WA Central | WA1 | Low | Moderate | High | High |
| AR0317 | 2020 | WA | Striker | WA Central | WA1 | Low | Low | High | High |
| AR0318 | 2020 | WA | Striker | WA Central | WA1 | High | High | High | High |
| AR0319 | 2020 | WA | Striker | WA Central | WA1 | Moderate | Low | High | Moderate |
| AR0320 | 2020 | WA | Striker | WA Central | WA1 | Moderate | High | High | High |
| AR0321 | 2020 | WA | Striker | WA Central | WA1 | Moderate | High | High | High |

**Table S2**. Summary of analysis of molecular variance (AMOVA) results for 230 *Ascochyta rabiei* isolates collected in 2013, 2014, 2015, 2016, 2017, and 2020 from different States and locations in Australia.

| **Source** | **d.f.^a^** | **SS^b^** | **MS^c^** | **variance** | **proportion of variation** | ***P* value** |
| --- | --- | --- | --- | --- | --- | --- |
| **Among years** | 5 | 2,398.63 | 479.72 | 2.37 | 1.0% | 0.322 |
| **Among States within years** | 11 | 7,7904.73 | 718.61 | 30.34 | 13.0% | 0.001 |
| **Among locations within States** | 34 | 12,111.780 | 367.02 | 46.94 | 21.0% | 0.001 |
| **Within samples** | 179 | 25,472.79 | 143.10 | 143.11 | 65.0% | 0.001 |
| **Total** | 229 | 47,887.94 | 210.96 | 222.76 | 100.0% | - |

^a^Degree of freedom

^b^Sum of squared observations

^c^Mean of squared observations

**Table S3.** Summary of analysis of molecular variance (AMOVA) results for 230 *Ascochyta rabiei* isolates collected in 2013, 2014, 2015, 2016, 2017, and 2020 from different agroecological zones and locations in Australia.

| **Source** | **d.f.^a^** | **SS^b^** | **MS^c^** | **variance** | **proportion of variation** | ***P* value** |
| --- | --- | --- | --- | --- | --- | --- |
| **Among years** | 5 | 2,398.63 | 382.84 | 7.34 | 3.0% | 0.143 |
| **Among zones within years** | 20 | 13,367.88 | 601.58 | 40.43 | 17.5% | 0.001 |
| **Among locations within Zone** | 25 | 6,648.63 | 235.76 | 30.59 | 14.0% | 0.001 |
| **Within samples** | 179 | 25,472.80 | 127.66 | 143.11 | 65.5% | 0.001 |
| **Total** | 229 | 47,887.94 | 186.42 | 221.47 | 100.0% | - |

^a^Degree of freedom

^b^Sum of squared observations

^c^Mean of squared observations

**Table S4.** Indices of genetic diversity of the Australian *Ascochyta rabiei* populations defined as groups of isolates collected from agroecological zones.

| **Population**^a^ | **N**^b^ | **MLL**^c^ | **eMLL**^d^ | **SE**^e^ | **λ**^f^ | **E5**^g^ | **CF**^h^ | **Hexp**^i^ |
| --- | --- | --- | --- | --- | --- | --- | --- | --- |
| NSW1 | 11 | 10 | 9.18 | 3.86e-01 | 0.98 | 0.95 | 9% | 0.00 |
| NSW/Qld1 | 39 | 32 | 9.48 | 6.67e-01 | 0.98 | 0.89 | 17% | 0.00 |
| NSW/Qld2 | 22 | 14 | 8.09 | 9.65e-01 | 0.95 | 0.88 | 36% | 0.00 |
| NSW/Vic | 37 | 26 | 9.05 | 8.41e-01 | 0.97 | 0.88 | 30% | 0.00 |
| SA1 | 31 | 29 | 9.81 | 4.12e-01 | 0.99 | 0.96 | 6% | 0.00 |
| SA/Vic1 | 13 | 13 | - | - | - | - | - | - |
| SA/Vic2 | 14 | 14 | - | - | - | - | - | - |
| Qld | 4 | 4 | - | - | - | - | - | - |
| WA1 | 35 | 23 | 8.69 | 9.58e-01 | 0.96 | 0.82 | 34% | 0.00 |
| WA2 | 20 | 18 | 9.53 | 5.80e-01 | 0.95 | 0.95 | 10% | 0.00 |

^a^ Agroecological zones include NSW1: NSW Central, NSW/Qld1: NSW NE/Qld SE, NSW/Qld2: NSW NW/Qld SW, NSW/Vic: NSW Victoria slopes, Qld: Queensland Central, SA1: SA Midnorth-Lower Yorke Eyre, SA/Vic1: SA Vic Bordertown-Wimmera, SA/Vic2: SA Vic Mallee, WA1: WA Central, WA2: WA Sandplain (Fig. 1).

^b^ N = population size.

^c^ MLL = Number of multi-locus lineages after contracting the data set using the *cutoff_predictor* and *mlg.filter* functions in *poppr* (1,2).

^d^ eMLL = expected number of MLLs at the lowest common sample size to allow comparison between populations.

^e^ SE = The standard error for the rarefaction analysis for eMLLs.

^f^ λ = defined here as unbiased Simpson's complement index of genotypic diversity, defined as the probability that two genotypes randomly chosen from the population are different (3), corrected for sample size.

^g^ E5 = Evenness as a measure of the distribution of MLL abundances, where in a population with equally abundant MLLs, E5 = 1, and in a population dominated by a single MLL, E5=0.

^h^ CF = Clonal fraction; (N-number of MLLs)/N, where N is total number of isolates.

^i^ H_exp_ = Nei's gene diversity (expected heterozygosity) (4).

**Table S5.** Genetic variations with significant impact on aggressiveness of *Ascochyta rabiei* on different chickpea cultivars based on pyseer (5) results. Variant type and impact based on SNPEff (6) and putative function of the adjacent loci based on BLASTp results are given. Predector score is provided with EffectorP 3.0 score in parentheses. Non-effector indicates a negative Predector score and > 0.5 non-effector score from EffectorP 3.0.

| **variation** | **Variation also detected in DAPC** | **DNA region detected in DAPC** | ***p* value** | | | | **Variant type and impact** | **Putative protein function** | **Predector Score (EffectorP 3.0 score)** | **Repeat family** |
| --- | --- | --- | --- | --- | --- | --- | --- | --- | --- | --- |
|  |  |  | **Genesis090** | **ICC3996** | **PBA HatTrick** | **PBA Seamer** |  |  |  |  |
| ctg01_1654657 | Yes | Yes, region 3 | - | - | - | 0.0273 | intergenic_region, modifier, EKO05_000484-EKO05_000485 | EKO05_000484: hypothetical protein with high similarity to fungal GTPases, EKO05_000485 hypothetical DNA binding protein with a transmembrane domain | non-effectors | LTR/Copia |
| ctg01_1665154 | Yes | Yes, region 3 | - | - | - | 0.0273 | same as ctg01_1654657 |  |  |  |
| ctg01_2020023 | - | - | - | - | - | 0.0321 | synonymous_variant, low impact, EKO05_000605 | hypothetical protein of unknown function with similarity to fungal peptidases | -3.346 (0.716: Cytoplasmic effector but no signal peptide detected) | - |
| ctg01_3303380 | - | - |  | - | - | 0.0321 | intergenic_region, modifier, EKO05_001003-EKO05_001004 | EKO05_001003: **hypothetical apoplastic effector**, EKO05_001004: hypothetical protein of unknown function | EKO05_001003: 1.475 (0.808: Apoplastic effector), EKO05_001004: non-effector | LTR/Gypsy |
| ctg02_2690139 | Yes | Yes, region 8 | - | - | - | 0.0273 | missense_variant, moderate impact, EKO05_001768 | hypothetical protein with similarity to fungal transcription factors with a zinc-finger domain | non-effector | - |
| ctg02_325563 | - | Yes, region 6 | - | - | - | 0.0347 | intergenic_region, modifier, EKO05_001071-EKO05_001072 | EKO05_001071: hypothetical HET domain containing protein with transmembrane domain, EKO05_001072: Phosphotransferase | non-effectors | LTR/Gypsy |
| ctg02_364191 | Yes | Yes, region 6 | - | - | - | 0.0273 | same as ctg_02_325563 |  |  |  |
| ctg02_83011 | - | - | - | - | - | 0.0347 | intergenic_region, modifier, EKO05_001011-EKO05_001012 | EKO05_001011**: hypothetical Apoplastic/cytoplasmic effector**, EKO05_001012: **hypothetical effector with similarity to pep1 protein** | EKO05_001011: 1.873 (0.832, Apoplastic /cytoplasmic effector, signal peptide detected), EKO05_001012: 0.841 (0.579, Apoplastic effector, signal peptide detected) | DNA/TcMar-Fot1 |
| ctg03_1415178 | Yes | Yes, region 12 | - | - | - | 0.0273 | intergenic_region, modifier, EKO05_002061-EKO05_002062 | EKO05_002061: hypothetical protein of unknown function, EKO05_002062: **hypothetical apoplastic effector** | EKO05_002061: non-effector, EKO05_002062: 0.676 (0.709, Apoplastic effector) | LTR/Gypsy |
| ctg03_1428702 | - | Yes, region 12 | - | - | - | 0.0347 | same as ctg_03_1415178 |  |  |  |
| ctg03_2025011 | - | Yes, region 14 | - | - | 0.0036 | 0.0339 | intergenic_region, modifier, EKO05_002252-EKO05_002253 | EKO05_002252-EKO05: hypothetical protein of unknown function, EKO05_002253: hypothetical protein with similarity to oxidoreductases | non-effectors | LTR/Gypsy |
| ctg03_357028 | - | - | - | - | 0.0489 | 0.0143 | intergenic_region, modifier, EKO05_001820, EKO05_001821 | EKO05_001820: hypothetical protein of unknown function with a transmembrane domain, EKO05_001821: hypothetical protein of unknown function | non-effectors | LTR/Copia |
| ctg03_479997 | - | - | - | - | - | 0.0321 | intergenic_region, modifier, EKO05_001827-EKO05_001828 | EKO05_001827: hypothetical protein with similarity to fungal Salicylate monooxygenases, EKO05_001828: **hypothetical apoplastic effector** | EKO05_001827: non-effector, EKO05_001828: 1.578 (0.861 Apoplastic effector) | LTR/Gypsy |
| ctg04_2574966 | - | - | - | - | 0.0185 | - | intergenic_region, modifier, EKO05_003200 | hypothetical protein of unknown function | non-effector | DNA/hAT-Charlie |
| ctg05_77541 | - | - | - | - | - | 0.0347 | intergenic_region, modifier, EKO05_003204-EKO05_003205 | hypothetical proteins of unknown function | non-effectors | LTR/Gypsy |
| ctg07_1292465 | - | - | - | - | - | 0.0361 | missense_variant, moderate impact, EKO05_005029 | hypothetical alkaline phosphatase | 0.561 (non-effector), signal peptide detected | - |
| ctg07_899731 | - | - | - | - | - | 0.0361 | missense_variant, moderate impact, EKO05_004868 | hypothetical protein of unknown function | non-effector | - |
| ctg08_1872808 | - | Yes, region 31 | - | - | 0.0109 | - | intergenic_region, modifier, EKO05_005581-CHR_END | hypothetical protein with transmembrane domains and high similarity to hexose transporters in fungi | non-effector | LTR/Gypsy |
| ctg08_1872842 | - | Yes, region 31 | 0.02 | - | 0.0102 | - | same as ctg_08_1872808 |  |  |  |
| ctg08_1872963 | - | Yes, region 31 | 0.0182 | 0.0165 | 0.0000188 | 0.00167 | same as ctg_08_1872808 |  |  |  |
| ctg08_1875370 | - | Yes, region 31 | - | - | 0.0102 | - | same as ctg_08_1872808 |  |  |  |
| ctg08_1875396 | - | Yes, region 31 | - | - | 0.0437 | - | same as ctg_08_1872808 |  |  |  |
| ctg08_1875471 | - | Yes, region 31 | - | - | 0.00528 | - | same as ctg_08_1872808 |  |  |  |
| ctg08_448981 | Yes | Yes, region 28 | - | - | - | 0.0364 | intergenic_region, modifier, EKO05_005280-EKO05_005281 | EKO05_005280-EKO05: **hypothetical apoplastic effector**, EKO05_005281: hypothetical protein of unknown function | EKO05_005280: -2.911 (0.509, Apoplastic effector), EKO05_005281: non-effector | LTR/Gypsy |
| ctg08_98214 | Yes | Yes, region 27 | - | - | - | 0.0273 | intergenic_region, modifier, EKO05_005201-EKO05_005202 | EKO05_005201: hypothetical apoplastic effector, EKO05_005202: hypothetical protein of unknown function | EKO05_005201: -1.476 (0.654, Apoplastic effector), EKO05_005202: -3.378 (0.675 Cytoplasmic effector but no signal peptide detected) | LTR/Gypsy |
| ctg09_169049 | - | - | - | - | - | 0.0347 | upstream_gene_variant, modifier, EKO05_005613, intergenic_region, modifier, EKO05_005611-EKO05_005612 | EKO05_005611: hypothetical protein with similarity to fungal polysaccharide lyases, EKO05_005612: hypothetical RNA methyl-transferase, EKO05_005613: hypothetical protein with similarity to fungal transcription factors with a zinc-finger domain | EKO05_005611: 0.688 (non-effector), signal peptide detected, EKO05_005612: -2.433 (0.577, Cytoplasmic effector, no signal peptide detected), EKO05_005613: non-effector | LTR/Gypsy |
| ctg09_1747152 | - | - | - | - | - | 0.0347 | intergenic_region, modifier, EKO05_006084-EKO05_006085 | EKO05_006084: hypothetical protein of unknown function, EKO05_006085: hypothetical aspartate ammonia-lyase | non-effectors | LTR/Gypsy |
| ctg09_54897 | - | - | - | 0.0305 | - | - | intergenic_region, modifier, EKO05_005587-EKO05_005588 | EKO05_00558: hypothetical DNA helicase, EKO05_005588: hypothetical protein with similarity to 2-dehydropantoate 2-reductase | non-effectors | unknown repeat family |
| ctg10_10643 | Yes | Yes, region 34 | - | - | 0.00895 | 0.0374 | synonymous_variant, low impact, EKO05_006140 | hypothetical protein of unknown function | -2.614 (0.522, Cytoplasmic effector but no signal peptide detected) | DNA/Kolobok-H |
| ctg10_157955 | Yes | Yes, region 35 | - | - | - | 0.0364 | intergenic_region, modifier, EKO05_006148-EKO05_006149 | EKO05_006148: hypothtical protein of unknown function, EKO05_006149: hypothetical oxidooreductase | non-effectors | DNA/TcMar-Fot1 |
| ctg10_157964 | Yes | Yes, region 35 | - | - | - | 0.0438 | Same as ctg_10_157955 |  |  |  |
| ctg10_158053 | Yes | Yes, region 35 | - | - | - | 0.0364 | Same as ctg_10_157955 |  |  |  |
| ctg10_1866672 | - | - | - | - | - | 0.0347 | intergenic_region, modifier, EKO05_006715-EKO05_006717 | EKO05_006715: hypothetical protein of unknown function with transmembrane domains, EKO05_006717 | EKO05_006715: non-effector, EKO05_006717: -2.708 (0.627, Cytoplasmic effector but no sugnal peptide detected) | - |
| ctg12_1225837 | - | Yes, region 45 | - | 0.0126 | - | - | intergenic_region, modifier, EKO05_007471-EKO05_007472 | EKO05_007471: hypothetical protein of unknown function, EKO05_007472: **hypothetical cytoplasmic effector** | EKO05_007471: non-effector, EKO05_007472: -1.071 (0.535, Cytoplasmic effector, signal peptide detected) | LTR/Gypsy |
| ctg12_1234964 | Yes | Yes, region 45 | - | - | - | 0.0273 | same as ctg_12_1225837 |  |  |  |
| ctg12_1234984 | Yes | Yes, region 45 | - | - | - | 0.0273 | same as ctg_12_1225837 |  |  |  |
| ctg12_1530432 | - | - | - | - | - | 0.00737 | intergenic_region, modifier, EKO05_007577- EKO05_007578 | EKO05_007577: hypothetical HECT-type E3 ubiquitin transferase, EKO05_007578: hypothetical cytoplasmic protein of unknown function | EKO05_007577: non-effector, EKO05_007578: 0.811 (non-effector) | - |
| ctg12_763572 | - | - | - | - | 0.0489 | 0.0143 | upstream_gene_variant, modifier, EKO05_007339, downstream_gene_variant, modifier, EKO05_007336, EKO05_007337, EKO05_007338 | EKO05_007336: hypothetical protein with similarity to MYG1 and fungal protein hydrolases, EKO05_007337, EKO05_007338: hypothetical GCD complex subunit gcd7, EKO05_007339: hypothetical protein-serine/threonine phosphatase | EKO05_007336: -3.188 (0.,628, Cytoplasmic effector but no signal peptide detected), EKO05_007337: -2.372 (0.732, Cytoplasmic effector but no signal peptide detected), EKO05_007338: non-effector, EKO05_007339: non-effector | - |
| ctg12_763578 | - | - | - | - | 0.0489 | 0.0143 | same as ctg_12_763572 |  |  | - |
| ctg12_763589 | - | - | - | - | 0.0489 | 0.0143 | same as ctg_12_763572 |  |  | - |
| ctg13_1358937 | - | Yes, region 50 | 0.0275 | 0.0225 | - | - | intergenic_region, modifier, EKO05_007941-EKO05_007942 | EKO05_007941: hypothetical protein with transmembrane domain siliar to fungal oligopeptide transporters, EKO05_007942: hypothetical protein of unknown function | non-effectors | LTR/Gypsy |
| ctg13_1474524 | - | Yes, region 51 | - | - | - | 0.0347 | intergenic_region, modifier, EKO05_007958-EKO05_007959 | EKO05_007958: hypothetical glycoside hydrolase, EKO05_007959: hypothetical protein of unknown function | EKO05_007958: -0.94 (0.878, Apoplastic effector), EKO05_007959: non-effector | DNA/TcMar-Fot1 |
| ctg13_192218 | - | - | - | - | - | 0.0347 | intergenic_region, modifier, EKO05_007660-EKO05_007661 | EKO05_007660: PQ-Loop domain containing protein, EKO05_007661: hypothetical protein with similarity to fungal DNA polymerases | non-effectors | DNA/Merlin |
| ctg13_842542 | - | - | - | - | - | 0.0347 | intergenic_region, modifier, EKO05_007845-EKO05_007846 | EKO05_007845: hypothetical protein of unknown function with transmembrane domains, EKO05_007846: hypothetical protein with similarity to clavaminate synthase proteins in fungi | EKO05_007845: non-effector, EKO05_007846: non-effector | LTR/Gypsy |
| ctg13_842613 | - | - | - | - | - | 0.0347 | same as ctg_13_842542 |  |  |  |
| ctg14_1472702 | - | Yes, region 55 | - | 0.0269 | - | - | intergenic_region, modifier, EKO05_008461-EKO05_008462 | EKO05_008461 hypothetical transmembrane protein, putative Mg transporter zinc transport protein, EKO05_008462: hypothetical apoplastic effector | EKO05_008461: non effector, EKO05_008462: 0.847 (0.671 - Apoplastic effector) | LTR/Gypsy |
| ctg14_1472722 | - | Yes, region 55 | - | 0.014 | - | - | same as ctg_14_1472702 |  |  |  |
| ctg14_1472755 | - | Yes, region 55 | - | 0.0156 | - | - | same as ctg_14_1472702 |  |  |  |
| ctg15_1247671 | - | - | - | - | - | 0.0347 | intergenic_region, modifier, EKO05_008834-EKO05_008835 | EKO05_008834: hypothetical transcription factor, EKO05_008835: Hypothetical protein with similarity to glycerate kinanses in fungi | non-effectors | LTR/Copia |
| ctg15_1503691 | - | Yes, region 62 | 0.0179 | - | 0.0154 | - | downstream_gene_variant, modifier, EKO05_008905 | EKO05_008905: DNA helicase with a transmembrane domain | non-effectors | unknown family |
| ctg15_1503696 | Yes | Yes, region 62 | 0.0199 | - | - | - | same as ctg_15_1503691 |  |  |  |
| ctg15_1503844 | - | Yes, region 62 | 0.00481 | 0.0155 | 0.00797 | - | same as ctg_15_1503691 |  |  |  |
| ctg15_1504039 | - | Yes, region 62 | 0.00481 | 0.0155 | 0.00797 | - | same as ctg_15_1503691 |  |  |  |
| ctg15_1504127 | - | Yes, region 62 | 0.00481 | 0.0155 | 0.00797 | - | same as ctg_15_1503691 |  |  |  |
| ctg15_1504334 | - | Yes, region 62 | 0.0166 | 0.0131 | 0.0236 | - | same as ctg_15_1503691 |  |  |  |
| ctg17_452611 | - | - | - | - | - | 0.0347 | intergenic_region, modifier, EKO05_009528-EKO05_009529: hypothetical vitamin B6 transporter | EKO05_009528: hypothetical protein disulfide-isomerase, EKO05_009529: non-effector | EKO05_009528: 0.254 (0.511, Cytoplasmic effector, signal peptide detected), EKO05_009529 |  |
| ctg19_104632 | - | - | - | - | 0.0489 | 0.0143 | downstream_gene_variant, modifier, EKO05_010260 | hypothetical protein of unknown function | non-effector | DNA/TcMar-Fot1 |
| ctg19_1102753 | - | - | - | - | 0.0146 | 0.019 | intergenic_region, modifier, EKO05_010490-EKO05_010491 | EKO05_010490: hypothetical acetylxylan esterase, EKO05_010491: **hypothetical apoplastic effector** | EKO05_010490: 0.82 (non-effector), EKO05_010491: 2.117 (0.835: Apoplastic effector), signal peptide detected | LTR/Gypsy |
| ctg19_1102770 | - | - | - | - | 0.0421 | 0.0116 | same as ctg_19_1102753 |  |  |  |
| ctg19_1102779 | - | - | - | - | 0.0421 | 0.0116 | same as ctg_19_1102753 |  |  |  |
| ctg19_1154999 | yes | Yes, region 71 | 0.00638 | - | 0.00293 | 0.000122 | intergenic_region, modifier, EKO05_010492-EKO05_010493 | EKO05_010492: hypothetical apoplastic effector, EKO05_010493: amino acid transporter with transmembrane domains | EKO05_010492: 2.244 (0.895, Apoplastic effector), EKO05_010493: non-effector | LTR/Gypsy |
| ctg19_1155004 | yes | Yes, region 71 | 0.00638 | - | 0.00293 | 0.000122 | same as ctg_19_1154999 |  |  |  |
| ctg19_1294610 | - | - | - | - | - | 0.0191 | intergenic_region, modifier, EKO05_010512-EKO05_010513 | EKO05_010512: hypothetical protein with a transmembrane domain; EKO05_010513: hypothetical protein, putative metal ion binding | EKO05_010512: -598 (0.636, Cytoplamic effector but no signal peptide detected), EKO05_010513: non-effector | DNA/TcMar-Fot1 |
| ctg19_422984 | - | - | - | - | - | 0.0347 | downstream_gene_variant, modifier, EKO05_010323, intergenic_region, modifier, EKO05_010324-EKO05_010325 | EKO05_010323: hypothetical apoplastic effect, EKO05_010324: hypothetical protein of unknown function, EKO05_010325: dehydroquinate dehydratase | EKO05_010323: -0.15 (0.560, Apoplastic protein), EKO05_010324: non-effector, EKO05_010325: non-effector | LTR/Gypsy |
| ctg20_20369 | - | - | - | - | 0.0269 | 0.0394 | upstream_gene_variant, modifier, EKO05_010525, intergenic_region, modifier, EKO05_010525-EKO05_010526 | EKO05_010525: hypothetical protein of unknown function with a transmembrane domain, EKO05_010526: hypothetical protein of unknown function | EKO05_010525: -3.346 (0.608, cytoplasmic effector but no signal peptide detected), EKO05_010526: non-effector | - |
| ctg20_20645 | - | - | - | - | 0.0378 | 0.0124 | same as ctg_20_20369 |  |  |  |
| ctg21_487980 | - | - | - | - | - | 0.0347 | missense_variant, moderate impact, EKO05_010986 | hypothetical protein of unknown function | non-effector | - |

**Table S6.** Genetic variations predicted to be associated with aggressiveness of *Ascochyta rabiei* isolates on different chickpea cultivars detected using DAPC analyses. Loading refers to the contribution of each locus to the observed differentiation, and sub-population refers to the clonal lineage (subpopulation) of isolates within which the association was detected (clusters/subpopulations detected in Figure 1). Predector score is provided with EffectorP 3.0 score in parentheses. Non-effector indicates a negative Predector score and > 0.5 non-effector score from EffectorP 3.0.

| **Region** | **Variation** | **Chickpea variety** | **Loading** | **Sub-population** | **Variant type and location** | **Putative protein function** | **Predector Score (EffectorP 3.0 score)** | **Repeat family** | **region detected in Pyseer** |
| --- | --- | --- | --- | --- | --- | --- | --- | --- | --- |
| 1 | ctg01_847649 | Genesis090 | 0.00519917 | Cluster3 | synonymous_variant, low impact, EKO05_000231 | Hypothetical protein with similarity to fungal RNA helicases | non-effector | - | No |
|  | ctg01_847688 | Genesis090 | 0.00639330 | Cluster3 | same as ctg01_847649 |  |  |  |  |
| 2 | ctg01_1510437 | PBA HatTrick | 0.00638156 | Cluster2 | missense_variant, moderate impact, EKO05_000430 | hypothetical protein of unknown function | non-effector | - | No |
| 3 | ctg01_1654657 | PBA Seamer | 0.01106706 | Cluster 1 | intergenic_region, modifier, EKO05_000484-EKO05_000485 | EKO05_000484: hypothetical protein with high similarity to fungal GTPases, EKO05_000485 hypothetical DNA binding protein with a transmembrane protein | non-effectors | LTR/Copia | Yes |
|  | ctg01_1665154 | PBA Seamer | 0.01106706 | Cluster 1 | same as ctg01_1654657 |  |  |  |  |
| 4 | ctg01_3363573 | Genesis090 | 0.01889603 | Cluster 2 | synonymous_variant, low impact, EKO05_001005 | hypothetical protein of unknown function | non-effector | DNA/Kolobok-H | No |
|  |  | ICC3996 | 0.00533564 | Cluster 2 |  |  |  |  |  |
| 5 | ctg02_5952 | PBA HatTrick | 0.00670503 | Cluster 1 | upstream_gene_variant, modifier, EKO05_001006 | hypothetical protein of unknown function | -2.078 (0.519, cytoplasmic effector but no signal peptide detected) | DNA/Kolobok-H | No |
|  | ctg02_7704 | ICC3996 | 0.00964527 | Cluster 1 | same as ctg02_5952 |  |  |  |  |
|  |  | PBA HatTrick | 0.00800386 | Cluster 1 |  |  |  |  |  |
| 6 | ctg02_352849 | PBA Seamer | 0.01106706 | Cluster 1 | intergenic_region, modifier, EKO05_001071-EKO05_001072 | EKO05_001071: hypothetical HET domain containing protein with a transmembrane domain, EKO05_001072: Phosphotransferase | non-effectors | LTR/Gypsy | Yes |
|  | ctg02_364191 | PBA Seamer | 0.01106706 | Cluster 1 | same as ctg02_352849 |  |  |  |  |
| 7 | ctg02_1660285 | PBA Seamer | 0.00772822 | Cluster 2 | missense_variant, moderate impact, EKO05_001443 | Hypothetical protein with similarity to fungal RNA helicases | non-effector | - | No |
| 8 | ctg02_2690139 | PBA Seamer | 0.01106706 | Cluster 1 | missense_variant, moderate impact, EKO05_001768 | hypothetical protein with similarity to fungal transcription factors with a zinc-finger domain | non-effector | - | Yes |
| 9 | ctg02_2788591 | PBA HatTrick | 0.00638157 | Cluster 2 | intergenic_region, modifier, EKO05_001778-EKO05_001779 | EKO05_001778: Hypothetical RNA helicase, EKO05_001779: hypothetical protein of unknown function | EKO05_001778: non-effector, EKO05_001779: 3.941 (0.862, Apoplastic effector) | LTR | No |
| 10 | ctg02_2843207 | ICC3996 | 0.01253206 | Cluster 1 | intergenic_region, modifier, EKO05_001780-CHR_END | hypothetical protein of unknown function | -2.613 (0.567, Cytoplasmic effector but no signal peptide detected) | DNA/Kolobok-H | No |
| 11 | ctg03_401008 | PBA HatTrick | 0.00638157 | Cluster 2 | intragenic_variant, modifier, EKO05_001821- EKO05_001822 | Hypothetical proteins of unknown function | EKO05_001821: non-effector, EKO05_001822: -2.643 (0.846, Cytoplasmic effector but no signal peptide) | LTR | No |
| 12 | ctg03_1415178 | PBA Seamer | 0.01106706 | Cluster 1 | intergenic_region, modifier, EKO05_002061-EKO05_002062 | EKO05_002061: hypothetical protein of unknown function, EKO05_002062: **hypothetical apoplastic effector** | EKO05_002061: non-effector, EKO05_002062: 0.676 (0.709, Apoplastic effector) | LTR/Gypsy | Yes |
| 13 | ctg03_1878424 | PBA Seamer | 0.00650038 | Cluster 1 | missense_variant, moderate impact, EKO05_002214 | Hypothetical protein of unknown functions | non-effector | - | No |
| 14 | ctg03_2014060 | PBA HatTrick | 0.00638156 | Cluster 2 | intergenic_region, modifier, EKO05_002252-EKO05_002253 | EKO05_002252-EKO05: hypothetical protein of unknown function, EKO05_002253: hypothetical protein with similarity to oxidoreductases | non-effectors | LTR/Gypsy | No |
|  | ctg03_2025024 | Genesis090 | 0.03814387 | Cluster 2 | same as ctg03_2014060 |  |  |  |  |
|  | ctg03_2039741 | PBA Seamer | 0.00772822 | Cluster 2 | same as ctg03_2014060 |  |  |  |  |
| 15 | ctg03_2550442 | PBA HatTrick | 0.00638157 | Cluster 2 | intergenic_region, modifier, EKO05_002440-EKO05_002441 | EKO05_002440: hypothetical protein of unknown function, EKO05_002441: hypothetical protein with similarity to fungal Salicylate monooxygenases | non-effectors | LTR/Gypsy | No |
|  | ctg03_2550592 | PBA HatTrick | 0.00638157 | Cluster 2 | same as ctg03_2550442 |  |  |  |  |
|  | ctg03_2556946 | PBA HatTrick | 0.00638157 | Cluster 2 | same as ctg03_2550442 |  |  |  |  |
| 16 | ctg04_114446 | PBA HatTrick | 0.00638157 | Cluster 2 | intergenic_region, modifier, EKO05_002463-EKO05_002464 | hypothetical protein of unknown function | Non-effectors | LINE/Tad1 | No |
|  | ctg04_132375 | PBA Seamer | 0.00772822 | Cluster 2 | same as ctg04_114446 |  |  |  |  |
| 17 | ctg04_1366342 | PBA HatTrick | 0.00638157 | Cluster 2 | intergenic_region, modifier, EKO05_002887-EKO05_002888 | hypothetical proteins of unknown function | EKO05_002887: -3.042 (0.724, Cytoplasmic effector but no signal peptide), EKO05_002888: non-effector | LTR/Gypsy | No |
| 18 | ctg05_4856 | Genesis090 | 0.01056547 | Cluster 1 | downstream_gene_variant, modifier, EKO05_003201 | ATP-dependant DNA helicase | non-effector | LTR | No |
| 19 | ctg05_298757 | PBA Seamer | 0.00554413 | Cluster 2 | intergenic_region, modifier, EKO05_003224-EKO05_003225 | hypothetical proteins of unknown function | EKO05_003224: -1.432 (0.504, apoplastic effector), EKO05_003225: -2.315 (0.753 Cytoplasmic effector but no signal peptide) | DNA/TcMar-Fot1 | No |
| 20 | ctg05_2328653 | Genesis090 | 0.00504442 | Cluster 1 | intergenic_region, modifier, EKO05_003903-EKO05_003904 | EKO05_003903: **hypothetical effector** based on Predector score, EKO05_003904: hypothetical protein of unknown function | EKO05_003903: 2.473 (non-effector), EKO05_003904: non-effector (0.779 Cytoplasmic effector but no signal peptide detected). | LTR | No |
|  | ctg05_2374056 | PBA HatTrick | 0.00758349 | Cluster 1 | Same as ctg05_2328653 |  |  |  |  |
|  |  | ICC3996 | 0.00684786 | Cluster 1 |  |  |  |  |  |
| 21 | ctg06_1231 | Genesis090 | 0.04526454 | Cluster 2 | intergenic_region, modifier, CHR_START-EKO05_003906 | ATP-dependant DNA helicase | non-effector | DNA/hAT-Charlie | No |
|  |  |  | 0.00960261 | Cluster 1 |  |  |  |  |  |
|  |  | PBA HatTrick | 0.00564256 | Cluster 2 |  |  |  |  |  |
| 22 | ctg06_32631 | PBA HatTrick | 0.01611868 | Cluster 1 | intergenic_region, modifier, EKO05_003912-EKO05_003913 | EKO05_003912: hypothetical cytoplasmic protein of unknown function with a signal peptide, EKO05_003913: **hypothetical apoplastic effector** | EKO05_003912: 2.473 (non-effector), EKO05_003913: 3.884 (0.89, Apoplastic effector) | DNA/TcMar-Fot1 | No |
| 23 | ctg06_364193 | PBA Seamer | 0.00554413 | Cluster 2 | intergenic_region, modifier, EKO05_003977-EKO05_003978 | hypothetical proteins of unknown function | EKO05_003977: -3.338 (0.839, Cytoplasmic effector but no signal peptide), EKO05_003977: non-effector | LTR | No |
|  | ctg06_367615 | PBA Seamer | 0.00772822 | Cluster 2 | Same as ctg06_364193 |  |  |  |  |
| 24 | ctg07_33708 | PBA Seamer | 0.00772822 | Cluster 2 | intergenic_region, modifier, EKO05_004591-EKO05_004592 | hypothetical proteins of unknown function | EKO05_004591: -3.042 (0.587, cytoplasmic effector but no signal peptide), EKO05_004592: -2.076 (0.842, cytoplasmic effector but no signal peptide) | - | No |
| 25 | ctg07_619873 | PBA HatTrick | 0.00638156 | Cluster 2 | intergenic_region, modifier, EKO05_004769-EKO05_004770 | EKO05_004769: Triacylglycerol lipase, EKO05_004770: hypothetical secreted protein with similarity to fungal transcription factors | non-effectors | LTR/Gypsy | No |
| 26 | ctg08_4047 | ICC3996 | 0.00502948 | Cluster 2 | downstream_gene_variant, modifier, EKO05_005191 | Hypothetical protein with high similarity to fungal helicases | Non-effector | unknown | No |
| 27 | ctg08_98214 | PBA Seamer | 0.01106706 | Cluster 1 | intergenic_region, modifier, EKO05_005201-EKO05_005202 | EKO05_005201: hypothetical apoplastic effector, EKO05_005202: hypothetical protein of unknown function | EKO05_005201: -1.476 (0.654, Apoplastic effector), EKO05_005202: -3.378 (0.675 Cytoplasmic effector but no signal peptide detected) | LTR/Gypsy | Yes |
| 28 | ctg08_448981 | PBA Seamer | 0.01461423 | Cluster 1 | intergenic_region, modifier, EKO05_005280-EKO05_005281 | EKO05_005280-EKO05: **hypothetical apoplastic effector**, EKO05_005281: hypothetical protein of unknown function | EKO05_005280: -2.911 (0.509, Apoplastic effector), EKO05_005281: non-effector | LTR/Gypsy | Yes |
|  | ctg08_498148 | PBA HatTrick | 0.00638156 | Cluster 2 | same as ctg08_448981 |  |  |  |  |
| 29 | ctg08_1002177 | PBA Seamer | 0.00903923 | Cluster 1 | intergenic_region, modifier, EKO05_005327-EKO05_005328 | EKO05_005327: hypothetical tetrahydrofolate synthase; EKO05_005328: hypothetical protein similar to rapid alkalinization factors (RALFs) | EKO05_005327: non-effector, EKO05_005328: 1.001 (0.739, **Apoplastic effector**) | DNA/TcMar-Fot1 | No |
|  |  | PBA HatTrick | 0.00613061 | Cluster 1 |  |  |  |  |  |
|  | ctg08_1002191 | PBA Seamer | 0.00903923 | Cluster 1 | same as ctg08_1002177 |  |  |  |  |
|  |  | PBA HatTrick | 0.00613061 | Cluster 1 |  |  |  |  |  |
| 30 | ctg08_1071033 | PBA HatTrick | 0.00638157 | Cluster 2 | intergenic_region, modifier, EKO05_005332-EKO05_005333 | EKO05_005332: hypothetical protein of unknown function, EKO05_005333: hypothetical cytoplasmic protein of unknown function with a signal peptide | EKO05_005332: -2.923 (0.666, Cytoplasmic effector but no signal peptide detected), EKO05_005333: non-effector | DNA/Merlin | No |
| 31 | ctg08_1874732 | ICC3996 | 0.0059840 | Cluster 2 | intergenic_region, modifier, EKO05_005581-CHR_END | hypothetical protein with transmembrane domains and high similarity to sugar transporters in fungi | non-effector | LTR/Gypsy | No |
| 32 | ctg09_207905 | Genesis090 | 0.00519918 | Cluster3 | synonymous_variant, low impact, EKO05_005621 | hypothetical protein of unknown function with similarity to fungal glucosyl hydrolases | -1.468 (0.663, cytoplasmic effector but no signal peptide) | - | No |
| 33 | ctg09_1023045 | Genesis090 | 0.00638157 | Cluster 2 | intergenic_region, modifier, EKO05_005894-EKO05_005895 | EKO05_005894: hypothetical cytoplasmic protein with similarity to fungal carbohydrate esterases, EKO05_005895: hypothetical protein of unknown function | EKO05_005894: 0.266 (0.704, non-effector), EKO05_005895: non-effector | DNA/TcMar-Fot1 | No |
| 34 | ctg10_10643 | ICC3996 | 0.00988308 | Cluster 2 | synonymous_variant, low impact, EKO05_006140 | hypothetical protein of unknown function | -2.614 (0.522 Cytoplasmic effector but no signal peptide detected) | DNA/Kolobok-H | Yes |
|  |  | ICC3996 | 0.00535019 | Cluster 3 |  |  |  |  |  |
|  |  | Genesis090 | 0.00754685 | Cluster 1 |  |  |  |  |  |
| 35 | ctg10_157955 | PBA Seamer | 0.01461423 | Cluster 1 | intergenic_region, modifier, EKO05_006148-EKO05_006149 | EKO05_006148: hypothetical protein of unknown function, EKO05_006149: hypothetical oxidoreductase | non-effectors | DNA/TcMar-Fot1 | Yes |
|  | ctg10_157964 | PBA Seamer | 0.01461423 | Cluster 1 | same as ctg10_157955 |  |  |  |  |
|  | ctg10_157976 | PBA Seamer | 0.01461424 | Cluster 1 | same as ctg10_157955 |  |  |  |  |
|  | ctg10_158053 | PBA Seamer | 0.01461423 | Cluster 1 | same as ctg10_157955 |  |  |  |  |
| 36 | ctg10_1065329 | PBA Seamer | 0.00772822 | Cluster 2 | synonymous_variant, low impact, EKO05_006457 | hypothetical protein of unknown function with similarity to ribosome bioproteinsis protein tsr1 | Non-effector | - | No |
| 37 | ctg10_1922864 | ICC3996 | 0.00857352 | Cluster 2 | downstream_gene_variant, modifier, EKO05_006718, EKO05_006719, EKO05_006720 | EKO05_006718: hypothetical protein of unknown function, EKO05_006719- EKO05_006720: hypothetical proteins of unknown function with similarity to DNA helicases | EKO05_006718: -3.179 (0.744 cytoplasmic effector but no signal peptide), EKO05_006719: non-effector, EKO05_006720: -3.042 (0.796, cytoplasmic effector but no signal peptide) | DNA/Kolobok-H | No |
| 38 | ctg11_4855 | Genesis090 | 0.01338384 | Cluster 2 | intergenic_region, modifier, CHR_START-EKO05_006721 | ATP-dependant DNA helicase | non-effector | DNA/hAT-Charlie | No |
| 39 | ctg 11_73427 | PBA Seamer | 0.00772822 | Cluster 2 | intergenic_region, modifier, EKO05_006726-EKO05_006727 | Hypothetical proteins of unknown function | non-effectors | LTR_retrotransposon | No |
| 40 | ctg11_355082 | PBA Seamer | 0.00554413 | Cluster 2 | intergenic_region, modifier, EKO05_006755-EKO05_006756 | Hypothetical proteins of unknown function | non-effectors | LTR/Gypsy | No |
| 41 | ctg11_500888 | PBA Seamer | 0.00650038 | Cluster 2 | intergenic_region, modifier EKO05_006793-EKO05_006794 | Hypothetical proteins of unknown function | non-effectors | - | No |
| 42 | ctg 11_1696540 | PBA Seamer | 0.00554413 | Cluster 2 | intergenic_region, modifier, EKO05_007124-EKO05_007125 | EKO05_007124: hypothetical protein with a signal peptide and similarity to peptidases, EKO05_007125: hypothetical protein of unknown function | EKO05_007124: 0.873 (non-effector), EKO05_007125: non-effector | LTR/Gypsy | No |
| 43 | ctg12_2872 | ICC3996 | 0.00804649 | Cluster 1 | upstream_gene_variant, modifier, EKO05_007132, EKO05_007133, EKO05_007134 | EKO05_007132: hypothetical protein on unknown function with a transmembrane domain, EKO05_007133: hypothetical DNA helicase, EKO05_007134: hypothetical protein of unknown function | non-effectors | DNA/hAT-Charlie | No |
|  | ctg12_3434 | Genesis090 | 0.005501452 | Cluster 2 | same as ctg12_2872 |  |  |  |  |
|  | ctg12_9283 | Genesis090 | 0.00757841 | Cluster 1 | downstream_gene_variant, modifier, EKO05_007132, EKO05_007133; intergenic_region, modifier, EKO05_007134-EKO05_007135 | as listed for ctg12_3434, EKO05_007135: hypothetical protein of unknown function | non-effectors | - | No |
|  | ctg12_10098 | ICC3996 | 0.00821206 | Cluster 2 | same as ctg12_9283 |  |  | LINE/Tad1 |  |
|  |  | PBA HatTrick | 0.00771867 | Cluster 1 |  |  |  |  |  |
|  | ctg12_64739 | PBA Seamer | 0.00554413 | Cluster 2 | same as ctg12_9283 |  |  | LINE/Tad1 |  |
|  | ctg12_64766 | PBA Seamer | 0.00554413 | Cluster 2 | same as ctg12_9283 |  |  | LINE/Tad1 |  |
|  | ctg12_64796 | PBA Seamer | 0.00554413 | Cluster 2 | same as ctg12_9283 |  |  | LINE/Tad1 |  |
| 44 | ctg12_902286 | PBA Seamer | 0.00554413 | Cluster 2 | intergenic_region, modifier, EKO05_007365-EKO05_007366 | Hypothetical proteins of unknown function | non-effector | LTR/Gypsy | No |
| 45 | ctg12_1234964 | PBA Seamer | 0.01106706 | Cluster 1 | intergenic_region, modifier, EKO05_007471-EKO05_007472 | EKO05_007471: hypothetical protein of unknown function, EKO05_007472: **hypothetical cytoplasmic effector** | EKO05_007471: non-effector, EKO05_007472: -1.071 (0.535, Cytoplasmic effector, signal peptide detected) | LTR/Gypsy | Yes |
|  | ctg12_1234970 | PBA Seamer | 0.01106706 | Cluster 1 | same as ctg12_1234964 |  |  |  |  |
|  | ctg12_1234984 | PBA Seamer | 0.01106706 | Cluster 1 | same as ctg12_1234964 |  |  |  |  |
| 46 | ctg12_1732383 | PBA HatTrick | 0.00638157 | Cluster 2 | intergenic_region, modifier, EKO05_007599-EKO05_007600 | EKO05_007599: **hypothetical apoplastic effector**, EKO05_007600: hypothetical protein of unknown function | EKO05_007599: 1.891 (0.808), Apoplastic effector | LTR/Copia | No |
| 47 | ctg13_5628 | Genesis090 | 0.01566084 | Cluster 3 | missense_variant, moderate impact, EKO05_007602, upstream_gene_variant, modifier, EKO05_007603 | Hypothetical protein with a transmembrane domain | non-effector | DNA/Kolobok-H | No |
|  | ctg13_5645 | Genesis090 | 0.00758281 | Cluster 2 | same as ctg13_5628 |  |  |  |  |
|  |  |  | 0.01533566 | Cluster 3 |  |  |  |  |  |
|  | ctg13_6273 | ICC3996 | 0.00671848 | Cluster 2 | missense_variant, low impact, EKO05_007602 | see above | non-effector | DNA/Kolobok-H | No |
| 48 | ctg13_39540 | PBA Seamer | 0.00554413 | Cluster 2 | intergenic_region, modifier, EKO05_007605-EKO05_007606 | Hypothetical proteins of unknown function | non-effectors | - | No |
| 49 | ctg13_822157 | PBA HatTrick | 0.00638157 | Cluster 2 | intergenic_region, modifier, EKO05_007845-EKO05_007846 | Hypothetical proteins of unknown function | non-effectors | LTR/Gypsy | No |
| 50 | ctg13_1358890 | ICC3996 | 0.00559696 | Cluster 2 | intergenic_region, modifier, EKO05_007941-EKO05_007942 | EKO05_007941: hypothetical protein with transmembrane domain siliar to fungal oligopeptide transporters, EKO05_007942: hypothetical protein of unknown function | non-effectors | LTR/Gypsy | Yes |
| 51 | ctg13_1474245 | PBA Seamer | 0.00772821 | Cluster 2 | intergenic_region, modifier, EKO05_007958-EKO05_007959 | EKO05_007958: hypothetical glycoside hydrolase, EKO05_007959: hypothetical protein of unknown function | EKO05_007958: -0.94 (0.878, **Apoplastic effector**), EKO05_007959: non-effector | DNA/TcMar-Fot1 | Yes |
|  | ctg13_1512039 | PBA HatTrick | 0.00638157 | Cluster 2 | same as ctg13_1474245 |  |  |  |  |
| 52 | ctg14_5555 | Genesis090 | 0.00516043 | Cluster 2 | downstream_gene_variant, modifier, EKO05_007960, upstream_gene_variant, modifier, EKO05_007961 | EKO05_007960: hypothetical DNA helicase with a transmembrane domain, EKO05_007961: hypothetical protein of unknown function | non-effectors | DNA/Kolobok-H | No |
|  | ctg14_6054 | Genesis090 | 0.01980639 | Cluster 3 | missense_variant, moderate impact, EKO05_007960 | as listed for ctg14_5555 |  | DNA/Kolobok-H |  |
|  |  |  | 0.02917822 | Cluster 2 |  |  |  |  |  |
|  |  | ICC3996 | 0.02164216 | Cluster 2 |  |  |  |  |  |
|  | ctg14_7795 | PBA Seamer | 0.00675340 | Cluster 3 | missense_variant, moderate impact, EKO05_007960 | as listed for ctg14_5555 |  |  |  |
|  | ctg14_7982 | Genesis090 | 0.00624210 | Cluster 3 | missense_variant, low impact, EKO05_007960 | as listed for ctg14_5555 |  |  |  |
|  | ctg14_8013 | ICC3996 | 0.03508566 | Cluster 2 | missense_variant, moderate impact, EKO05_007960 | as listed for ctg14_5555 |  | DNA/Kolobok-H |  |
| 53 | ctg14_128865 | PBA Seamer | 0.00772822 | Cluster2 | upstream_gene_variant, modifier, EKO05_007983, EKO05_007984, EKO05_007985 |  |  | - |  |
|  | ctg14_128972 | PBA Seamer | 0.00772822 | Cluster2 | same as ctg14_128865 |  |  | - |  |
|  | ctg14_128985 | PBA Seamer | 0.00772822 | Cluster2 | same as ctg14_128865 |  |  | - |  |
| 54 | ctg14_1128676 | Genesis090 | 0.0114578 | Cluster 1 | upstream_gene_variant, modifier, EKO05_008334, EKO05_008335 | EKO05_008334: telomere length regulation protein, EKO05_008335: ARP2/3 actin-organizing complex subunit Sop2 | non-effectors | - | No |
|  | ctg14_1128681 | Genesis090 | 0.01229865 | Cluster 1 | same as ctg14_1128676 |  |  |  |  |
| 55 | ctg14_1471881 | PBA Seamer | 0.009731598 | Cluster 2 | intergenic_region, modifier, EKO05_008461-EKO05_008462 | EKO05_008461 hypothetical transmembrane protein, putative Mg transporter zinc transport protein, EKO05_008462: **hypothetical apoplastic effector** | EKO05_008461: non effector, EKO05_008462: 0.847 (0.671 - Apoplastic effector) | LTR/Gypsy | Yes |
|  |  | Genesis090 | 0.00969599 | Cluster 1 |  |  |  |  |  |
|  | ctg14_1472550 | ICC3996 | 0.022482043 | Cluster 2 | same as ctg14_1471881 |  |  |  |  |
|  | ctg14_1472885 | ICC3996 | 0.007242732 | Cluster 3 | same as ctg14_1471881 |  |  |  |  |
|  | ctg14_1473114 | PBA HatTrick | 0.00512752 | Cluster 1 | same as ctg14_1471881 |  |  |  |  |
|  |  | ICC3996 | 0.00732687 | Cluster 3 |  |  |  |  |  |
|  |  | ICC3996 | 0.00670026 | Cluster 2 |  |  |  |  |  |
|  | ctg14_1473121 | ICC3996 | 0.01110045 | Cluster 2 | same as ctg14_1471881 |  |  |  |  |
|  |  |  | 0.00791407 | Cluster 3 |  |  |  |  |  |
|  |  | PBA Seamer | 0.00707533 | Cluster 2 |  |  |  |  |  |
|  |  | PBA HatTrick | 0.00509446 | Cluster 2 |  |  |  |  |  |
|  |  |  | 0.00512752 | Cluster 1 |  |  |  |  |  |
|  | ctg14_1473129 | Genesis090 | 0.00795573 | Cluster 2 | same as ctg14_1473121 |  |  |  |  |
|  |  | PBA Seamer | 0.005560388 | Cluster 2 |  |  |  |  |  |
|  |  | PBA HAtTrick | 0.00512752 | Cluster 1 |  |  |  |  |  |
|  |  | ICC3996 | 0.00547121 | Cluster 3 |  |  |  |  |  |
|  | ctg14_1473135 | Genesis090 | 0.01022132 | Cluster 2 | same as ctg14_1473121 |  |  |  |  |
|  | ctg14_1474269 | Genesis090 | 0.01648159 | Cluster 2 | same as ctg14_1473121 |  |  |  |  |
|  | ctg14_1474335 | ICC3996 | 0.01738445 | Cluster 2 | same as ctg14_1473121 |  |  |  |  |
|  | ctg14_1474599 | PBA Seamer | 0.00561586 | Cluster 2 | same as ctg14_1473121 |  |  |  |  |
|  | ctg14_1474618 | ICC3996 | 0.00883718 | Cluster 2 | same as ctg14_1473121 |  |  |  |  |
|  |  | PBA Seamer | 0.00650151 | Cluster 2 |  |  |  |  |  |
|  | ctg14_1475147 | Genesis090 | 0.005236324 | Cluster 2 | same as ctg14_1473121 |  |  |  |  |
|  | ctg14_1486225 | PBA HatTrick | 0.00638157 | Cluster 2 | same as ctg14_1473121 |  |  |  |  |
| 56 | ctg14_261308 | PBA Seamer | 0.00772822 | Cluster 2 | synonymous_variant, low impact, EKO05_008034 | Rho GTPase activating protein | non-effector | - | No |
| 57 | ctg14_368054 | PBA Seamer | 0.00554413 | Cluster 2 | intergenic_region, modifier, EKO05_008063-EKO05_008064 | Hypothetical proteins of unknown function | non-effectors | LTR/Gypsy | No |
| 58 | ctg15_4819 | Genesis090 | 0.00614213 | Cluster 1 | downstream_gene_variant, modifier, EKO05_008466 | Hypothetical protein with similarity to DNA helicases | non-effector | LTR_Retrotransposon | No |
| 59 | ctg15_958303 | Genesis090 | 0.00519918 | Cluster 3 | synonymous_variant, low impact, EKO05_008749 | Glyceraldehyde 3-phosphate dehydrogenase | 0.254 | - | No |
| 60 | ctg15_1240897 | PBA HatTrick | 0.00638157 | Cluster 2 | intergenic_region, modifier, EKO05_008834-EKO05_008835 | EKO05_008834: hypothetical protein on unknown function, EKO05_008835: Glycerate 2-kinase | non-effectors | LTR/Gypsy | No |
| 61 | ctg15_1498555 | PBA HatTrick | 0.00638157 | Cluster 2 | upstream_gene_variant, modifier, EKO05_008903 | hypothetical protein of unknown function | non-effector | DNA/Kolobok-H | No |
| 62 | ctg15_1503638 | Genesis090 | 0.02617707 | Cluster 1 | downstream_gene_variant, modifier, EKO05_008905 | EKO05_008905: DNA helicase with a transmembrane domain | non-effectors | repeat region, unknown family |  |
|  |  | PBA HatTrick | 0.006215497 | Cluster 1 |  |  |  |  |  |
|  |  | ICC3996 | 0.005432997 | Cluster 1 |  |  |  |  |  |
|  |  | ICC3996 | 0.00520425 | Cluster 2 |  |  |  |  |  |
|  | ctg15_1503666 | Genesis090 | 0.01738969 | Cluster 1 | same as ctg15_1503638 |  |  |  |  |
|  |  | PBA HatTrick | 0.008843174 | Cluster 1 |  |  |  |  |  |
|  |  | ICC3996 | 0.006234541 | Cluster 1 |  |  |  |  |  |
|  | ctg15_1503691 | Genesis090 | 0.00613647 | Cluster 3 | same as ctg15_1503638 |  |  |  |  |
|  |  | ICC3996 | 0.00873476 | Cluster 2 |  |  |  |  |  |
|  | ctg15_1503696 | Genesis090 | 0.00784413 | Cluster 3 | same as ctg15_1503638 |  |  |  |  |
|  |  | ICC3996 | 0.02923538 | Cluster 2 |  |  |  |  |  |
| 63 | ctg16_5960 | Genesis090 | 0.01381917 | Cluster 2 | upstream_gene_variant, modifier, EKO05_008906, EKO05_008907 | EKO05_008906: ATP-dependant helicase with a transmemrane domain, EKO05_008907: hypothetical non-cytoplasmic protein | non-effectors | DNA/Kolobok-H | No |
|  |  |  | 0.00882174 | Cluster 1 |  |  |  |  |  |
|  |  |  | 0.00882174 | Cluster 2 |  |  |  |  |  |
|  |  | ICC3996 | 0.007729018 | Cluster 2 |  |  |  |  |  |
| 64 | ctg16_949437 | PBA Seamer | 0.00772822 | Cluster 2 | missense_variant, moderate impact, EKO05_009246 | hypothetical protein of unknown function | non-effector | - | No |
| 65 | ctg16_1448099 | PBA HatTrick | 0.00638157 | Cluster 2 | downstream_gene_variant, modifier, EKO05_009408 | hypothetical protein of unknown function | non-effector | DNA/hAT-Charlie | No |
| 66 | ctg18_77913 | PBA HatTrick | 0.00638157 | Cluster 2 | intergenic_region, modifier, EKO05_009874-EKO05_009875 | hypothetical proteins of unknown function | non-effectors | - | No |
| 67 | ctg18_543932 | PBA Seamer | 0.00554413 | Cluster 2 | intergenic_region, modifier, EKO05_010017-EKO05_010018 | hypothetical proteins of unknown function | non-effectors | DNA/TcMar-Fot1 |  |
| 68 | ctg19_438760 | PBA Seamer | 0.00554413 | Cluster 2 | intergenic_region, modifier, EKO05_010324-EKO05_010325 | EKO05_010324: hypothetical protein of unknown function, EKO05_010325: 3-dehydroquinate dehydratase (3-dehydroquinase) | non-effectors | LTR/Gypsy | No |
|  | ctg19_448773 | PBA Seamer | 0.00772822 | Cluster 2 | same as ctg19_438760 |  |  |  |  |
| 69 | ctg19_52221 | PBA Seamer | 0.00554413 | Cluster 2 | upstream_gene_variant, modifier, EKO05_010258 | Hypothetical **Apoplastic effector** | 1.724 (0.711 Apoplastic effector) |  |  |
|  | ctg19_52228 | PBA Seamer | 0.00554413 | Cluster 2 | same as ctg19_52221 |  |  |  |  |
| 70 | ctg19_714127 | PBA Seamer | 0.00554413 | Cluster 2 | intergenic_region, modifier, EKO05_010378-EKO05_010379 | EKO05_010378: hypothetical protein of unknown function, EKO05_010379: hypothetical lycerophosphodiester phosphodiesterase | non-effector | - | No |
| 71 | ctg19_1154947 | PBA Seamer | 0.01089556 | Cluster 3 | intergenic_region, modifier, EKO05_010492-EKO05_010493 | EKO05_010492: **hypothetical apoplastic effector**, EKO05_010493: amino acid transporter with transmembrane domains | EKO05_010492: 2.244 (0.895, Apoplastic effector), EKO05_010493: non-effector | LTR/Gypsy | Yes |
|  |  | PBA HatTrick | 0.005482253 | Cluster 3 |  |  |  |  |  |
|  | ctg19_1154955 | PBA Seamer | 0.01089556 | Cluster 3 | same as ctg19_1154947 |  |  |  |  |
|  |  | PBA HatTrick | 0.00548225 | Cluster 3 |  |  |  |  |  |
|  | ctg19_1154999 | PBA Seamer | 0.01089556 | Cluster 3 | same as ctg19_1154947 |  |  |  |  |
|  |  | PBA HatTrick | 0.00504485 | Cluster 3 |  |  |  |  |  |
|  | ctg19_1155004 | PBA Seamer | 0.01089556 | Cluster 3 | same as ctg19_1154947 |  |  |  |  |
|  |  | PBA HatTrick | 0.005482253 | Cluster 3 |  |  |  |  |  |
|  | ctg19_1155061 | PBA Seamer | 0.01089556 | Cluster 3 | same as ctg19_1154947 |  |  |  |  |
|  |  | PBA HatTrick | 0.005044855 | Cluster 3 |  |  |  |  |  |
|  | ctg19_1155087 | PBA Seamer | 0.01089556 | Cluster 3 | same as ctg19_1154947 |  |  |  |  |
|  | ctg19_1155193 | PBA Seamer | 0.01077375 | Cluster 3 | same as ctg19_1154947 |  |  |  |  |
|  | ctg19_1155290 | PBA Seamer | 0.008570053 | Cluster 3 | same as ctg19_1154947 |  |  |  |  |
|  | ctg19_1155317 | PBA Seamer | 0.01089556 | Cluster 3 | same as ctg19_1154947 |  |  |  |  |
|  | ctg19_1155345 | PBA Seamer | 0.008575561 | Cluster 2 | same as ctg19_1154947 |  |  |  |  |
| 72 | ctg20_10055 | ICC3996 | 0.01182705 | Cluster 3 | upstream_gene_variant, modifier, EKO05_010523, EKO05_010524 | EKO05_010523: DNA helicase with a transmembrane domain, EKO05_010524: hypothetical non-cytoplasmic protein of unknown function | non-effectors | LTR_retrotransposon | No |
|  |  | PBA HatTrick | 0.005817049 | Cluster 1 |  |  |  |  |  |
|  | ctg20_6546 | ICC3996 | 0.00554558 | Cluster 1 | missense_variant, moderate impact, EKO05_010523 | as listed for ctg20_10055 |  |  |  |
|  | ctg20_6490 | ICC3996 | 0.01815324 | Cluster 2 | missense_variant, moderate impact, EKO05_010523 | as listed for ctg20_10055 |  |  |  |
|  | ctg20_7536 | ICC3996 | 0.0262593 | Cluster 2 | same as ctg20_10055 |  |  |  |  |
|  |  | Genesis090 | 0.00555415 | Cluster 2 |  |  |  |  |  |
|  | ctg20_7609 | Genesis090 | 0.00589039 | Cluster 2 | same as ctg20_10055 |  |  |  |  |
|  |  | ICC3996 | 0.00621149 | Cluster 2 |  |  |  |  |  |
|  | ctg20_7659 | ICC3996 | 0.00621149 | Cluster 2 | same as ctg20_10055 |  |  |  |  |
|  | ctg20_8917 | ICC3996 | 0.02051209 | Cluster 2 | same as ctg20_10055 |  |  |  |  |
|  | ctg20_9902 | PBA HatTrick | 0.00742752 | Cluster 1 | same as ctg20_10055 |  |  |  |  |
| 73 | ctg20_254353 | Genesis090 | 0.01066103 | Cluster 1 | missense_variant, moderate impact, EKO05_010561 | hypothetical HECT-type E3 ubiquitin transferase | non-effector | - | No |
| 74 | ctg20_332105 | ICC3996 | 0.00949823 | Cluster 2 | intergenic_region, modifier, EKO05_010582-EKO05_010583 | EKO05_010582: **hypothetical apoplastic effector,** EKO05_010583: hypothetical Xylan 1,4-beta-xylosidase with a transmembrane domain | EKO05_010582: 0.981 (0.607, Apoplastic effector), EKO05_010583: non-effector | LTR_retrotransposon | No |
|  | ctg20_332127 | ICC3996 | 0.00949823 | Cluster 2 | same as ctg20_332105 |  |  |  |  |
|  | ctg20_342634 | PBS Seamer | 0.00554413 | Cluster 2 | same as ctg20_332105 |  |  |  |  |
| 75 | ctg21_153969 | PBA HatTrick | 0.00638156 | Cluster 2 | intergenic_region, modifier, EKO05_010892-EKO05_010893 | EKO05_010892: acetyltransferase, EKO05_010893: hypothetical protein of unknown function with a transmembrane domain | non-effectors | LTR/Gypsy | No |
|  | ctg21_174491 | PBA HatTrick | 0.00638156 | Cluster 2 | same as ctg21_153969 |  |  |  |  |
|  | ctg21_181970 | PBA HatTrick | 0.00638157 | Cluster 2 | same as ctg21_153969 |  |  |  |  |
|  | ctg21_185186 | PBA HatTrick | 0.00628248 | Cluster 2 | same as ctg21_153969 |  |  |  |  |
|  | ctg22_4536 | Genesis090 | 0.00842176 | Cluster 3 | downstream_gene_variant, modifier, EKO05_011220, EKO05_011221 | EKO05_011220: DNA helicase with a transmembrane domain, EKO05_011221: hypothetical DNA helicase | non-effectors | DNA/Kolobok-H | No |
|  | ctg22_4612 | Genesis090 | 0.00698665 | Cluster 1 | same as ctg22_4536 |  |  |  |  |
|  |  | Genesis090 | 0.0053572 | Cluster 2 |  |  |  |  |  |
|  | ctg22_4619 | Genesis090 | 0.00698665 | Cluster 1 | same as ctg22_4536 |  |  |  |  |
|  | ctg22_4623 | ICC3996 | 0.00536662 | Cluster 1 | same as ctg22_4536 |  |  |  |  |
|  |  | Genesis090 | 0.00656978 | Cluster 1 |  |  |  |  |  |
|  |  |  | 0.00617011 | Cluster 2 |  |  |  |  |  |

**Table S7.** Genetic variations between closely related *Ascochyta rabiei* isolates with contrasting aggressiveness (Pathogenicity Group (PG) 0/1 versus 4/5)

| **Pairwise comparison** | **Cluster** | **Variation** | **Variant detected in DAPC** | **DNA region detected in DAPC** | **Variant detected in other pairwise comparisons?** | **Variant type and location** | **Putative protein function** | **Predector Score (EffectorP 3.0 score)** | **Repeat family** | **region detected in Pyseer** |
| --- | --- | --- | --- | --- | --- | --- | --- | --- | --- | --- |
| AR0184 (PG0) and AR0304 (PG4) | 2 | ctg01_3369118 | - | Yes | - | intergenic_region, modifier, EKO05_001005-CHR_END | hypothetical protein of unknown function | non-effector | DNA/Kolobok-H | Yes |
|  |  | ctg06_1231 | Yes | - | Ar0230/AR0226 | intergenic_region, modifier, CHR_START-EKO05_003906 | ATP-dependant DNA helicase | non-effector | DNA/hAT-Charlie | No |
|  |  | ctg06_32631 | Yes | Yes | AR0037/AR0033, 16RUP013/ andF17191-1 | intergenic_region, modifier, EKO05_003912-EKO05_003913 | EKO05_003912: hypothetical cytoplasmic protein of unknown function with a signal peptide, EKO05_003913: **hypothetical apoplastic effector** | EKO05_003912: 2.473 (non-effector), EKO05_003913: 3.884 (0.89, Apoplastic effector) | DNA/TcMar-Fot1 | No |
|  |  | ctg13_5645 | Yes | Yes | Ar0230/AR0226, AR0219/AR0212, AR0189/AR0179 | missense_variant, moderate impact, EKO05_007602 | Hypothetical protein with a transmembrane domain | non-effector | DNA/Kolobok-H | No |
|  |  | ctg14_6054 | Yes | - | Ar0230/AR0226, AR0189/AR0179 | missense_variant, moderate impact, EKO05_007960 | hypothetical DNA helicase with a transmembrane domain | non-effector | DNA/Kolobok-H | No |
|  |  | ctg14_7982 | Yes | Yes | - | synonymous_variant, low impact, EKO05_007960 | hypothetical DNA helicase with a transmembrane domain | non-effector | DNA/Kolobok-H | No |
|  |  | ctg20_7609 | Yes | Yes | AR0189/AR0179 | upstream_gene_variant, modifier, EKO05_010523, EKO05_010524 | EKO05_010523: DNA helicase with a transmembrane domain, EKO05_010524: hypothetical non-cytoplamic protein of unknown function | non-effectors | LTR_retrotransposon | No |
| AR0037 (PG0) and AR0033) | 2 | ctg05_2388561 | - | Yes | - | missense_variant, moderate impact, EKO05_003905 | hypothetical DNA helicase with a transmembrane domain | non-effector | DNA/Kolobok-H | No |
|  |  | ctg06_32631 | Yes | Yes | AR0184/AR0304, 16RUP013/F17191-1 | See variations between AR0184 (PG0) and AR0304 (PG4) |  |  |  | No |
|  |  | ctg08_1874732 | Yes | Yes | - | intergenic_region, modifier, EKO05_005581-CHR_END | hypothetical protein with transmembrane domains and high similarity to sugar transporters in fungi | non-effector | LTR/Gypsy | No |
|  |  | ctg10_10643 | Yes | Yes | - | synonymous_variant, low impact, EKO05_006140 | hypothetical protein of unknown function | -2.614 (0.522 Cytoplasmic effector but no signal peptide detected) | DNA/TcMar-Fot1 | Yes |
|  |  | ctg13_5578 | - | Yes | - | synonymous_variant\|LOW\|EKO05_007602 | Hypothetical protein with a transmembrane domain | non-effector | DNA/Kolobok-H | No |
|  |  | ctg13_6273 | Yes | Yes | - | synonymous_variant\|LOW\|EKO05_007602- | Hypothetical protein with a transmembrane domain | non-effector | DNA/Kolobok-H | No |
| 16RUP013 (PG1) andF17191-1 (PG5) | 1 | ctg05_2328653 | Yes | Yes | - | upstream_gene_variant, modifier, EKO05_003903 | EKO05_003903: hypothetical effector based on Predector score | EKO05_003903: 2.473 (non-effector) | LTR | No |
|  |  | ctg05_2328659 | - | Yes | - | same as ctg05_2328653 |  |  |  | No |
|  |  | ctg05_2328866 | - | Yes | - | same as ctg05_2328653 |  |  |  | No |
|  |  | ctg06_32631 | Yes | Yes | AR0184/AR0304, AR0037/AR0033 | See variations between AR0184 (PG0) and AR0304 (PG4) |  |  |  | No |
|  |  | ctg09_10576 | - | - | - | stop_gained, high impact, EKO05_005585 | hypothetical protein of unknown function | non-effector | DNA/Kolobok-H | No |
|  |  | ctg14_1128676 | Yes | Yes | - | upstream_gene_variant, modifier, EKO05_008334, EKO05_008335 | EKO05_008334: telomere length regulation protein, EKO05_008335: ARP2/3 actin-organizing complex subunit Sop2 | non-effectors | - | No |
|  |  | ctg14_1474269 | Yes | Yes | - | intergenic_region, modifier, EKO05_008461-EKO05_008462 | EKO05_008461 hypothetical transmembrane protein, putative Mg transporter zinc transport protein, EKO05_008462: hypothetical apoplastic effector | EKO05_008461: non effector, EKO05_008462: 0.847 (0.671 - Apoplastic effector) | LTR/Gypsy | Yes |
|  |  | ctg14_1474599 | Yes | Yes | AR0189/AR0179 | same as ctg14_1474269 |  |  |  | Yes |
|  |  | ctg15_1503638 | Yes | Yes | - | downstream_gene_variant, modifier, EKO05_008905 | EKO05_008905: DNA helicase with a transmembrane domain | non-effectors | repeat region, unknown family | Yes |
|  |  | ctg15_1503666 | Yes | Yes | - | same as ctg15_1503638 |  |  |  | Yes |
|  |  | ctg20_945131 | - | - | - | intergenic_region, modifier, EKO05_010755-EKO05_010756 | EKO05_010755: hypothetical protein with transmembrane domains and similarity to copper transporters in fungi, EKO05_010756: hypothetical cytoplasmic protein with similarity to alpha-trehalases | EKO05_010755: non-effector, EKO05_010756: 0.37 (non-effector) | LTR/Copia | No |
| Ar0230 (PG0) and AR0226 (PG4) | 3 | ctg06_1231 | Yes | Yes | AR0184/AR0304 | See variations between AR0184 (PG0) and AR0304 (PG4) |  |  |  |  |
|  |  | ctg13_5391 | - | Yes | - | missense_variant, moderate impact, EKO05_007602 | Hypothetical protein with a transmembrane domain | non-effector | DNA/Kolobok-H | No |
|  |  | ctg13_5645 | Yes | Yes | AR0184/AR0304, AR0219/AR0212, AR0189/AR0179 | missense_variant, moderate impact, EKO05_007602 | Hypothetical protein with a transmembrane domain | non-effector | DNA/Kolobok-H | No |
|  |  | ctg14_6054 | Yes | Yes | AR0184/AR0304, AR0189/AR0179 | missense_variant, moderate impact, EKO05_007960 | EKO05_007960: hypothetical DNA helicase with a transmembrane domain | non-effectors | DNA/Kolobok-H | No |
|  |  | ctg15_1503691 | yes | Yes | AR0219/AR0212 | downstream_gene_variant, modifier, EKO05_008905 | EKO05_008905: DNA helicase with a transmembrane domain | non-effectors | unknown family | Yes |
|  |  | ctg15_1503696 | Yes | Yes | AR0219/AR0212 | same as ctg15_1503691 |  |  |  | Yes |
|  |  | ctg17_6257 | - | - | - | synonymous_variant, low impact, EKO05_009409\|EKO05_009409 | hypothetical DNA helicases | non-effector | DNA/Kolobok-H | No |
| AR0219 (PG0) and AR0212 (PG4) | 3 | ctg03_2025024 | Yes | Yes | - | intergenic_region, modifier, EKO05_002252-EKO05_002253 | EKO05_002252-EKO05: hypothetical protein of unknown function, EKO05_002253: hypothetical protein with similarity to oxidoreductases | non-effectors | LTR/Gypsy | Yes |
|  |  | ctg08_1875607 | - | Yes | - | intergenic_region, modifier, EKO05_005581-CHR_END | hypothetical protein with transmembrane domains and high similarity to sugar transporters in fungi | non-effector | LTR/Gypsy | No |
|  |  | ctg13_5645 | Yes | Yes | AR0184/AR0304, Ar0230/AR0226, AR0189/AR0179 | missense_variant, moderate impact, EKO05_007602 | Hypothetical protein with a transmembrane domain | non-effector | DNA/Kolobok-H | No |
|  |  | ctg13_870015 | - | Yes | - | intergenic_region, modifier, EKO05_007845-EKO05_007846 | EKO05_007845: hypothetical protein of unknown function with transmembrane domains, EKO05_007846: hypothetical protein with similarity to clavaminate synthase proteins in fungi | EKO05_007845: non-effector, EKO05_007846: non-effector | LTR/Gypsy | Yes |
|  |  | ctg14_5555 | Yes | Yes | - | downstream_gene_variant, modifier, EKO05_007960, upstream_gene_variant, modifier, EKO05_007961 | EKO05_007960: hypothetical DNA helicase with a transmembrane domain, EKO05_007961: hypothetical protein of unknown function | non-effectors | DNA/Kolobok-H | No |
|  |  | ctg14_1527717 | - | - | - | missense_variant, moderate impact, EKO05_008465 | hypothetical DNA helicase | noneffector | DNA/Kolobok-H | No |
|  |  | ctg14_1527726 | - | - | - | upstream_gene_variant, modifier, EKO05_008465 | hypothetical DNA helicase | noneffector | DNA/Kolobok-H | No |
|  |  | ctg14_1527758 | - | - | - | same as ctg14_1527726 |  |  |  | No |
|  |  | ctg15_1503691 | Yes | Yes | Ar0230/AR0226 | downstream_gene_variant, modifier, EKO05_008905 | EKO05_008905: DNA helicase with a transmembrane domain | non-effectors | unknown family | Yes |
|  |  | ctg15_1503696 | Yes | Yes | Ar0230/AR0226 | same as ctg15_1503691 |  |  |  | Yes |
|  |  | ctg15_1504673 | - | Yes | AR0189, AR0179 | same as ctg15_1503691 |  |  |  | Yes |
|  |  | ctg20_326251 | - | Yes | - | intergenic_region, modifier, EKO05_010582-EKO05_010583 | EKO05_010582: hypothetical apoplastic effector, EKO05_010583: hypothetical Xylan 1,4-beta-xylosidase with a transmembrane domain | EKO05_010582: 0.981 (0.607, Apoplastic effector), EKO05_010583: non-effector | LTR_retrotransposon | No |
|  |  | ctg22-12495 | - | Yes | - | upstream_gene_variant, modifier, EKO05_011220-EKO05_011221 | EKO05_011220: DNA helicase with a transmembrane domain, EKO05_011221: hypothetical DNA helicase | non-effectors | DNA/Kolobok-H | No |
| AR0189 (PG0) and AR0179 (PG5) | 3 | ctg01_3363573 | Yes | Yes | - | synonymous_variant, low impact, EKO05_001005 | hypothetical protein of unknown function | non-effector | DNA/Kolobok-H | No |
|  |  | ctg13_3621 | - | Yes | - | upstream_gene_varian, modifier, EKO05_007603 | Hypothetical protein with a transmembrane domain | non-effector | DNA/Kolobok-H | No |
|  |  | ctg13_5645 | Yes | Yes | AR0184/AR0304, Ar0230/AR0226, AR0219/ AR0212 | missense_variant, moderate impact, EKO05_007602, upstream_gene_varian, modifier, EKO05_007603 | Hypothetical protein with a transmembrane domain | non-effector | DNA/Kolobok-H | No |
|  |  | ctg14_6054 | Yes | Yes | AR0184/AR0304, Ar0230/AR0226 | missense_variant, moderate impact, EKO05_007960 | EKO05_007960: hypothetical DNA helicase with a transmembrane domain, EKO05_007961: hypothetical protein of unknown function | non-effector | DNA/Kolobok-H | No |
|  |  | ctg14_1474335 | Yes | Yes | - | intergenic_region, modifier, EKO05_008461-EKO05_008462 | EKO05_008461 hypothetical transmembrane protein, putative Mg transporter zinc transport protein, EKO05_008462: hypothetical apoplastic effector | EKO05_008461: non effector, EKO05_008462: 0.847 (0.671 - Apoplastic effector) | LTR/Gypsy | No |
|  |  | ctg14_1474599 | Yes | Yes | 16RUP013/F17191-1 | same as ctg14_1474335 |  |  |  |  |
|  |  | ctg14_1474618 | Yes | Yes | - | same as ctg14_1474335 |  |  |  |  |
|  |  | ctg14_1475238 | Yes | Yes | - | same as ctg14_1474335 |  |  |  |  |
|  |  | ctg15_1504673 | - | Yes | AR0219/AR0212 | downstream_gene_variant, modifier, EKO05_008905 | EKO05_008905: DNA helicase with a transmembrane domain | non-effectors | unknown family | No |
|  |  | ctg20_7609 | Yes | Yes | AR0184/AR0304 | upstream_gene_variant, modifier, EKO05_010523, EKO05_010524 | EKO05_010523: DNA helicase with a transmembrane domain, EKO05_010524: hypothetical non-cytoplamic protein of unknown function | non-effectors | LTR_retrotransposon | No |
|  |  | ctg22_4536 | Yes | Yes | - | downstream_gene_variant, modifier, EKO05_011220, EKO05_011221 | EKO05_011220: DNA helicase with a transmembrane domain, EKO05_011221: hypothetical DNA helicase | non-effectors | DNA/Kolobok-H | No |
|  |  | ctg22_10183 | - | Yes | - | upstream_gene_variant, modifier, EKO05_011220, EKO05_011221 | EKO05_011220: DNA helicase with a transmembrane domain, EKO05_011221: hypothetical DNA helicase | non-effectors | DNA/Kolobok-H | No |

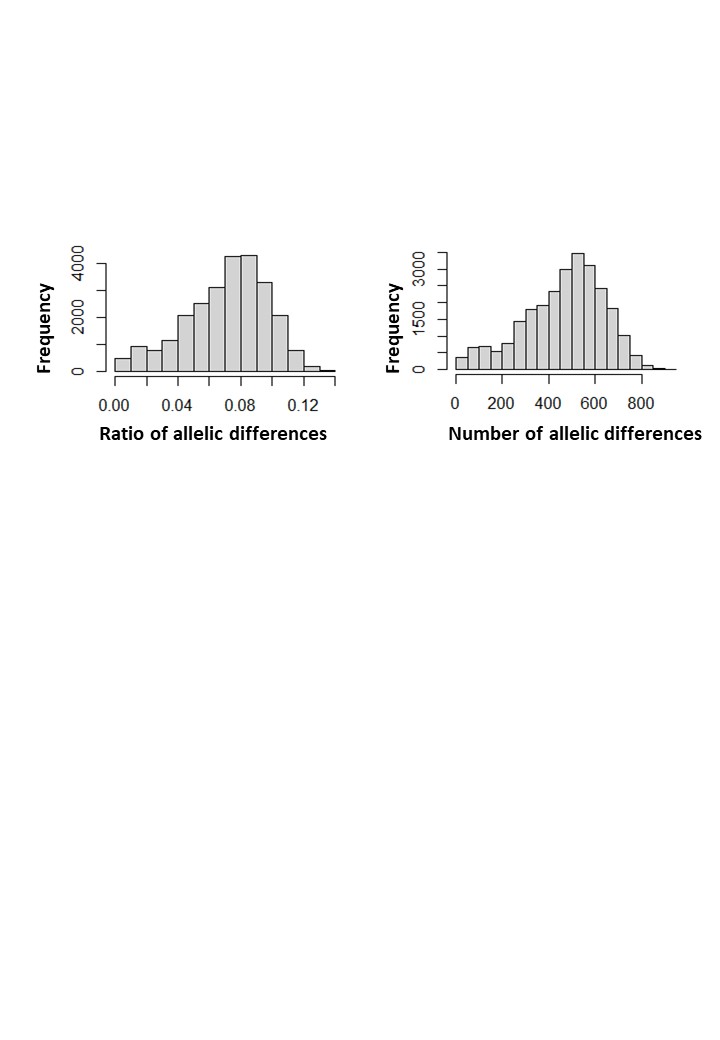

**Fig. S1.** Histograms of genetic distances between *Ascochyta rabiei* isolates collected from 2013 to 2020 in Australia, estimated via *diss.dist* function in *poppr*. The histogram on the right shows the number of SNP differences for all pairwise comparisons of *A. rabiei* isolates, and the histogram on the left shows the ratio of the number of observed differences by the number of possible differences.

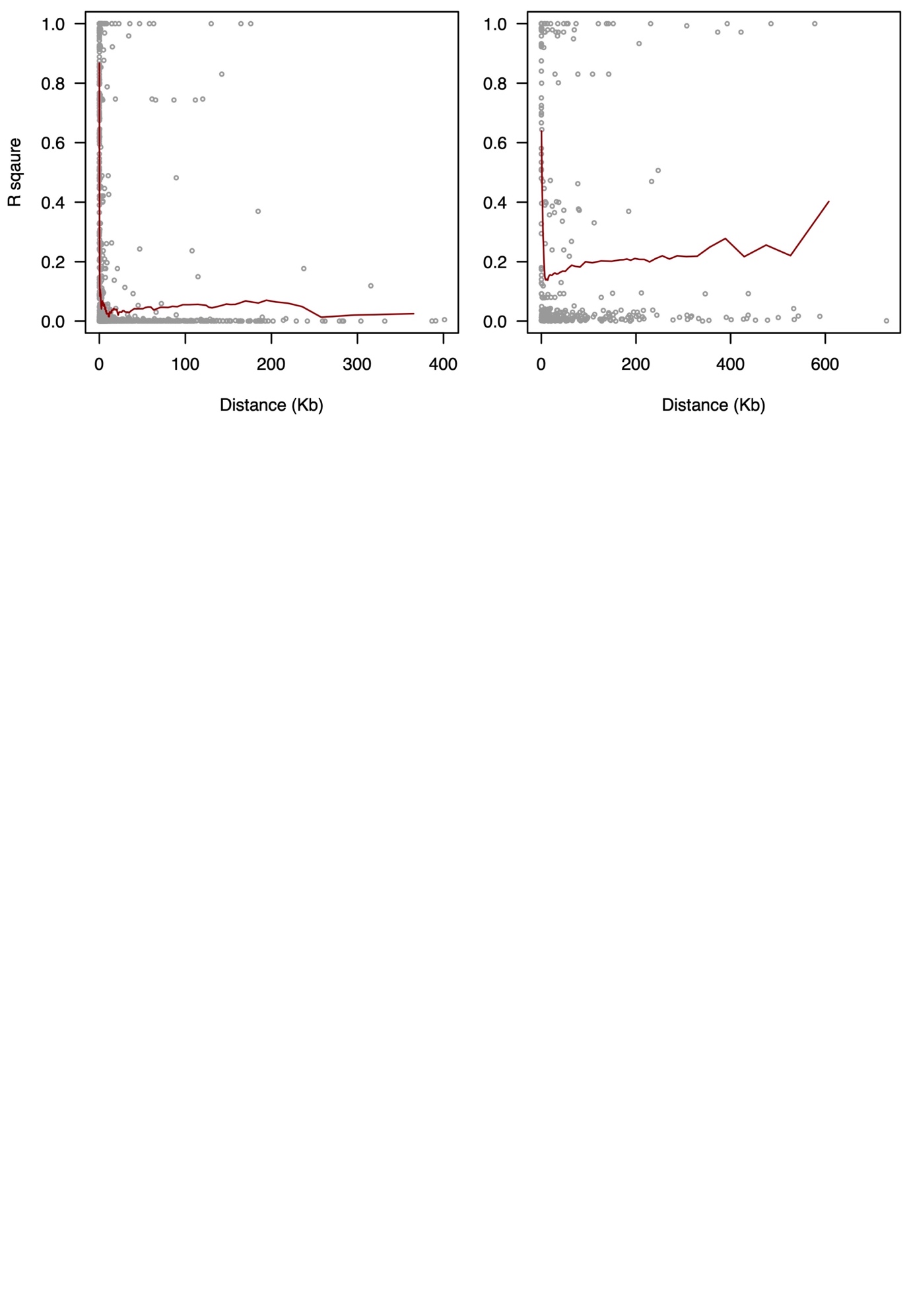

**Fig. S2.** Linkage disequilibrium (LD) decay of loci in *Ascochyta rabiei* genome (strain ArME14) measured as R square of pairwise markers plotted against distance. Linkage disequilibrium was calculated on sliding windows with 100 adjacent genetic markers using GAPIT v.3. for SNP dataset filtered for variants with minimum allele frequency of 1% (left) and 5% (right). The red line is the moving average of the 10 adjacent markers.

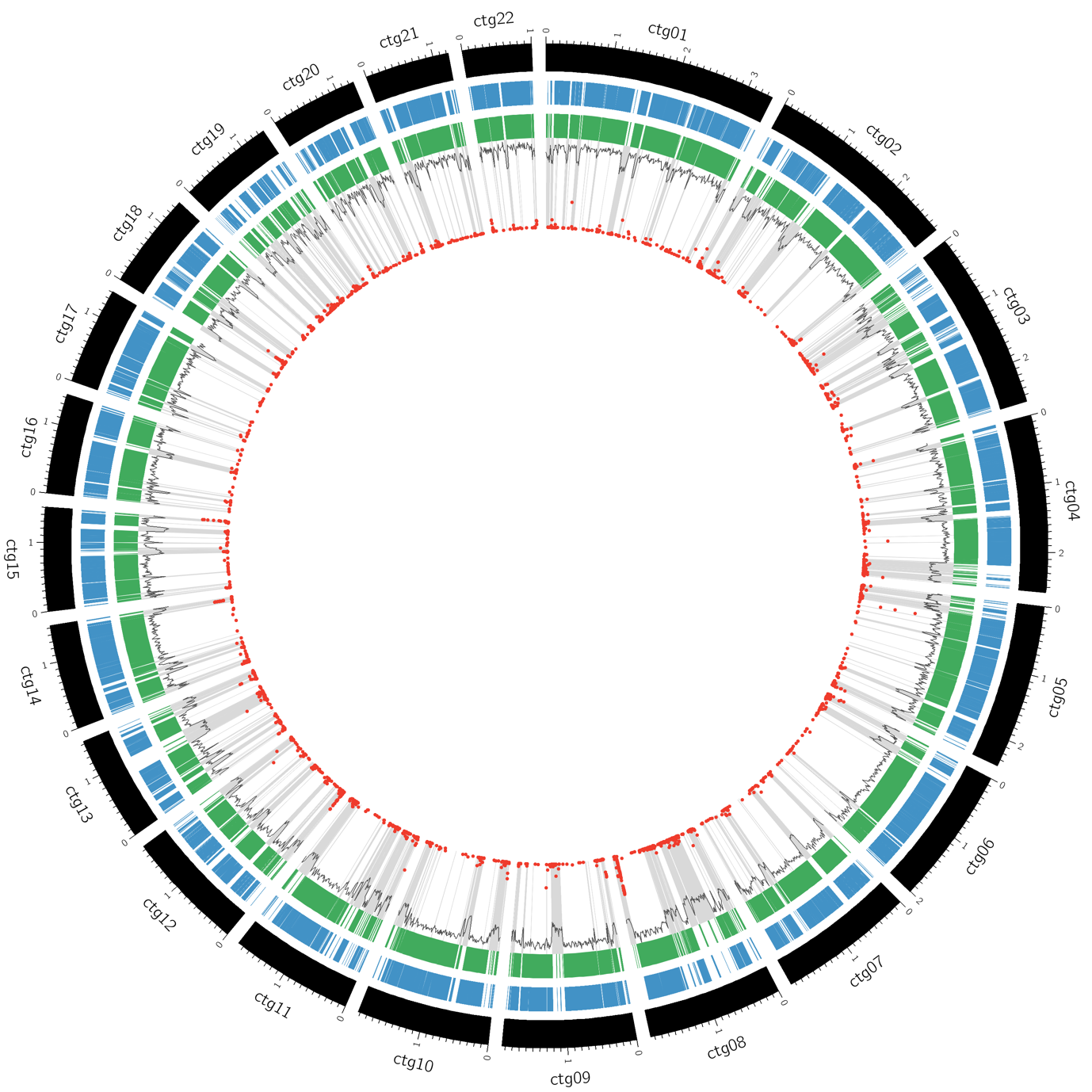

**Fig. S3.** CIRCOS plot showing key features of *A. rabiei* ArME14 genome and distribution of SNPs, showing *A. rabiei* contigs (black), annotated genes (blue), high GC regions detected in OcculterCut (green), percent GC content (gray histogram). The red points show the frequency of alternate SNP alleles at each site. Highlighted in grey are regions of low GC, which correspond to gene-sparse region and SNP hotspots.

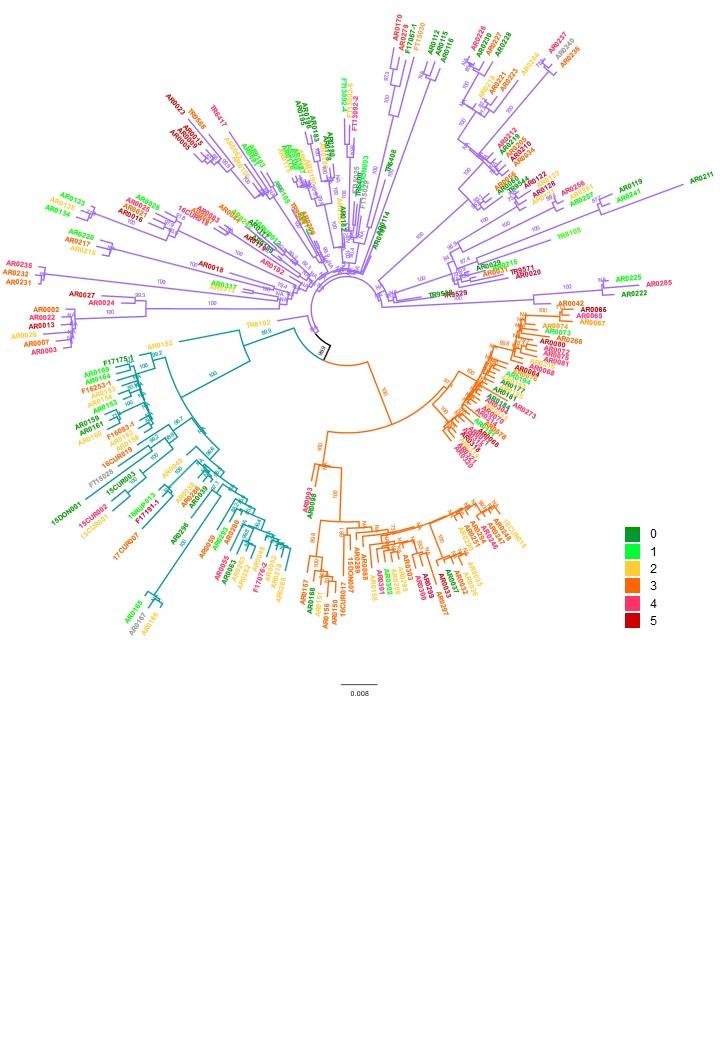

**Fig. S4.** Neighbour-joining dendrogram of 130 *Ascochyta rabiei* isolates sequenced in this study based on bitwise genetic distance using 3,283 genome-wide variations. Tip colours correspond to the aggressiveness (pathogenicity group) of the isolates as indicated in the legend. Branches coloured green, orange and purple denote isolates identified in clusters 1, 2, and 3 in the DAPC analysis shown in Figure 1.
