## Supplementary material for "Population-level whole genome sequencing of *Ascochyta rabiei* identifies genomic loci associated with isolate aggressiveness": Data: Vaghefi et al - Microbial_genomics.R

```
#######################################
### VISUALIZE OUTPUTS FROM VCFTOOLS ###
#######################################

## Site-based statistics and individual-based statistics were calculated for the original VCF using vcfTools
# This will help estimating cut-off values for site depth and quality
library(tidyverse)

## Variant quality: (Phred encoded) site quality
var_qual <- read_delim("Out.lqual", delim = "\t", col_names = c("chr", "pos", "qual"), skip = 1)
a <- ggplot(var_qual, aes(qual)) + geom_density(fill = "dodgerblue1", colour = "black", alpha = 0.3)
a + theme_light()
summary(var_qual)

## Variant mean depth: the mean depth for each variant. 
# This is the number of reads that have mapped to this position. 
var_depth <- read_delim("Out.ldepth.mean", delim = "\t", col_names = c("chr", "pos", "mean_depth", "var_depth"), skip = 1)
a <- ggplot(var_depth, aes(mean_depth)) + geom_density(fill = "dodgerblue1", colour = "black", alpha = 0.3)
a + theme_light() + xlim(0,268)
summary(var_depth)

## Variant missingness: proportion of missingness at each variant. 
# This is a measure of how many individuals lack a genotype at a call site
var_miss <- read_delim("Out.lmiss", delim = "\t", col_names = c("chr", "pos", "nchr", "nfiltered", "nmiss", "fmiss"), skip = 1)
a <- ggplot(var_miss, aes(fmiss)) + geom_density(fill = "dodgerblue1", colour = "black", alpha = 0.3)
a + theme_light()
summary(var_miss$fmiss)

## Distribution of allele frequencies will help inform  minor-allele frequency (MAF) thresholds
var_freq <- read_delim("Out.frq", delim = "\t", col_names = c("chr", "pos", "nalleles", "nchr", "a1", "a2"), skip = 1)
# find minor allele frequency
# It is an important measure because low MAF alleles may only occur in one or two individuals. 
# It is possible that some of these low frequency alleles are in fact unreliable base calls - i.e. a source of error.
var_freq$maf <- var_freq %>% select(a1, a2) %>% apply(1, function(z) min(z))
a <- ggplot(var_freq, aes(maf)) + geom_density(fill = "dodgerblue1", colour = "black", alpha = 0.3)
a + theme_light()
summary(var_freq$maf)
summary(var_freq)

## Distribution of mean depth among individuals.
ind_depth <- read_delim("Out.idepth", delim = "\t", col_names = c("ind", "nsites", "depth"), skip = 1)
a <- ggplot(ind_depth, aes(depth)) + geom_histogram(fill = "dodgerblue1", colour = "black", alpha = 0.3, binwidth = 1)
a + theme_light()
view(ind_depth)
summary(ind_depth)

## Proportion of missing data per individual
ind_miss  <- read_delim("G:/My Drive/GRDC-Ascochyta/Paper_Microbial_Genomics_2023/DATA_for_paper/Out.imiss", delim = "\t", col_names = c("ind", "ndata", "nfiltered", "nmiss", "fmiss"), skip = 1)
a <- ggplot(ind_miss, aes(fmiss)) + geom_histogram(fill = "dodgerblue1", colour = "black", alpha = 0.3, bins = 233)
a + theme_light()
view (ind_miss)
summary(ind_miss)
# The vcf file was then filtered in vcftools for 
# GT (--minDP 7 --maxDP 178 --minGQ 20), 
# sites (--max-missing 0.9 --minQ 30 --min-alleles 2 --max-alleles 2)
# individuals with > 0.1 missing data

#######################################################
### VISUALIZE OUTPUTS FROM VCFTOOLS After filtering ###
#######################################################

library(tidyverse)

## Variant quality: (Phred encoded) site quality
var_qual <- read_delim("Out2.lqual", delim = "\t", col_names = c("chr", "pos", "qual"), skip = 1)
a <- ggplot(var_qual, aes(qual)) + geom_density(fill = "dodgerblue1", colour = "black", alpha = 0.3)
a + theme_light()
summary(var_qual)

## Variant mean depth: the mean depth for each variant. 
var_depth <- read_delim("Out2.ldepth.mean", delim = "\t", col_names = c("chr", "pos", "mean_depth", "var_depth"), skip = 1)
a <- ggplot(var_depth, aes(mean_depth)) + geom_density(fill = "dodgerblue1", colour = "black", alpha = 0.3)
a + theme_light() + xlim(0,268)
summary(var_depth)

## Variant missingness: proportion of missingness at each variant. 
var_miss <- read_delim("Out2.lmiss", delim = "\t", col_names = c("chr", "pos", "nchr", "nfiltered", "nmiss", "fmiss"), skip = 1)
a <- ggplot(var_miss, aes(fmiss)) + geom_density(fill = "dodgerblue1", colour = "black", alpha = 0.3)
a + theme_light()
summary(var_miss$fmiss)

## Distribution of allele frequencies will help inform  minor-allele frequency (MAF) thresholds
var_freq <- read_delim("Out2.frq", delim = "\t", col_names = c("chr", "pos", "nalleles", "nchr", "freq"), skip = 1)
var_freq$maf <- var_freq %>% select(freq) %>% apply(1, function(z) min(z))
a <- ggplot(var_freq, aes(maf)) + geom_density(fill = "dodgerblue1", colour = "black", alpha = 0.3)
a + theme_light()
summary(var_freq$maf)
summary(var_freq)

## Distribution of mean depth among individuals.
ind_depth <- read_delim("Out2.idepth", delim = "\t", col_names = c("ind", "nsites", "depth"), skip = 1)
a <- ggplot(ind_depth, aes(depth)) + geom_histogram(fill = "dodgerblue1", colour = "black", alpha = 0.3, binwidth = 1)
a + theme_light()
view(ind_depth)
summary(ind_depth)

## Proportion of missing data per individual
ind_miss  <- read_delim("Out2.imiss", delim = "\t", col_names = c("ind", "ndata", "nfiltered", "nmiss", "fmiss"), skip = 1)
a <- ggplot(ind_miss, aes(fmiss)) + geom_histogram(fill = "dodgerblue1", colour = "black", alpha = 0.3, bins = 233)
a + theme_light()
view (ind_miss)
summary(ind_miss)

###########################################
### IMPORT DATA WITH REPLICATES REMOVED ###
###########################################

## This is all the data after filtering and removing replicates
library(vcfR)
library(poppr)
Ar230_maf1_vcf <- read.vcfR("All_230_sorted.vcf")
Ar230_genind_maf1 <- vcfR2genind(Ar230_maf1_vcf)
Ar230_genind_maf1
indNames(Ar230_genind_maf1)
X <- genind2df(Ar230_genind_maf1)
Ar230snps_maf1 <- df2genind(X, ploidy = 1, ncode=1)
indNames(Ar230snps_maf1)
Ar230_maf1 <- poppr::as.genclone(Ar230snps_maf1)

##  SET STRATA
strata.df <- read.csv("230_strata.csv", head = FALSE, sep = ",")
strata(Ar230_maf1) <- strata.df
splitStrata(Ar230_maf1) <- ~Year/State/Location/Farm/Host/Zone/ICC3996_rating/Genesis090_rating/HatTrick_rating/Seamer_rating/Path1/Path2/Oldcluster/group

## VIEW ALL STRATA
library(dplyr)
Allstrata <- strata(Ar230_maf1) %>% group_by(Year,State, Location, Farm, Host, Zone, ICC3996_rating, Genesis090_rating, HatTrick_rating, Seamer_rating, Path1, Path2) %>% summarize(Count = n())
View(Allstrata)
nameStrata(Ar230_maf1)

## DISTANCE BETWEEN INDIVIDUALS (PAIRWISE)
library("poppr")
library("pegas")
library("ape")
library("adegenet")
library("ade4")

ratio_of_allelic_differences <- diss.dist(Ar230_genind_maf1, percent = TRUE, mat = FALSE)
hist(ratio_of_allelic_differences, freq=FALSE)

Number_of_allelic_differences <- diss.dist(Ar230_genind_maf1, percent = FALSE, mat = FALSE)
hist(Number_of_allelic_differences, freq=FALSE)

ratio_of_allelic_differences <- diss.dist(Ar230_genind_maf1, percent = TRUE, mat = FALSE)
hist(ratio_of_allelic_differences)

Number_of_allelic_differences <- diss.dist(Ar230_genind_maf1, percent = FALSE, mat = FALSE)
hist(Number_of_allelic_differences)

########################################################
### IMPORT LOW Modifier DATA WITH REPLICATES REMOVED ###
########################################################

Ar230_LM_maf1_vcf <- read.vcfR("SNP_LM_230_sorted.vcf")
Ar230_LM_genind_maf1 <- vcfR2genind(Ar230_LM_maf1_vcf)
Ar230_LM_genind_maf1
indNames(Ar230_LM_genind_maf1)
X <- genind2df(Ar230_LM_genind_maf1)
Ar230snps_LM_maf1 <- df2genind(X, ploidy = 1, ncode=1)
indNames(Ar230snps_LM_maf1)
Ar230_LM_maf1 <- poppr::as.genclone(Ar230snps_LM_maf1)
Ar230_LM_maf1
indNames(Ar230_LM_maf1)

## SET STRATA
strata.df <- read.csv("230_strata.csv", head = FALSE, sep = ",")
strata(Ar230_LM_maf1) <- strata.df
splitStrata(Ar230_LM_maf1) <-  ~Year/State/Location/Farm/Host/Zone/ICC3996_rating/Genesis090_rating/HatTrick_rating/Seamer_rating/Path1/Path2/Cluster/Group

## VIEW ALL STRATA
library(dplyr)
Allstrata <- strata(Ar230_LM_maf1) %>% group_by(Year,State, Location, Farm, Host, Zone, ICC3996_rating, Genesis090_rating, HatTrick_rating, Seamer_rating, Path1, Path2) %>% summarize(Count = n())
View(Allstrata)
nameStrata(Ar230_LM_maf1)

## DELINEATE CLONAL LINEAGES
# to predict a cut off value
Ar230_LM_maf1_thresholds <- mlg.filter(Ar230_LM_maf1, distance = bitwise.dist, stats = "THRESHOLDS", threshold = 1)
pthresh  <- filter_stats(Ar230_LM_maf1, distance=bitwise.dist, plot = TRUE, stats = "THRESHOLD", threads = 1L)
cutoff_predictor(pthresh$farthest, fraction = 0.5)
mlg.filter(Ar230_LM_maf1, distance = bitwise.dist) <- 0.004502741

#############################################
###  MLG ID & Cross pop _ AFTER FILTERING ###
#############################################

Ar230_LM_maf1.MLGid <- mlg.id(Ar230_LM_maf1)
Ar230_LM_maf1.MLGid

Ar230_LM_maf1.MLGtable <- mlg.table(Ar230_LM_maf1)
write.table(Ar230_LM_maf1.MLGtable, "Ar230_LM_maf1.MLGtable_nomiss.csv")

setPop(Ar230_LM_maf1) <- ~Year
(maf5_117_Year_crosspop <- mlg.crosspop(Ar230_LM_maf1, df=TRUE))
library(ggplot2)
qplot(y=MLG, x=Population, data=maf5_117_Year_crosspop,size=Count) + theme(axis.text.x = element_text(size=10, angle=-45, hjust=0, color="black"), axis.text.y = element_text(size=10, color="black"))

setPop(Ar230_LM_maf1) <- ~State
(maf5_117_State_crosspop <- mlg.crosspop(Ar230_LM_maf1, df=TRUE))
qplot(y=MLG, x=Population, data=maf5_117_State_crosspop,size=Count) + theme(axis.text.x = element_text(size=10, angle=-45, hjust=0, color="black"), axis.text.y = element_text(size=10, color="black"))

setPop(Ar230_LM_maf1) <- ~Field
(maf5_117_Field_crosspop <- mlg.crosspop(Ar230_LM_maf1, df=TRUE))
qplot(y=MLG, x=Population, data=maf5_117_Field_crosspop,size=Count) + theme(axis.text.x = element_text(size=10, angle=-45, hjust=0, color="black"), axis.text.y = element_text(size=10, color="black"))

setPop(Ar230_LM_maf1) <- ~host
(maf5_117_host_crosspop <- mlg.crosspop(Ar230_LM_maf1, df=TRUE))
qplot(y=MLG, x=Population, data=maf5_117_host_crosspop,size=Count) + theme(axis.text.x = element_text(size=10, angle=-45, hjust=0, color="black"), axis.text.y = element_text(size=10, color="black"))

setPop(Ar230_LM_maf1) <- ~Variety
(maf5_117_Variety_crosspop <- mlg.crosspop(Ar230_LM_maf1, df=TRUE))
qplot(y=MLG, x=Population, data=maf5_117_Variety_crosspop,size=Count) + theme(axis.text.x = element_text(size=10, angle=-45, hjust=0, color="black"), axis.text.y = element_text(size=10, color="black"))

setPop(Ar230_LM_maf1) <- ~Location
(maf5_117_Location_crosspop <- mlg.crosspop(Ar230_LM_maf1, df=TRUE))
qplot(y=MLG, x=Population, data=maf5_117_Location_crosspop,size=Count) + theme(axis.text.x = element_text(size=10, angle=-45, hjust=0, color="black"), axis.text.y = element_text(size=10, color="black"))

##################################
### POPULATION DIFFERENTIATION ###
##################################

## AMOVA 
setPop(Ar230_LM_maf1) <- ~Year
popNames(Ar230_LM_maf1)
# Make a subpob without 2013 and 2014 as these are too small
Ar230_LM_maf1_year <- popsub (Ar230_LM_maf1, c(2,3,4,5))
indNames(Ar230_LM_maf1_year)
(Ar230_LM_maf1_amova <- poppr.amova(Ar230_LM_maf1_year, ~Year/State/Location, filter = TRUE, threshold = 0.00))
(Ar230_LM_maf1_amova_Pval   <- randtest(Ar230_LM_maf1_amova, nrepet = 999))
plot(Ar230_LM_maf1_amova_Pval)
# Zones
(Ar230_LM_maf1_amova <- poppr.amova(Ar230_LM_maf1_year, ~Year/Zone/Location, filter = TRUE, threshold = 0.00))
(Ar230_LM_maf1_amova_Pval   <- randtest(Ar230_LM_maf1_amova, nrepet = 999))
plot(Ar230_LM_maf1_amova_Pval)
# Without removing 2013 and 2014
(Ar230_LM_maf1_amova <- poppr.amova(Ar230_LM_maf1, ~Year/State/Location, filter = TRUE, threshold = 0.00))
(Ar230_LM_maf1_amova_Pval   <- randtest(Ar230_LM_maf1_amova, nrepet = 999))
plot(Ar230_LM_maf1_amova_Pval)
# Zones
(Ar230_LM_maf1_amova <- poppr.amova(Ar230_LM_maf1, ~Year/Zone/Location, filter = TRUE, threshold = 0.00))
(Ar230_LM_maf1_amova_Pval   <- randtest(Ar230_LM_maf1_amova, nrepet = 999))
plot(Ar230_LM_maf1_amova_Pval)

## POPULATION DIFFERENTIATION
setPop(Ar230_LM_maf1) <- ~Year
popNames(Ar230_LM_maf1)
library(mmod)
#global GST
Gst_Hedrick(Ar230_LM_maf1_year)
#pairwise
(D_Ar2021 <- pairwise_D(Ar230_LM_maf1_year))
(GST_Ar2021 <- pairwise_Gst_Nei(Ar230_LM_maf1_year))
(GST_Ar2021 <- pairwise_Gst_Hedrick(Ar230_LM_maf1_year))

##  DISTANCE TREE GENLIGHT OBJECT for LM loci
Ar230_LM_maf1_vcf <- read.vcfR("SNP_LM_230_sorted.vcf")
Ar230_LM_maf1_vcf
Ar230_LM_maf1_genlight <- vcfR2genlight(Ar230_LM_maf1_vcf)
Ar230_LM_maf1_genlight
indNames(Ar230_LM_maf1_genlight)
ploidy(Ar230_LM_maf1_genlight) <- 1
strata.df <- read.csv("230_strata.csv", head = FALSE, sep = ",")
strata(Ar230_LM_maf1_genlight) <- strata.df
splitStrata(Ar230_LM_maf1_genlight) <- ~Year/State/Location/Farm/Host/Zone/ICC3996_rating/Genesis090_rating/HatTrick_rating/Seamer_rating/Path1/Path2/Cluster/Group
Ar230_LM_maf1_genlight

# NJ tree
ind_dist <- bitwise.dist(Ar230_LM_maf1_genlight) 
# When differences_only = TRUE, the output will reflect the number of different loci. 
# The default setting, differences_only = FALSE, reflects the number of different alleles
# Note: this has no effect on haploid organisms since 1 locus = 1 allele
# Missing match determines whether two samples differing by missing data in a location should be counted as matching at that location. 
# Default set to TRUE, which forces missing data to match with anything. 
# FALSE forces missing data to not match with any other information, including other missing data
# percent = TRUE is default, that is, distance represented from 0 to 1
# FALSE will return the distance represented as integers from 1 to n where n is the number of loci.
# This option has no effect if euclidean = TRUE
ind_dist_no_of_loci <- bitwise.dist(Ar230_LM_maf1_genlight, percent = FALSE)
library(MASS)
write.matrix (ind_dist_no_of_loci, file = "ind_dist_no_of_loci")

Ar2021_genlight_tree <- aboot(Ar230_LM_maf1_genlight, tree = "nj", distance = bitwise.dist, sample = 1000, showtree = T, cutoff = 70, quiet = T, missing = ignore)
library(ape)
ape::write.tree(Ar2021_genlight_tree, file='individual_NJ_tree.txt')
# Colour by Zone
setPop(Ar230_LM_maf1_genlight) <- ~Zone
popNames(Ar230_LM_maf1_genlight)
cols <- c("Yellow2", "dark green", "green","gray", "turquoise", "sky blue", "magenta", "orange", "purple", "blue", "red")
plot.phylo(Ar2021_genlight_tree, type = "phylogram", cex = 0.9, font = 2, adj = 0, tip.color = cols[Ar230_LM_maf1_genlight$pop], label.offset = 0.0001, node.pos = NULL)
plot.phylo(Ar2021_genlight_tree, type="fan", cex = 0.5, font = 2, adj = 0, tip.color = cols[Ar230_LM_maf1_genlight$pop], label.offset = 0.0001, direcion="downwards", align.tip.label=FALSE)
nodelabels(Ar2021_genlight_tree$node.label, adj = c(1.3, -0.5), frame = "n", cex = 0.6, font = 0.5, xpd = TRUE)
axisPhylo(3)
legend('bottomright', legend = c("SA/Vic2",  "SA/Vic1", "NSW/Qld1", "unknown","NSW/Vic",  "WA2","Qld",  "SA1",  "WA1","NSW/Qld2", "NSW1"), fill = cols, border = FALSE, bty = "n", cex = 0.8)
# Colour by tate
setPop(Ar230_LM_maf1_genlight) <- ~State
popNames(Ar230_LM_maf1_genlight)
cols <- c("dark green", "red","Purple", "orange", "blue")
plot.phylo(Ar2021_genlight_tree, type = "phylogram", cex = 0.9, font = 2, adj = 0, tip.color = cols[Ar230_LM_maf1_genlight$pop], label.offset = 0.0001, node.pos = NULL)
plot.phylo(Ar2021_genlight_tree, type="fan", cex = 0.5, font = 2, adj = 0, tip.color = cols[Ar230_LM_maf1_genlight$pop], label.offset = 0.0001, direcion="downwards", align.tip.label=FALSE)
nodelabels(Ar2021_genlight_tree$node.label, adj = c(1.3, -0.5), frame = "n", cex = 0.6, font = 0.5, xpd = TRUE)
axisPhylo(3)
legend('bottomright', legend = c("VIC", "NSW", "WA", "QLD", "SA"), fill = cols, border = FALSE, bty = "n", cex = 0.8)
# Colour by Year
setPop(Ar230_LM_maf1_genlight) <- ~Year
popNames(Ar230_LM_maf1_genlight)
cols <- c("darkorange1", "purple", "light blue","cyan", "blue", "Yellow2")
plot.phylo(Ar2021_genlight_tree, type = "phylogram", cex = 0.9, font = 2, adj = 0, tip.color = cols[Ar230_LM_maf1_genlight$pop], label.offset = 0.0001, node.pos = NULL)
plot.phylo(Ar2021_genlight_tree, type="fan", cex = 0.5, font = 2, adj = 0, tip.color = cols[Ar230_LM_maf1_genlight$pop], label.offset = 0.0001, direcion="downwards", align.tip.label=FALSE)
nodelabels(Ar2021_genlight_tree$node.label, adj = c(1.3, -0.5), frame = "n", cex = 0.6, font = 0.5, xpd = TRUE)
axisPhylo(3)
# Colour by cluster
setPop(Ar230_LM_maf1_genlight) <- ~Cluster
popNames(Ar230_LM_maf1_genlight)
cols <- c("purple4", "darkgreen", "Darkorange3")
plot.phylo(Ar2021_genlight_tree, type = "phylogram", cex = 0.9, font = 2, adj = 0, tip.color = cols[Ar230_LM_maf1_genlight$pop], label.offset = 0.0001, node.pos = NULL)
plot.phylo(Ar2021_genlight_tree, type="fan", cex = 0.5, font = 2, adj = 0, tip.color = cols[Ar230_LM_maf1_genlight$pop], label.offset = 0.0001, direcion="downwards", align.tip.label=FALSE)
nodelabels(Ar2021_genlight_tree$node.label, adj = c(1.3, -0.5), frame = "n", cex = 0.6, font = 0.5, xpd = TRUE)
axisPhylo(3)
legend('bottomright', legend = c("3", "1", "2"), fill = cols, border = FALSE, bty = "n", cex = 0.8)
# Colour by Pathogenecity_1 data
setPop(Ar230_LM_maf1_genlight) <- ~Path1
popNames(Ar230_LM_maf1_genlight)
cols <- c("yellowgreen", "yellow", "orange3", "green", "orange", "red" )
plot.phylo(Ar2021_genlight_tree, type = "phylogram", cex = 0.9, font = 2, adj = 0, tip.color = cols[Ar230_LM_maf1_genlight$pop], label.offset = 0.0001, node.pos = NULL)
plot.phylo(Ar2021_genlight_tree, type="fan", cex = 0.5, font = 2, adj = 0, tip.color = cols[Ar230_LM_maf1_genlight$pop], label.offset = 0.0001, direcion="downwards", align.tip.label=FALSE)
nodelabels(Ar2021_genlight_tree$node.label, adj = c(1.3, -0.5), frame = "n", cex = 0.6, font = 0.5, xpd = TRUE)
axisPhylo(3)
legend('bottomright', legend = c("1", "2", "4", "0", "3", "5"), fill = cols, border = FALSE, bty = "n", cex = 0.8)
# Colour by ICC3996_rating
setPop(Ar230_LM_maf1_genlight) <- ~ICC3996_rating
popNames(Ar230_LM_maf1_genlight)
cols <- c("darkgreen", "yellow", "red", "gray" )
plot.phylo(Ar2021_genlight_tree, type = "phylogram", cex = 0.9, font = 2, adj = 0, tip.color = cols[Ar230_LM_maf1_genlight$pop], label.offset = 0.0001, node.pos = NULL)
plot.phylo(Ar2021_genlight_tree, type="fan", cex = 0.5, font = 2, adj = 0, tip.color = cols[Ar230_LM_maf1_genlight$pop], label.offset = 0.0001, direcion="downwards", align.tip.label=FALSE)
nodelabels(Ar2021_genlight_tree$node.label, adj = c(1.3, -0.5), frame = "n", cex = 0.6, font = 0.5, xpd = TRUE)
axisPhylo(3)
legend('bottomright', legend = c("ICC3996 low", "ICC3996 moderate", "ICC3996 high", "unknown"), fill = cols, border = FALSE, bty = "n", cex = 0.8)
# Colour by Seamer_rating
setPop(Ar230_LM_maf1_genlight) <- ~Seamer_rating
popNames(Ar230_LM_maf1_genlight)
cols <- c("black", "darkgreen", "yellow", "red" )
plot.phylo(Ar2021_genlight_tree, type = "phylogram", cex = 0.9, font = 2, adj = 0, tip.color = cols[Ar230_LM_maf1_genlight$pop], label.offset = 0.0001, node.pos = NULL)
plot.phylo(Ar2021_genlight_tree, type="fan", cex = 0.5, font = 2, adj = 0, tip.color = cols[Ar230_LM_maf1_genlight$pop], label.offset = 0.0001, direcion="downwards", align.tip.label=FALSE)
nodelabels(Ar2021_genlight_tree$node.label, adj = c(1.3, -0.5), frame = "n", cex = 0.6, font = 0.5, xpd = TRUE)
axisPhylo(3)
legend('bottomright', legend = c("unknown", "Seamer low", "Seamer moderate", "Seamer high"), fill = cols, border = FALSE, bty = "n", cex = 0.8)
# Colour by Genesis090_rating
setPop(Ar230_LM_maf1_genlight) <- ~Genesis090_rating
popNames(Ar230_LM_maf1_genlight)
cols <- c("darkgreen", "yellow", "red", "gray" )
plot.phylo(Ar2021_genlight_tree, type = "phylogram", cex = 0.9, font = 2, adj = 0, tip.color = cols[Ar230_LM_maf1_genlight$pop], label.offset = 0.0001, node.pos = NULL)
plot.phylo(Ar2021_genlight_tree, type="fan", cex = 0.5, font = 2, adj = 0, tip.color = cols[Ar230_LM_maf1_genlight$pop], label.offset = 0.0001, direcion="downwards", align.tip.label=FALSE)
nodelabels(Ar2021_genlight_tree$node.label, adj = c(1.3, -0.5), frame = "n", cex = 0.6, font = 0.5, xpd = TRUE)
axisPhylo(3)
legend('bottomright', legend = c("Genesis090 low", "Genesis090 moderate", "Genesis090 high", "unknown"), fill = cols, border = FALSE, bty = "n", cex = 0.8)
# Colour by HatTrick_rating
setPop(Ar230_LM_maf1_genlight) <- ~HatTrick_rating
popNames(Ar230_LM_maf1_genlight)
cols <- c("red", "darkgreen", "yellow", "gray")
plot.phylo(Ar2021_genlight_tree, type = "phylogram", cex = 0.9, font = 2, adj = 0, tip.color = cols[Ar230_LM_maf1_genlight$pop], label.offset = 0.0001, node.pos = NULL)
plot.phylo(Ar2021_genlight_tree, type="fan", cex = 0.5, font = 2, adj = 0, tip.color = cols[Ar230_LM_maf1_genlight$pop], label.offset = 0.0001, direcion="downwards", align.tip.label=FALSE)
nodelabels(Ar2021_genlight_tree$node.label, adj = c(1.3, -0.5), frame = "n", cex = 0.6, font = 0.5, xpd = TRUE)
axisPhylo(3)
legend('bottomright', legend = c("HatTrick high", "HatTrick low", "HatTrick moderate", "unknown"), fill = cols, border = FALSE, bty = "n", cex = 0.8)

##  DISTANCE TREE GENLIGHT OBJECT for ALL loci
Ar230_all_maf1_vcf <- read.vcfR("All_230_sorted.vcf")
Ar230_all_maf1_genlight <- vcfR2genlight(Ar230_all_maf1_vcf)
indNames(Ar230_all_maf1_genlight)
ploidy(Ar230_all_maf1_genlight) <- 1
strata.df <- read.csv("230_strata.csv", head = FALSE, sep = ",")
strata(Ar230_all_maf1_genlight) <- strata.df
splitStrata(Ar230_all_maf1_genlight) <- ~Year/State/Location/Farm/Host/Zone/ICC3996_rating/Genesis090_rating/HatTrick_rating/Seamer_rating/Path1/Path2/Cluster/Group
Ar230_all_maf1_genlight

# NJ tree
Ar2021_genlight_tree_all <- aboot(Ar230_all_maf1_genlight, tree = "nj", distance = bitwise.dist, sample = 1000, showtree = T, cutoff = 70, quiet = T, missing=ignore)
library(ape)
ape::write.tree(Ar2021_genlight_tree_all, file='individual_NJ_tree_all.txt')
# Colour by pathogenecity_1 scores
setPop(Ar230_all_maf1_genlight) <- ~Path1
popNames(Ar230_all_maf1_genlight)
cols <- c("yellowgreen", "yellow", "pink3", "darkgreen", "orange", "red3", "black")
plot.phylo(Ar2021_genlight_tree_all, type = "phylogram", cex = 0.9, font = 2, adj = 0, tip.color = cols[Ar230_all_maf1_genlight$pop], label.offset = 0.0001, node.pos = NULL)
plot.phylo(Ar2021_genlight_tree_all, type="fan", cex = 0.5, font = 2, adj = 0, tip.color = cols[Ar230_all_maf1_genlight$pop], label.offset = 0.0001, direcion="downwards", align.tip.label=FALSE)
nodelabels(Ar2021_genlight_tree_all$node.label, adj = c(1.3, -0.5), frame = "n", cex = 0.6, font = 0.5, xpd = TRUE)
axisPhylo(3)
legend('bottomright', legend = c("1", "2", "4", "0", "3", "5", "unknown"), fill = cols, border = FALSE, bty = "n", cex = 0.8)
# Colour by Cluster
setPop(Ar230_all_maf1_genlight) <- ~Cluster
popNames(Ar230_all_maf1_genlight)
cols <- c("purple4", "darkgreen", "Darkorange3")
plot.phylo(Ar2021_genlight_tree_all, type = "phylogram", cex = 0.9, font = 2, adj = 0, tip.color = cols[Ar230_all_maf1_genlight$pop], label.offset = 0.0001, node.pos = NULL)
plot.phylo(Ar2021_genlight_tree_all, type="fan", cex = 0.5, font = 2, adj = 0, tip.color = cols[Ar230_all_maf1_genlight$pop], label.offset = 0.0001, direcion="downwards", align.tip.label=FALSE)
nodelabels(Ar2021_genlight_tree_all$node.label, adj = c(1.3, -0.5), frame = "n", cex = 0.6, font = 0.5, xpd = TRUE)
axisPhylo(3)
legend('bottomright', legend = c("3", "1", "2"), fill = cols, border = FALSE, bty = "n", cex = 0.8)

####################################
##  INDICES OF GENETIC DIVERSITY ###
####################################

setPop(Ar230_LM_maf1) <- ~Zone
popNames(Ar230_LM_maf1)
(Ar2021_diversity <- poppr(Ar230_LM_maf1, sample = 1000))
(N <- Ar2021_diversity$N)
(lambda <- Ar2021_diversity$lambda) 
#corrected simpson's index for sample size
(N/(N-1))*lambda
(MLG <- Ar2021_diversity$MLG)
(Clonal_Fraction <- (1-(MLG/N)))

setPop(Ar230_LM_maf1) <- ~State
popNames(Ar230_LM_maf1)
(Ar2021_diversity <- poppr(Ar230_LM_maf1, sample = 1000))
(N <- Ar2021_diversity$N)
(lambda <- Ar2021_diversity$lambda) 
#corrected simpson's index for sample size
(N/(N-1))*lambda
(MLG <- Ar2021_diversity$MLG)
(Clonal_Fraction <- (1-(MLG/N)))

#####################################################
##  DISCRIMINANT ANALYSIS OF PRINCIPAL COMPONENTS ###
#####################################################

# DAPC with populations pre-defined based on year of collection
setPop(Ar230_LM_maf1) <- ~Year
popNames(Ar230_LM_maf1)
cols3 <- c("darkorange1", "purple", "light blue","cyan", "blue", "Yellow2")
# Cross-validation (xvalDapc) is an objective way to identify no. of PCs to retain.
set.seed(999)
xvalAr2021 <- xvalDapc(tab(Ar230_LM_maf1, NA.method = "mean"), pop(Ar230_LM_maf1), training.set = 0.7)
set.seed(999)
xvalAr2021 <- xvalDapc(tab(Ar230_LM_maf1, NA.method = "mean"), pop(Ar230_LM_maf1), parallel = "multicore", ncpus = 4L, training.set = 0.9, n.rep = 1000, n.pca = 18:40)
xvalAr2021 [2:6]
#32
popNames(Ar230_LM_maf1)
dapc_year <- dapc(Ar230_LM_maf1, var.contrib = TRUE, scale = FALSE, n.pca = 32, n.da = nPop(Ar230_LM_maf1) - 1)
# The ellipses represent the maximum area spanned by 95% of the data in a population by year of collection
scatter(dapc_year, col = cols3, clabel=FALSE, xax = 1, yax = 2,inset.solid = 1, pch=20, solid = 0.6, cstar=0, cellipse = 2.5, legend = TRUE, cleg = 0.75, posi.leg = "bottomright", cex = 1.5, scree.da = TRUE, scree.pca = TRUE, posi.pca = "topleft", ratio.da = 0.15, ratio.pca = 0.15, inset.da = 0.03, inset.pca = 0.03, posi.da = "bottomleft")

# DAPC with populations pre-defined based on AE Zone
setPop(Ar230_LM_maf1) <- ~Zone
popNames(Ar230_LM_maf1)
# Removing isolates with unknown Zone
Ar230_LM_maf1_zone <- popsub(Ar230_LM_maf1, c(1,2,3,5,6,7,8,9,10,11))
popNames(Ar230_LM_maf1_zone)
set.seed(999)
xvalAr2021 <- xvalDapc(tab(Ar230_LM_maf1_zone, NA.method = "mean"), pop(Ar230_LM_maf1_zone), training.set = 0.7)
set.seed(999)
xvalAr2021 <- xvalDapc(tab(Ar230_LM_maf1_zone, NA.method = "mean"), pop(Ar230_LM_maf1_zone), parallel = "multicore", ncpus = 4L, training.set = 0.9, n.rep = 1000, n.pca = 35:70)
xvalAr2021 [2:6]
#57
popNames(Ar230_LM_maf1_zone)
cols <- c("Yellow2", "dark green", "green", "turquoise", "sky blue", "magenta", "orange", "purple", "blue", "red")
dapc.zone <- dapc(Ar230_LM_maf1_zone, var.contrib = TRUE, scale = FALSE, n.pca = 58, n.da = nPop(Ar230_LM_maf1_zone) - 1)
scatter(dapc.zone, col = cols, clabel=FALSE, xax = 1, yax = 2,inset.solid = 1, cstar=0, solid = 0.6, cellipse = 2.5, legend = TRUE, cleg = 0.75, posi.leg = "bottomright", cex = 1.5, scree.da = TRUE, scree.pca = TRUE, posi.pca = "topleft", ratio.da = 0.15, ratio.pca = 0.15, inset.da = 0.03, inset.pca = 0.03, posi.da = "bottomleft")

# DAPC with populations pre-defined based on chickpea cultivar
setPop(Ar230_LM_maf1) <- ~Host
popNames(Ar230_LM_maf1)
# Removing isolates with unknown Zone
Ar230_LM_maf1_Host <- popsub(Ar230_LM_maf1, c(1,2,3,5,6,7,8,9,10,11, 12, 13, 14, 16, 17, 18, 19, 20, 21, 22, 23, 24, 25, 26, 27, 28))
popNames(Ar230_LM_maf1_Host)
set.seed(999)
xvalAr2021 <- xvalDapc(tab(Ar230_LM_maf1_Host, NA.method = "mean"), pop(Ar230_LM_maf1_Host), training.set = 0.7)
set.seed(999)
xvalAr2021 <- xvalDapc(tab(Ar230_LM_maf1_Host, NA.method = "mean"), pop(Ar230_LM_maf1_Host), parallel = "multicore", ncpus = 4L, training.set = 0.9, n.rep = 1000, n.pca = 15:51)
xvalAr2021 [2:6]
#57
popNames(Ar230_LM_maf1_Host)
cols <- rainbow(27)
dapc.zone <- dapc(Ar230_LM_maf1_Host, var.contrib = TRUE, scale = FALSE, n.pca = 48, n.da = nPop(Ar230_LM_maf1_Host) - 1)
scatter(dapc.zone, col = cols, clabel=FALSE, xax = 1, yax = 2,inset.solid = 1, cstar=0, solid = 0.6, cellipse = 2.5, legend = TRUE, cleg = 0.75, posi.leg = "bottomright", cex = 1.5, scree.da = TRUE, scree.pca = TRUE, posi.pca = "topleft", ratio.da = 0.15, ratio.pca = 0.15, inset.da = 0.03, inset.pca = 0.03, posi.da = "bottomleft")

#######################################################
### FIND CLUSTERS - underlying population structure ###
#######################################################

library(adegenet)
library(vcfR)
Ar230_LM_maf1_genlight
indNames(Ar230_LM_maf1_genlight)
Ar230_LM_maf1_genlight.clusters <- find.clusters(Ar230_LM_maf1_genlight, max.n.clust = 10, n.start =1000)
#250
#3
dapcAr2021_LM_maf1 <- dapc(Ar230_LM_maf1_genlight, Ar230_LM_maf1_genlight.clusters$grp, var.contrib = TRUE)
# Choose the number PCs to retain (>=1): 
#50
#2
scatter(dapcAr2021_LM_maf1, legend = TRUE, cleg = 1, col = cols, clabel = 0, cex = 1.5, scree.da = TRUE, scree.pca = TRUE, posi.pca = "topleft", posi.da = "bottomleft", ratio.da = 0.15, ratio.pca = 0.15, inset.da = 0.01, inset.pca = 0.01, posi.leg = "bottomright")
# Check which isolates belong to each of the three clusters
dapcAr2021_LM_maf1
dapcAr2021_LM_maf1$grp

###################################
### ALL LOCI as GenClone object ###
###################################

## GenClone object produced previously
Ar230_maf1
## DELINEATE CLONAL LINEAGES
# to predict a cut off value
Ar230_maf1_thresholds <- mlg.filter(Ar230_maf1, distance = bitwise.dist, stats = "THRESHOLDS", threshold = 1)
pthresh  <- filter_stats(Ar230_maf1, distance=bitwise.dist, plot = TRUE, stats = "THRESHOLD", threads = 1L)
cutoff_predictor(pthresh$farthest, fraction = 0.5)
mlg.filter(Ar230_maf1, distance = bitwise.dist) <- 0.002284496
Ar230_maf1
indNames(Ar230_maf1)

##########################
### SUB POPULATIONS    ###
##########################

setPop(Ar230_maf1) <- ~Cluster
popNames(Ar230_maf1)
Ar230_maf1_1 <- popsub(Ar230_maf1, sublist = c(2))
Ar230_maf1_1

Ar230_maf1_2 <- popsub(Ar230_maf1, sublist = c(3))
Ar230_maf1_2

Ar230_maf1_3 <- popsub(Ar230_maf1, sublist = c(1))
Ar230_maf1_3

setPop(Ar230_maf1) <- ~Path1
popNames(Ar230_maf1)
known_path_maf1 <- popsub(Ar230_maf1, sublist = c(1,2,3,4,5,6))
known_path_maf1

#######################################################################
###   Discriminant Analysis of Principal components    Cluster 1    ###
#######################################################################

setPop(Ar230_maf1_1) <- ~ICC3996_rating
popNames(Ar230_maf1_1)
xvalAr2021 <- xvalDapc(tab(Ar230_maf1_1, NA.method = "mean"), pop(Ar230_maf1_1), training.set = 0.7)
xvalAr2021 <- xvalDapc(tab(Ar230_maf1_1, NA.method = "mean"), pop(Ar230_maf1_1), parallel = "multicore", ncpus = 4L, training.set = 0.9, n.rep = 1000, n.pca = 10:25)
xvalAr2021 [2:6]
# Check Number of PCs Achieving Lowest MS or Number of PCs Achieving Highest Mean Success
# Use the lowest RMSE as the 'optimum' n.pca in the DAPC analysis.
# no. of PCs retained for ICC3996_rating: 13
dapc.ICC <- dapc(Ar230_maf1_1, var.contrib = TRUE, scale = FALSE, n.pca = 13, n.da = nPop(Ar230_maf1_1) - 1)
popNames(Ar230_maf1_1)
cols <- c("darkgreen", "yellow2","Red3")
scatter(dapc.ICC, col = cols, clabel=FALSE, legend = TRUE, cleg = 0.7, cstar=1, solid = 0.6, cellipse = 2.5, posi.leg = "bottomright", cex = 1.5, scree.da = FALSE, scree.pca = FALSE, posi.pca = "topleft", ratio.pca = 0.32, inset.pca = 0.0)
contrib_ICC <- loadingplot(dapc.ICC$var.contr, axis = 1, thres = 0.005)
contrib_ICC

setPop(Ar230_maf1_1) <- ~Genesis090_rating
popNames(Ar230_maf1_1)
xvalAr2021 <- xvalDapc(tab(Ar230_maf1_1, NA.method = "mean"), pop(Ar230_maf1_1), training.set = 0.7)
xvalAr2021 <- xvalDapc(tab(Ar230_maf1_1, NA.method = "mean"), pop(Ar230_maf1_1), parallel = "multicore", ncpus = 4L, training.set = 0.9, n.rep = 1000, n.pca = 10:27)
xvalAr2021 [2:6]
# no. of PCs retained for Genesis090_rating: 25
dapc.Gen <- dapc(Ar230_maf1_1, var.contrib = TRUE, scale = FALSE, n.pca = 25, n.da = nPop(Ar230_maf1_1) - 1)
popNames(Ar230_maf1_1)
cols <- c("yellow2","Red3", "darkgreen")
scatter(dapc.Gen, col = cols, clabel=FALSE, legend = TRUE, cleg = 0.7, posi.leg = "bottomright",  cstar=1, solid = 0.6, cellipse = 2.5, cex = 1.5, scree.da = FALSE, scree.pca = FALSE, posi.pca = "topright", ratio.pca = 0.32, inset.pca = 0.0)
contrib_Gen <- loadingplot(dapc.Gen$var.contr, axis = 1, thres = 0.005)
contrib_Gen

setPop(Ar230_maf1_1) <- ~Seamer_rating
popNames(Ar230_maf1_1)
Ar_Seamer_maf1_1 <- popsub(Ar230_maf1_1, sublist = c(2,3,4))
Ar_Seamer_maf1_1
xvalAr2021 <- xvalDapc(tab(Ar_Seamer_maf1_1, NA.method = "mean"), pop(Ar_Seamer_maf1_1), training.set = 0.7)
xvalAr2021 <- xvalDapc(tab(Ar_Seamer_maf1_1, NA.method = "mean"), pop(Ar_Seamer_maf1_1), parallel = "multicore", ncpus = 4L, training.set = 0.9, n.rep = 1000, n.pca = 4:24)
xvalAr2021 [2:6]
# no. of PCs retained for Genesis090_rating: 4
dapc.Sea <- dapc(Ar_Seamer_maf1_1, var.contrib = TRUE, scale = FALSE, n.pca = 4, n.da = nPop(Ar_Seamer_maf1_1) - 1)
popNames(Ar_Seamer_maf1_1)
cols <- c("darkgreen","yellow2","Red3")
scatter(dapc.Sea, col = cols, clabel=FALSE, legend = TRUE, cleg = 0.7, posi.leg = "bottomleft",  cstar=1, solid = 0.6, cellipse = 2.5, cex = 1.5, scree.da = FALSE, scree.pca = FALSE, posi.pca = "topright", ratio.pca = 0.32, inset.pca = 0.0)
contrib_Sea <- loadingplot(dapc.Sea$var.contr, axis = 1, thres = 0.005)
contrib_Sea

setPop(Ar230_maf1_1) <- ~HatTrick_rating
popNames(Ar230_maf1_1)
Ar_Hattrick_maf1_1 <- popsub(Ar230_maf1_1, sublist = c(1,2,3))
Ar_Hattrick_maf1_1
xvalAr2021 <- xvalDapc(tab(Ar_Hattrick_maf1_1, NA.method = "mean"), pop(Ar_Hattrick_maf1_1), training.set = 0.7)
xvalAr2021 <- xvalDapc(tab(Ar_Hattrick_maf1_1, NA.method = "mean"), pop(Ar_Hattrick_maf1_1), parallel = "multicore", ncpus = 4L, training.set = 0.9, n.rep = 1000, n.pca = 14:20)
xvalAr2021 [2:6]
# no. of PCs retained for Genesis090_rating: 14
dapc.Hat <- dapc(Ar_Hattrick_maf1_1, var.contrib = TRUE, scale = FALSE, n.pca = 14, n.da = nPop(Ar_Hattrick_maf1_1) - 1)
popNames(Ar_Hattrick_maf1_1)
cols <- c("Red3", "darkgreen","yellow2")
scatter(dapc.Hat, col = cols, clabel=FALSE, legend = TRUE, cleg = 0.7, posi.leg = "topleft", cstar=1, solid = 0.6, cellipse = 2.5, cex = 1.5, scree.da = FALSE, scree.pca = FALSE, posi.pca = "topright", ratio.pca = 0.32, inset.pca = 0.0)
contrib_Hat <- loadingplot(dapc.Hat$var.contr, axis = 1, thres = 0.006)
contrib_Hat

#######################################################################
###   Discriminant Analysis of Principal components    Cluster 2    ###
#######################################################################

setPop(Ar230_maf1_2) <- ~ICC3996_rating
popNames(Ar230_maf1_2)
Ar_ICC_maf1_2 <- popsub(Ar230_maf1_2, sublist = c(1,2,3))
xvalAr2021 <- xvalDapc(tab(Ar_ICC_maf1_2, NA.method = "mean"), pop(Ar_ICC_maf1_2), training.set = 0.7)
xvalAr2021 <- xvalDapc(tab(Ar_ICC_maf1_2, NA.method = "mean"), pop(Ar_ICC_maf1_2), parallel = "multicore", ncpus = 4L, training.set = 0.9, n.rep = 1000, n.pca = 14:36)
xvalAr2021 [2:6]
# no. of PCs retained for ICC3996_rating: 34
dapc2.ICC <- dapc(Ar_ICC_maf1_2, var.contrib = TRUE, scale = FALSE, n.pca = 34, n.da = nPop(Ar_ICC_maf1_2) - 1)
popNames(Ar_ICC_maf1_2)
cols <- c("yellow2","darkgreen","Red3")
scatter(dapc2.ICC, col = cols, clabel=FALSE, legend = TRUE, cleg = 0.7, posi.leg = "bottomright", cex = 1.5, cstar=1, solid = 0.6, cellipse = 2.5, scree.da = FALSE, scree.pca = FALSE, posi.pca = "topright", ratio.pca = 0.32, inset.pca = 0.0)
contrib_ICC <- loadingplot(dapc2.ICC$var.contr, axis = 1, thres = 0.005)
contrib_ICC

setPop(Ar230_maf1_2) <- ~Genesis090_rating
popNames(Ar230_maf1_2)
Ar_Gen_maf1_2 <- popsub(Ar230_maf1_2, sublist = c(1,2,3))
xvalAr2021 <- xvalDapc(tab(Ar_Gen_maf1_2, NA.method = "mean"), pop(Ar_Gen_maf1_2), training.set = 0.7)
xvalAr2021 <- xvalDapc(tab(Ar_Gen_maf1_2, NA.method = "mean"), pop(Ar_Gen_maf1_2), parallel = "multicore", ncpus = 4L, training.set = 0.9, n.rep = 1000, n.pca = 14:36)
xvalAr2021 [2:6]
# no. of PCs retained for Genesis090_rating: 26
dapc2.Gen <- dapc(Ar_Gen_maf1_2, var.contrib = TRUE, scale = FALSE, n.pca = 26, n.da = nPop(Ar_Gen_maf1_2) - 1)
popNames(Ar_Gen_maf1_2)
cols <- c("yellow2", "darkgreen","Red3")
scatter(dapc2.Gen, col = cols, clabel=FALSE, legend = TRUE, cleg = 0.7, posi.leg = "bottomright", cex = 1.5, scree.da = FALSE, scree.pca = FALSE,cstar=1, solid = 0.6, cellipse = 2.5, posi.pca = "topleft", ratio.pca = 0.32, inset.pca = 0.0)
contrib_Gen <- loadingplot(dapc2.Gen$var.contr, axis = 1, thres = 0.005)
contrib_Gen

setPop(Ar230_maf1_2) <- ~Seamer_rating
popNames(Ar230_maf1_2)
Ar_Seamer_maf1_2 <- popsub(Ar230_maf1_2, sublist = c(2,3,4))
Ar_Seamer_maf1_2
xvalAr2021 <- xvalDapc(tab(Ar_Seamer_maf1_2, NA.method = "mean"), pop(Ar_Seamer_maf1_2), training.set = 0.7)
xvalAr2021 <- xvalDapc(tab(Ar_Seamer_maf1_2, NA.method = "mean"), pop(Ar_Seamer_maf1_2), parallel = "multicore", ncpus = 4L, training.set = 0.9, n.rep = 1000, n.pca = 1:40)
xvalAr2021 [2:6]
# no. of PCs retained for Genesis090_rating: 6
dapc2.Sea <- dapc(Ar_Seamer_maf1_2, var.contrib = TRUE, scale = FALSE, n.pca =6, n.da = nPop(Ar_Seamer_maf1_2) - 1)
popNames(Ar_Seamer_maf1_2)
cols <- c("darkgreen","yellow2","Red3")
scatter(dapc2.Sea, col = cols, clabel=FALSE, legend = TRUE, cleg = 0.7, posi.leg = "bottomleft", cex = 1.5, scree.da = FALSE, scree.pca = TRUE, cstar=1, solid = 0.6, cellipse = 2.5, posi.pca = "topleft", ratio.pca = 0.32, inset.pca = 0.0)
contrib_Sea <- loadingplot(dapc2.Sea$var.contr, axis = 1, thres = 0.005)
contrib_Sea

Ar230_maf1_2
setPop(Ar230_maf1_2) <- ~HatTrick_rating
popNames(Ar230_maf1_2)
Ar_Hattrick_maf1_2 <- popsub(Ar230_maf1_2, sublist = c(1,2,3))
Ar_Hattrick_maf1_2
xvalAr2021 <- xvalDapc(tab(Ar_Hattrick_maf1_2, NA.method = "mean"), pop(Ar_Hattrick_maf1_2), training.set = 0.7)
xvalAr2021 <- xvalDapc(tab(Ar_Hattrick_maf1_2, NA.method = "mean"), pop(Ar_Hattrick_maf1_2), parallel = "multicore", ncpus = 4L, training.set = 0.9, n.rep = 1000, n.pca = 5:28)
xvalAr2021 [2:6]
# no. of PCs retained for Genesis090_rating: 5
dapc2.Hat <- dapc(Ar_Hattrick_maf1_2, var.contrib = TRUE, scale = FALSE, n.pca = 5, n.da = nPop(Ar_Hattrick_maf1_2) - 1)
popNames(Ar_Hattrick_maf1_2)
cols <- c("Red3", "yellow2","darkgreen")
scatter(dapc2.Hat, col = cols, clabel=FALSE, legend = TRUE, cleg = 0.7, posi.leg = "bottomleft", cex = 1.5, scree.da = FALSE, scree.pca = FALSE, posi.pca = "topleft",cstar=1, solid = 0.6, cellipse = 2.5, ratio.pca = 0.32, inset.pca = 0.0)
contrib_Hat <- loadingplot(dapc2.Hat$var.contr, axis = 1, thres = 0.005)
contrib_Hat

#######################################################################
###   Discriminant Analysis of Principal components    Cluster 3    ###
#######################################################################

setPop(Ar230_maf1_3) <- ~ICC3996_rating
popNames(Ar230_maf1_3)
Ar_ICC_maf1_3 <- popsub(Ar230_maf1_3, sublist = c(1,2,3))
xvalAr2021 <- xvalDapc(tab(Ar_ICC_maf1_3, NA.method = "mean"), pop(Ar_ICC_maf1_3), training.set = 0.7)
xvalAr2021 <- xvalDapc(tab(Ar_ICC_maf1_3, NA.method = "mean"), pop(Ar_ICC_maf1_3), parallel = "multicore", ncpus = 4L, training.set = 0.9, n.rep = 1000, n.pca = 25:32)
xvalAr2021 [2:6]
# no. of PCs retained for ICC3996_rating: 26
dapc3.ICC <- dapc(Ar_ICC_maf1_3, var.contrib = TRUE, scale = FALSE, n.pca = 26, n.da = nPop(Ar_ICC_maf1_3) - 1)
popNames(Ar_ICC_maf1_3)
cols <- c("darkgreen","yellow2","Red3")
scatter(dapc3.ICC, col = cols, clabel=FALSE, legend = TRUE, cleg = 0.7, posi.leg = "topright", cstar=1, solid = 0.6, cellipse = 2.5, cex = 1.5, scree.da = FALSE, scree.pca = FALSE, posi.pca = "topleft", ratio.pca = 0.32, inset.pca = 0.0)
contrib_ICC <- loadingplot(dapc3.ICC$var.contr, axis = 1, thres = 0.005)
contrib_ICC

setPop(Ar230_maf1_3) <- ~Genesis090_rating
popNames(Ar230_maf1_3)
Ar_Gen_maf1_3 <- popsub(Ar230_maf1_3, sublist = c(1,2,3))
xvalAr2021 <- xvalDapc(tab(Ar_Gen_maf1_3, NA.method = "mean"), pop(Ar_Gen_maf1_3), training.set = 0.7)
xvalAr2021 <- xvalDapc(tab(Ar_Gen_maf1_3, NA.method = "mean"), pop(Ar_Gen_maf1_3), parallel = "multicore", ncpus = 4L, training.set = 0.9, n.rep = 1000, n.pca = 29:33)
xvalAr2021 [2:6]
# no. of PCs retained for Genesis090_rating: 31
dapc3.Gen <- dapc(Ar_Gen_maf1_3, var.contrib = TRUE, scale = FALSE, n.pca = 31, n.da = nPop(Ar_Gen_maf1_3) - 1)
popNames(Ar_Gen_maf1_3)
cols <- c("darkgreen", "Red3", "yellow2")
scatter(dapc3.Gen, col = cols, clabel=FALSE, legend = TRUE, cleg = 0.7, posi.leg = "topright", cex = 1.5,cstar=1, solid = 0.6, cellipse = 2.5, scree.da = FALSE, scree.pca = FALSE, posi.pca = "topright", ratio.pca = 0.32, inset.pca = 0.0)
ontrib_Gen <- loadingplot(dapc3.Gen$var.contr, axis = 1, thres = 0.005)
contrib_Gen

setPop(Ar230_maf1_3) <- ~Seamer_rating
popNames(Ar230_maf1_3)
Ar_Seamer_maf1_3 <- popsub(Ar230_maf1_3, sublist = c(2,3,4))
Ar_Seamer_maf1_3
xvalAr2021 <- xvalDapc(tab(Ar_Seamer_maf1_3, NA.method = "mean"), pop(Ar_Seamer_maf1_3), training.set = 0.7)
xvalAr2021 <- xvalDapc(tab(Ar_Seamer_maf1_3, NA.method = "mean"), pop(Ar_Seamer_maf1_3), parallel = "multicore", ncpus = 4L, training.set = 0.9, n.rep = 1000, n.pca = 27:33)
xvalAr2021 [2:6]
# no. of PCs retained for Genesis090_rating: 27
dapc3.Sea <- dapc(Ar_Seamer_maf1_3, var.contrib = TRUE, scale = FALSE, n.pca = 27 , n.da = nPop(Ar_Seamer_maf1_3) - 1)
popNames(Ar_Seamer_maf1_3)
cols <- c("yellow2", "darkgreen", "Red3")
scatter(dapc3.Sea, col = cols, clabel=FALSE, legend = TRUE, cleg = 0.7, posi.leg = "bottomleft", cex = 1.5, cstar=1, solid = 0.6, cellipse = 2.5, scree.da = FALSE, scree.pca = FALSE, posi.pca = "topleft", ratio.pca = 0.32, inset.pca = 0.0)
contrib_Sea <- loadingplot(dapc3.Sea$var.contr, axis = 1, thres = 0.005)
contrib_Sea

setPop(Ar230_maf1_3) <- ~HatTrick_rating
popNames(Ar230_maf1_3)
Ar_Hattrick_maf1_3 <- popsub(Ar230_maf1_3, sublist = c(1,2,3))
xvalAr2021 <- xvalDapc(tab(Ar_Hattrick_maf1_3, NA.method = "mean"), pop(Ar_Hattrick_maf1_3), training.set = 0.7)
xvalAr2021 <- xvalDapc(tab(Ar_Hattrick_maf1_3, NA.method = "mean"), pop(Ar_Hattrick_maf1_3), parallel = "multicore", ncpus = 4L, training.set = 0.9, n.rep = 1000, n.pca = 20:44)
xvalAr2021 [2:6]
# no. of PCs retained for Genesis090_rating: 23
dapc3.Hat <- dapc(Ar_Hattrick_maf1_3, var.contrib = TRUE, scale = FALSE, n.pca = 23, n.da = nPop(Ar_Hattrick_maf1_3) - 1)
popNames(Ar_Hattrick_maf1_3)
cols <- c("Red3", "yellow2","darkgreen")
scatter(dapc3.Hat, col = cols, clabel=FALSE, legend = TRUE, cleg = 0.7, posi.leg = "bottomleft", cstar=1, solid = 0.6, cellipse = 2.5, cex = 1.5, scree.da = FALSE, scree.pca = FALSE, posi.pca = "topleft", ratio.pca = 0.32, inset.pca = 0.0)
contrib_Hat <- loadingplot(dapc3.Hat$var.contr, axis = 1, thres = 0.005)
contrib_Hat
```
